# *β*-actin is essential for structural integrity and physiological function of the retina

**DOI:** 10.1101/2023.03.27.534392

**Authors:** Pavan Vedula, Marie E. Fina, Brent A. Bell, Sergei S. Nikonov, Anna Kashina, Dawei W. Dong

**Affiliations:** Department of Biomedical Sciences, School of Veterinary Medicines; Department of Neuroscience, Perelman School of Medicine; Ophthalmology Department, Perelman School of Medicine; Institute for Biomedical Informatics, Perelman School of Medicine; University of Pennsylvania, Philadelphia, PA 19104

## Abstract

Lack of non-muscle *β*-actin gene (Actb) leads to early embryonic lethality in mice, however mice with *β*- to *γ*-actin replacement develop normally and show no detectable phenotypes at young age. Here we investigated the effect of this replacement in the retina. During aging, these mice have accelerated de-generation of retinal structure and function, including elongated microvilli and defective mitochondria of retinal pigment epithelium (RPE), abnormally bulging photoreceptor outer segments (OS) accompanied by reduced transducin concentration and light sensitivity, and accumulation of autofluorescent microglia cells in the subretinal space between RPE and OS. These defects are accompanied by changes in the F-actin binding of several key actin interacting partners, including ezrin, myosin, talin, and vinculin known to play central roles in modulating actin cytoskeleton and cell adhesion and mediating the phagocytosis of OS. Our data show that *β*-actin protein is essential for maintaining normal retinal structure and function.

## Introduction

Actins are abundant and ubiquitously expressed proteins that are essential components of the cytoskeleton in both muscle and non-muscle cells. Higher vertebrates express two non-muscle actin isoforms: *β*- and *γ*-cytoplasmic actin (referred to as *β*- and *γ*-actin hereon). These two proteins differ by only 4 conservative substitutions in their N-terminus, and are ubiquitously expressed in every mammalian cell. Lack of *β*-actin gene (Actb) leads to early embryonic lethality in mice [1], while *γ*-actin gene (Actg1) knockout leads to much milder phenotypes [2]. Remarkably, this differential role in organism survival is critically dependent on the nucleotide, rather than amino acid sequences: targeted editing of Actb coding sequence by introducing five point mutations to encode *γ*-actin protein (*β*-coded *γ*-actin or Actb*^cg/cg^* ) in mice, which have no *β*-actin protein, does not affect viability [3, 4]. Furthermore, the roles of *β*- and *γ*-actin in cell migration also critically depend on the nucleotide sequence differences and are linked to the differential translation dynamics of Actb and Actg1 [5].

At a first glance, these data suggest that non-muscle actin isoforms are interchangeable at the amino acid level, as long as their nucleotide sequences are mostly intact. However, the four amino acid differences between *β*- and *γ*-actin are conserved among all amniotes, suggesting a separate evolutionary pressure that points to specific role of these amino acid residues in *β*- and *γ*-actin isoforms’ *in vivo* function. In support, Actb*^cg/cg^* mice exhibit stereocilia defects in the inner ear that lead to progressive loss of high pitch hearing [4]. Even though these mice are viable, they likely exhibit other phenotypic changes. Notably, all these changes, if present, would arise because of the amino acid substitutions that convert the entire *β*-actin protein pool in these mice to *γ*-actin. Thus, these mice are an ideal model to address protein-specific functions of non-muscle actin isoforms in vivo.

One of the prominent non-muscle tissues that rely predominantly on non-muscle actins is the retina, in which *β*-and *γ*-actin constitute the majority of the actin cytoskeleton (according to the proteomic data [6, 7]). Photoreceptors in the retina critically depend on actin dynamics for their physiology, in particular, the constant renewal of the disk membranes which provide the light-sensing surfaces in the outer segment (OS). New disk membranes are formed by a special type of actin dependent ectocytosis at the base of the OS [8]. Old disk membranes of the OS tips are digested by the retina pigment epithelium (RPE) through phagocytosis, a process that also relies on the actin cytoskeleton [9]. Proteomic data indicate that both *β*- and *γ*-actin isoforms are present in the photoreceptors [7, 10] and RPE cells [11, 12]. However, the roles played by individual non-muscle actin isoform in maintaining retina structure and function is a mystery.

Here, we investigated the roles of *β*- and *γ*-actin in the retina using the Actb*^cg/cg^* mice. We found that these two actins exhibit significant differences in their localization within the retina layers. The lack of *β*-actin in Actb*^cg/cg^* mice results in accelerated retina degeneration during aging, accompanied by structural defects consistent with defects in the digesting of OS tips by the RPE. Physiologically, replacement of *β*-actin with *γ*-actin protein in Actb*^cg/cg^* mice leads to a significant decrease in rod transducin concentration which causes impaired light sensitivity, measured by both electroretinographs in live animals and electrode recordings of photoreceptors. At the molecular level, several actin binding proteins that mediate the phagocytosis of OS tips by the RPE show differential association for *β*- and *γ*-actin, suggesting their direct involvement in the observed defects in Actb*^cg/cg^* mice. Thus, actin isoform balance is a key factor that mediates normal retina health and slows down age-related retina degeneration.

## Results

### β- and γ-actin prominently localize to the synaptic layers and microvilli of RPE and Müller cells in the retina

Actin cytoskeleton in the retina is composed predominantly of *β* and *γ* non-muscle actins [6, 7], however the distribution of these actin isoforms in the retina layers has not been characterized. To address this, we used actin isoform specific antibodies [13] to analyze the distribution of *β*- and *γ*-actin in the retina sections of dark adapted mice (Figure 1A, S1).

**Figure 1.**
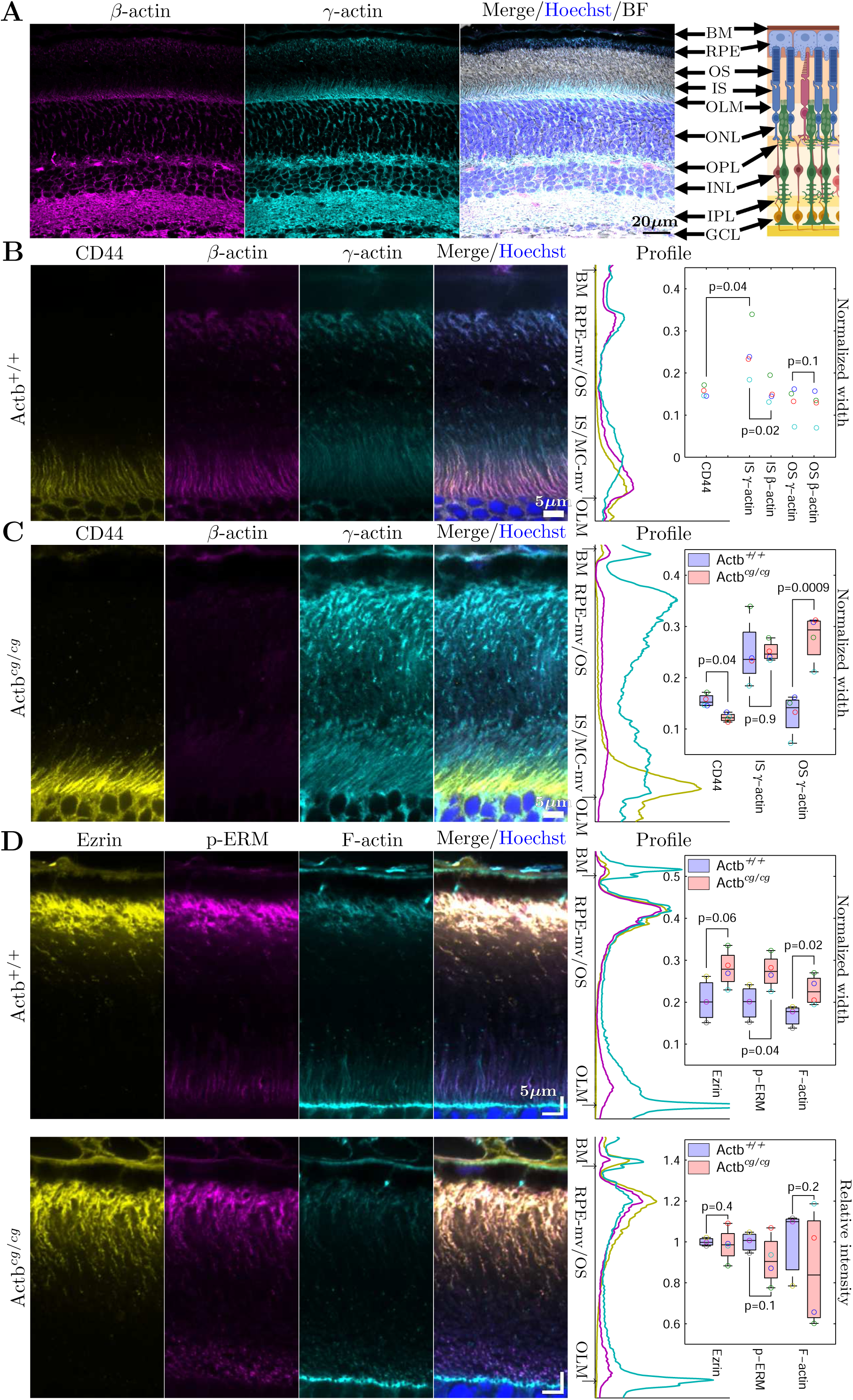
Retina of Actb*^cg/cg^* mouse with *β*-actin replaced by *γ*-actin has abnormal interface between photoreceptors and RPE. (A) Localization of non-muscle actin isoforms in the wildtype retina. Scale bar: 20*µ*m. Pink: *β*-actin. Cyan: *γ*-actin. Blue: DAPI. BM: Bruch’s membrane. RPE: retinal pigment epithelium. OS/IS: outer/inner segment. OLM: outer limiting membrane. ONL/INL: outer/inner nuclear layer. OPL/IPL: outer/inner plexiform layer. GCL: ganglion cell layer. Immunostaining of wildtype (B) and Actb*^cg/cg^* (C) photoreceptors and RPE cells. Yellow: CD44. Pink: *β*-actin. Cyan: *γ*-actin. Scale bar: 5*µ*m. (B,C, right) 1-dimensional projection profiles of staining intensities. (B,C, inset) Quantification of the normalized widths show: in Müller cell microvilli, a significant difference between *β* and *γ* actin isoforms of wildtype but no difference of *γ* actin between wildtype and Actb*^cg/cg^*; and in RPE apical microvilli no significant difference between *β* and *γ* actin isoforms of wildtype but a significant difference of *γ* actin between wildtype and Actb*^cg/cg^* . Each pair is marked with a unique color. The p-values are from paired Student’s t-test with *n* = 4 pairs. (D) Immunostaining of microvilli marker proteins, ezrin (yellow), p-ERM (pink), and F-actin (cyan), shows their co-localization and the increased spread of RPE microvilli of the Actb*^cg/cg^* but no change in staining intensity. (D insets) Each of 3 wildtype and 4 Actb*^cg/cg^* is marked with a unique color. The average width of the distribution profiles at the RPE/OS interface is significantly wider in the Actb*^cg/cg^* than the wildtype (0.26/0.19, p=0.04). The p-values are from one tailed Welch’s t-test. Each box shows the lower quartile, median, and upper quartile values. The maximum length of the whiskers are 1.5 times of the inter quartile range.

Both *β*-actin and *γ*-actin were present in all layers of the retina and showed prominent enrichment in the areas containing the neuronal termini, the outer plexiform layer (OPL) and the inner plexiform layer (IPL), as well as in the areas containing the microvilli of the retinal pigment epithelium (RPE) and Müller cells (MC) intersecting the outer and inner segments (OS and IS) of photoreceptors, respectively. In most of these areas *β*- and *γ*-actin largely colocalized with each other. The only exception was the area just above the outer limiting membrane (OLM), where *γ*-actin covered a broader zone that contains the microvilli of the Müller cells and IS, while *β*-actin localized to a narrower area coinciding with the Müller cell (MC) microvilli (Figure 1A, S1).

To quantify the actin isoform distribution in these areas, we co-stained retina with *β*- and *γ*-actin antibodies and an antibody to CD44, a Müller cell specific marker (Figure 1B). We integrated fluorescence signal for each marker across a wide zone sectioned across the retina layers to produce one-dimensional intensity profiles, and normalized it to the distance between OLM and Bruch’s membrane (BM) to quantify the spread of each peak (Figure 1B, right). The normalization eliminated the unavoidable variations in sectioning angles and gave a proportional size of retinal structures. These measurements confirmed that *β*- and *γ*-actin co-localized at the interface of RPE and OS, while CD44 co-localized with *β*-actin at the interface of MC and IS where both CD44 and *β*-actin distributed closer to OLM and extended significantly less than *γ*-actin (Figure 1B, inset). These results suggested that *β*-actin is enriched in the MC microvilli and *γ*-actin is enriched in the IS, while both are present similarly in the RPE microvilli (Movie M1). Such differences point to potential differences in the roles they play in the retinal structure and function.

### Replacement of β-actin with γ-actin leads to elongation of the RPE microvilli surrounding outer segments of photoreceptors

To test the individual roles of actin isoforms in the retina, we utilized our previously developed mouse model, beta-coded gamma (Actb*^cg/cg^* ) mice, in which *β*-actin gene is edited by introducing five point mutations into its coding sequence to encode *γ*-actin protein ([3] and Figure S2A). In these mice, the non-muscle actin genes are nearly intact, but *β*-actin protein is completely absent, and *γ*-actin protein is produced from both Actg1 (*γ*-actin) gene and the edited Actb (*β*-actin) gene containing five point mutations in the coding sequence. Thus, any phenotypes in these Actb*^cg/cg^* mice are directly linked to the absence of *β*-actin protein, making this the ideal model to dissect the specific role of *β*-actin protein in the functions of the actin cytoskeleton.

Comparison of the wildtype and the Actb*^cg/cg^* (Figure 1B, 1C, Movie M1, M2) confirmed that while in the Actb*^cg/cg^* mice all the *β*-actin protein was replaced with *γ*-actin, the layered structure of the Actb*^cg/cg^* retina looked remarkably normal (Figure S2B). However, the MC microvilli, marked by CD44, had a narrower spread in the Actb*^cg/cg^* compared to the wildtype (Figure 1C, inset). In contrast, the RPE microvilli, marked by *γ*-actin, had a 2-fold wider spread in the Actb*^cg/cg^* retina compared to the wildtype (Figure 1C, right). This shows that loss of *β*-actin protein differentially affects the cellular protrusions in different parts of the retina: while the MC microvilli get shorter, the RPE microvilli get significantly longer.

To confirm that the RPE microvilli were indeed elongated in the Actb*^cg/cg^* compared to wildtype, we stained the retina with antibodies to some of the known RPE microvilli markers [14]: ezrin, p-ERM (phosphorylated ezrin-radixin-moesin), and F-actin (Figure 1D). We found that they had robust co-localization at the RPE/OS interface of both the wildtype and the Actb*^cg/cg^* (Figure 1D, left), representing the RPE microvilli distribution. Analysis of these data confirmed that the average width of the distribution was significantly wider in the Actb*^cg/cg^* than in the wildtype (Figure 1D, top inset). The staining intensities of the wildtype and the Actb*^cg/cg^* at the RPE/OS interface were not significantly different for all three markers (Figure 1D, bottom inset). However, the staining intensity for F-actin, but not ezrin or p-ERM, negatively correlated with the average width (Pearson/Spearman correlation coefficient -0.76/-0.86 with p-value 0.045/0.014), indicating that for the Actb*^cg/cg^* RPE microvilli or their actin cytoskeleta, the more elongated they are, the less dense they are.

Thus, replacement of *β* actin with *γ* actin affects the actin cytoskeleton at the RPE/OS interface.

### Actb^cg/cg^ mouse retina has defects in RPE and rod outer segments

To look at potential structural changes in the retina layers, we used optical coherence tomography (OCT) for non-invasive imaging of the structure of the retina with an infrared (IR) light source. OCT measures IR light reflected from various retinal layers. The most noticeable difference between the Actb*^cg/cg^* and wildtype OCT images was near RPE/OS interface, marked by the upward arrows in Figure 2A. We calculated the contrast between the peak of the RPE band and the peak and the OS band. This contrast was significantly higher in the Actb*^cg/cg^* than the wildtype under both light and dark adapted conditions

**Figure 2.**
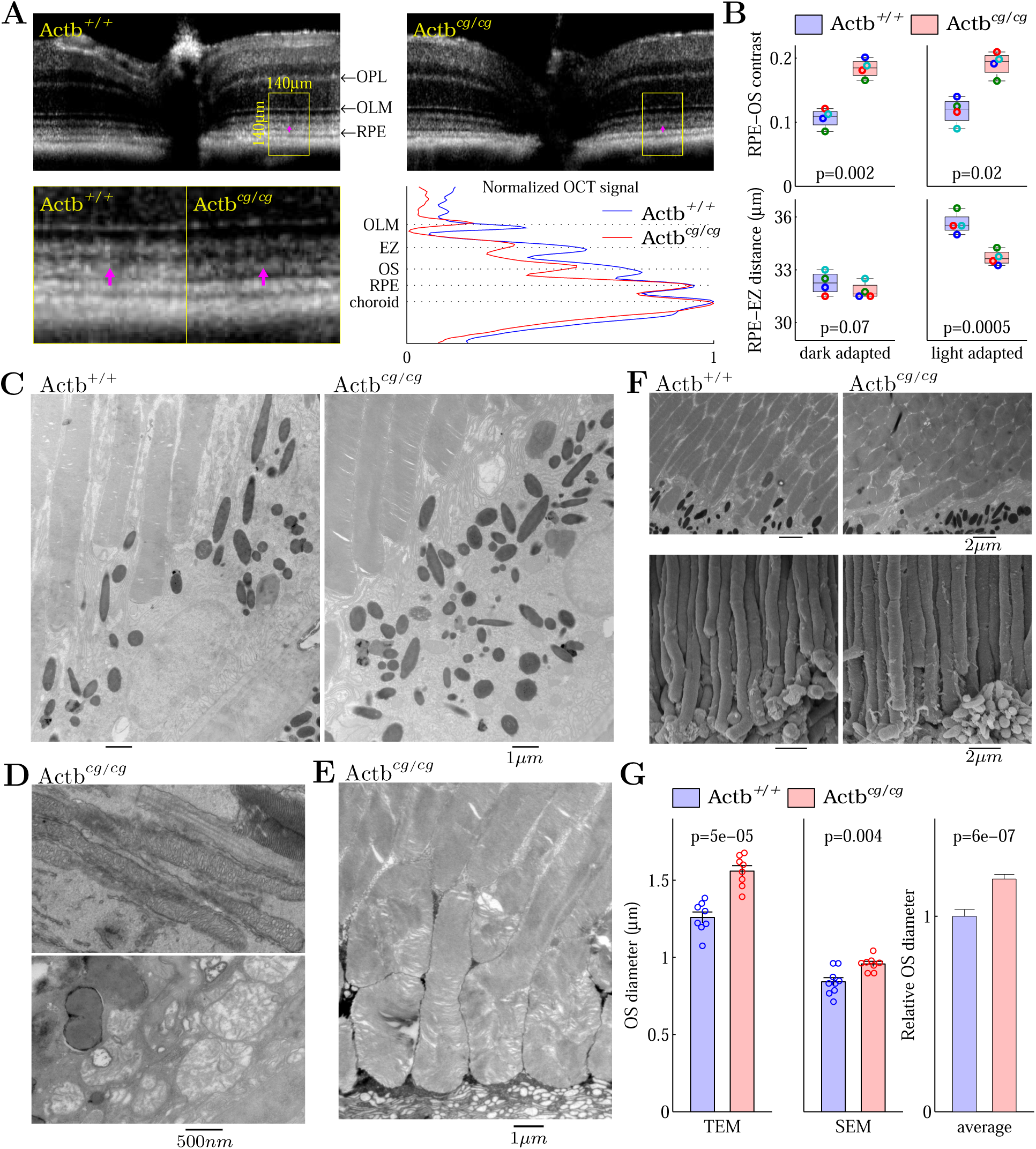
Actb*^cg/cg^* retina had increased number of granules and defective mitochondria in the RPE and swelling OS. (A) The OCT images of the retinas of a littermate pair of wildtype and Actb*^cg/cg^* mice show the difference in signal contrast between the RPE band of the OS band (marked with pink arrows). The magnified patches from wildtype and Actb*^cg/cg^* and their projection profiles from OLM to choroid are shown below. The OS band marked here is known in OCT literature as interdigitation zone or OS-tips band, where RPE microvilli encase OS. The EZ band marked here is known in OCT literature as ellipsoid zone or IS/OS band, which resides at the IS/OS interface. (B) The Actb*^cg/cg^* had significantly higher contrast between RPE band and OS band in both dark and light adapted retinas (top) and significantly shorter distance between RPE band and EZ band for light adapted retinas but similar one for dark adapted retinas (bottom). Each pair is marked with a unique color. The p-values are from paired Student’s t-test with *n* = 4 pairs. (C) TEM images of a matching pair of wildtype and Actb*^cg/cg^* retinas show increased number of granules in Actb*^cg/cg^* . (D) Actb*^cg/cg^* had bloated mitochondria in RPE (bottom) but not in IS (top). (E) Actb*^cg/cg^* OS had swellings and convolutions. (F) Both TEM (top) and SEM (bottom) show significantly increased diameters of Actb*^cg/cg^* OS (G). Note: the OSs in the SEM images were visibly thinner than in the TEM, due to shrinkage of the cellular structures during SEM preparation, The p-values in (G) are from Welch’s t-test.

of the retina (Figure 2B, top). Using the choroid band as the reference, we also found that under dark adaptation the OS band was significantly darker and the RPE band was significantly brighter in the Actb*^cg/cg^* compared to the wildtype, indicating defects in both the OS and the RPE, while the OLM band and EZ band did not change significantly (Figure S3A). The relative locations of various layers did not change between Actb*^cg/cg^* and wildtype retinas under dark adaptation (Figure S3B). However, the distance between the peak of RPE band and the peak of EZ band in the Actb*^cg/cg^* was significantly shorter (indicating shorter OS) compared to the wildtype under light adaptation (Figure 2B, bottom), corroborating a specific defect of the rod OS shown later in the results. To look at ultra-structures of the OS and the RPE, we used transmission and scanning electron microscopy (TEM and SEM). We found that the RPE cells of Actb*^cg/cg^* retina contain more lipofuscin and melanolipofuscin granules (Figure 2C), most of which were particles *∼*1 micrometer in size that are well suited to contribute to OCT under IR light. In addition, while the mitochondria in the inner segments looked normal (Figure 2D, top), many mitochondria in RPE were bloated (Figure 2D, bottom), indicating the RPE cells were selectively affected in Actb*^cg/cg^* retina. There was no obvious defect at the OS/IS interface (Figure 2D, top). However, on the side of the RPE/OS interface, the diameter of Actb*^cg/cg^* rod OS was larger, while the distance between rods remained the same, and as a result the microvilli appeared tightly squeezed between adjacent rods (Figure 2C, F). Notably, while the inner segments of wildtype rods are normally tightly packed, the outer segment of a rod only occupies *∼*50% of the cross-section *area* of the inner segment [15], an arrangement that allows each rod in the wildtype retina to be surrounded by several RPE microvilli. The Actb*^cg/cg^* rods still had space between them, but the space appeared tighter. In addition to the abnormally large OS diameter, some of the rod OS tips were not “rod” shaped but instead appeared convoluted (Figure 2E). These changes of OS likely contributed to the decreased reflectance of the OS band in the Actb*^cg/cg^* OCT and indicate a likely defect in phagocytosis of rod OS by the RPE.

To quantify the OS diameters of the rods, we used TEM images which transected the OS and measured the short axis of the oval shaped transection (Figure 2F, top). The diameter of the Actb*^cg/cg^* OS was significantly larger than the wildtype under dark adaptation (Figure 2G, left). To further confirm this, we used SEM images which were parallel to the rod OS and measured the diameter of OS at its widest, unobscured stalk away from its tip (Figure 2F, bottom). The diameter of the Actb*^cg/cg^* OS by this measure was also significantly larger than the wildtype (Figure 2G, center). On average, the Actb*^cg/cg^* OS diameter was *∼*20% larger than the wildtype (Figure 2G, right).

### Actb^cg/cg^ retina accelerates age-related degeneration

The lipofuscin accumulation and the bloated mitochondria in the RPE (Figure 2C, D) commonly develop during age-related degeneration of the retina [16]. The swellings and convolutions of rod outer segments (Figure 2E, F) have also been reported in the aging rods of human [17]. Thus, we investigated the Actb*^cg/cg^* retina for another hallmark of aging: age-dependent accumulation of autofluorescent microglia in the subretinal space between the OS and the RPE [18, 19, 20, 21].

*In vivo* retinal autofluorescence was imaged using the confocal scanning laser ophthalmoscope (SLO). By adjusting the confocal aperture, we focused at the RPE/OS interface and found a striking number of autofluorescent spots in 12 month old Actb*^cg/cg^* retina which were hardly seen in the age-matched wildtype (Figure 3A, left). In fact, with the exception of one outlier, the wildtype retinas exhibited over an order of magnitude fewer spots at the age of 12 month (Figure 3B, left) and only reached reached the level of spots comparable to 12-month-old Actb*^cg/cg^* retinas at 21-26 months of age (Figure 3A, B). These spots develop during normal aging of the retina due to the increased migration of microglia cells to the subretinal space, which ingest OS disks and RPE granules, and accumulate autofluorescent lipofuscin [19, 20]. The Actb*^cg/cg^* retinas developed these spots much earlier. In Actb*^cg/cg^* mice, examples of such microglia cells were also visible at both the gross histology level and at the ultrastructural level as early as 8-month-old (Figure 3C, D). Thus, replacement of *β*-actin with *γ*-actin in mice leads to the acceleration of age-related retinal degeneration.

**Figure 3.**
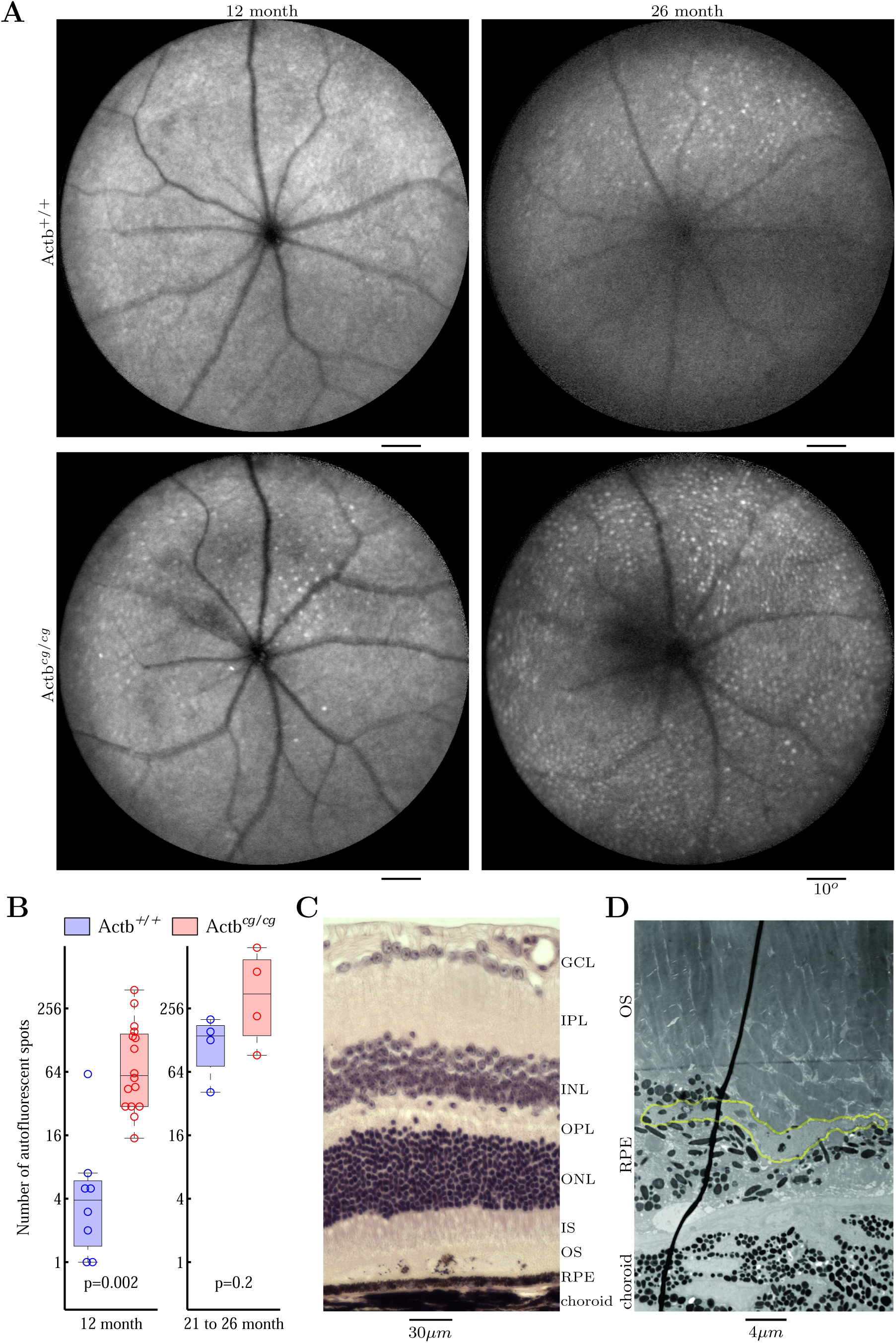
Actb*^cg/cg^* retinas have age-related microglia infiltration into subretinal space at earlier age than wildtype. (A) Confocal SLO imaging of the subretinal space shows autofluorescent spots on the retinas of a matching pairs of wildtype (top) and Actb*^cg/cg^* (bottom) at the age of 12 month (left) and 26 month (right). Notably, these spots are uniform in size and shape—same as those shown previously of the age-related microglia infiltration into subretinal space in normal aging mice [20]—very different from the great variation observed in genetic diseases such as rd8 [22] and traumatic injury [23, 24]. (B) The Actb*^cg/cg^* retinas at 12 month and 2 year old have similar amount of autofluorescent spots in comparison to wildtype around 2 years old, significantly more than wildtype at 12 month. (C) H&E section of a Actb*^cg/cg^* retina of 8 month old male shows a microglia infiltrated into the subretinal space. (D) TEM section of a Actb*^cg/cg^* retina of 8 month old female shows the part of a microglia in the subretinal space (outlined in green). The higher resolution TEM images for this is shown in Figure S4.

### Actb^cg/cg^ rod outer segments have decreased transducin concentration

Under the dark adaptation, the OCT showed no difference in the distance from the distal edge of EZ band to the apical edge of RPE band between the Actb*^cg/cg^* and the wildtype (Figure 2A, S3B). This indicates that the outer segment equivalent length [25] did not change. Therefore, the enlarged diameter of Actb*^cg/cg^* rods revealed by TEM and SEM (Figure 2F) means that the Actb*^cg/cg^* rods had increased volume.

To test if the levels of proteins involved in rod phototransduction also increased with the increased volume of the Actb*^cg/cg^* rods, we first quantified the levels of rhodopsin and G*β*1 of the rod transducin trimer using immunofluorescence (Figure 4A). Surprisingly, neither of these proteins showed any significant change in the average staining intensity over the retinal layer covering the outer segments (Figure 4B, top). There was also no difference in OS length marked by either rhodopsin or G*β*1 (Figure 4B, bottom), confirming what we observed in OCT. To ascertain that the total rod transducin available to phototransduction did not increase in the Actb*^cg/cg^* , we used the another method in which mice were adapted to the room light (*>*30*cd/m*^2^ in the cage) for at least 4 hours. Under these conditions all rod transducins are released from the OS disk membranes resulting in no rod responses [26, 27]. We then quantified the amounts of transducins in the supernatant of the retinal lysates using mass spectrometry. None of 3 components of the rod transducin trimer had significant increase in the Actb*^cg/cg^* supernatants (Figure 4C). Taking the rod OS lengths measured in OCT, the rod OS diameters measured in TEM and SEM, and the amount of rod transducins measured in mass spectrometry, we calculated the transducin concentration in the OS under dark adaptation. It was significantly lower in Actb*^cg/cg^* than in wildtype (Figure 4D, left).

**Figure 4.**
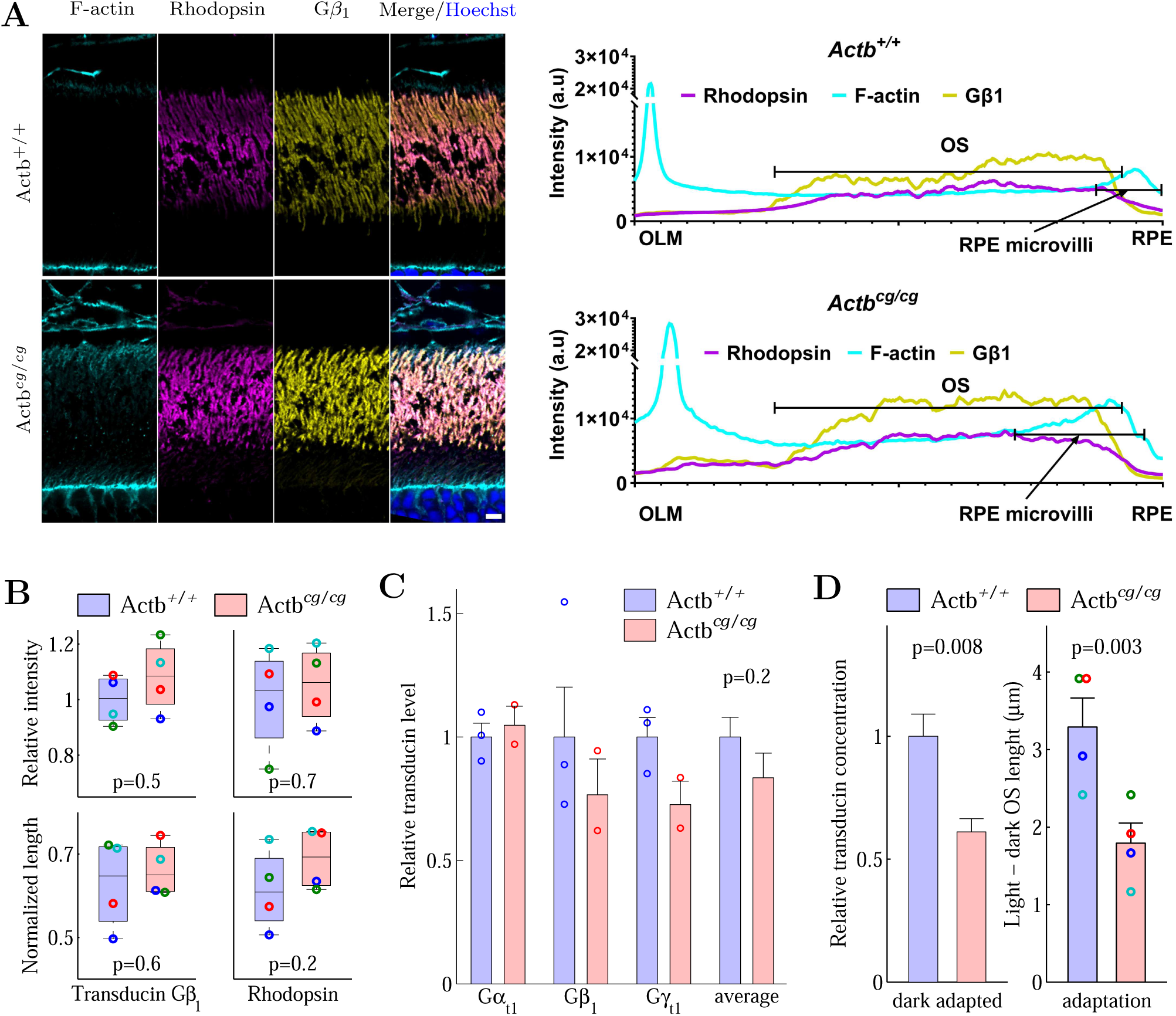
Decreased transducin concentration of Actb*^cg/cg^* retina. (A) Staining of rhodopsin and rod transducin *β* of dark adapted retinas. Scale bar: 5*µ*m. (B) Comparing Actb*^cg/cg^* to wildtype, there is no difference in the staining intensity (top) and no difference in the normalized (to the distance between OLM and BM) OS length (bottom), which also means there is no difference in the OS length since it is normalized to the distance between OLM and BM, which did not change significantly for dark adapted Actb*^cg/cg^* and wildtype retinas. (C) The *amount* of transducins involved in phototransduction in rod OS in the retina did not change significantly in Actb*^cg/cg^* . (D) The relative *concentration* in the rods (left) decreased significantly in Actb*^cg/cg^* and the expansion of the length of rod OS from dark to light adaptation was significantly less in Actb*^cg/cg^* (right). Eyes of each pair of Actb*^cg/cg^* and wildtype are marked with a unique color. The p-values for (B) and (D, right) are from paired Student’s t-test (*n* = 4). The p-values for (C) and (D, left) are from Welch’s t-test.

Additional evidence that the Actb*^cg/cg^* rod OS had lower rod transducin concentration came from the adaptation experiment (Figure 2B). The rod OS length increases with light adaptation in comparison to dark adapted OS [28, 29] and the amount of increase is positively correlated with the concentration of transducin released from the disk membrane of the rod OS [29]. For both the Actb*^cg/cg^* and the wildtype, the OS length increased with light adaptation (Figure 2B, bottom) but the increase for the Actb*^cg/cg^* was significantly smaller than the increase for the wildtype (Figure 4D, right), proportional to the calculated transducin concentration (Figure 4D, left).

Reduced transducin *concentration* would result in slower responses to light, since the phototransduction in the outer segments is time-limited by the diffusion of activated transducins [30].

### Actb^cg/cg^ rod outer segments have slower light responses

To test the functional effects of *β*-actin replacement on light response in photoreceptors, we used an established protocol of recording electroretinographs (ERG) on dark adapted mice. In this protocol, after a light flash, the initial downward A-wave (Figure 5A, left) is followed by the upward B-wave (Figure 5A, middle). For dark adapted retina, A-waves originate mostly in the outer segments of the rods, while B-waves stimulated by dim flashes are dominated by the rod bipolar cells [26, 27].

**Figure 5.**
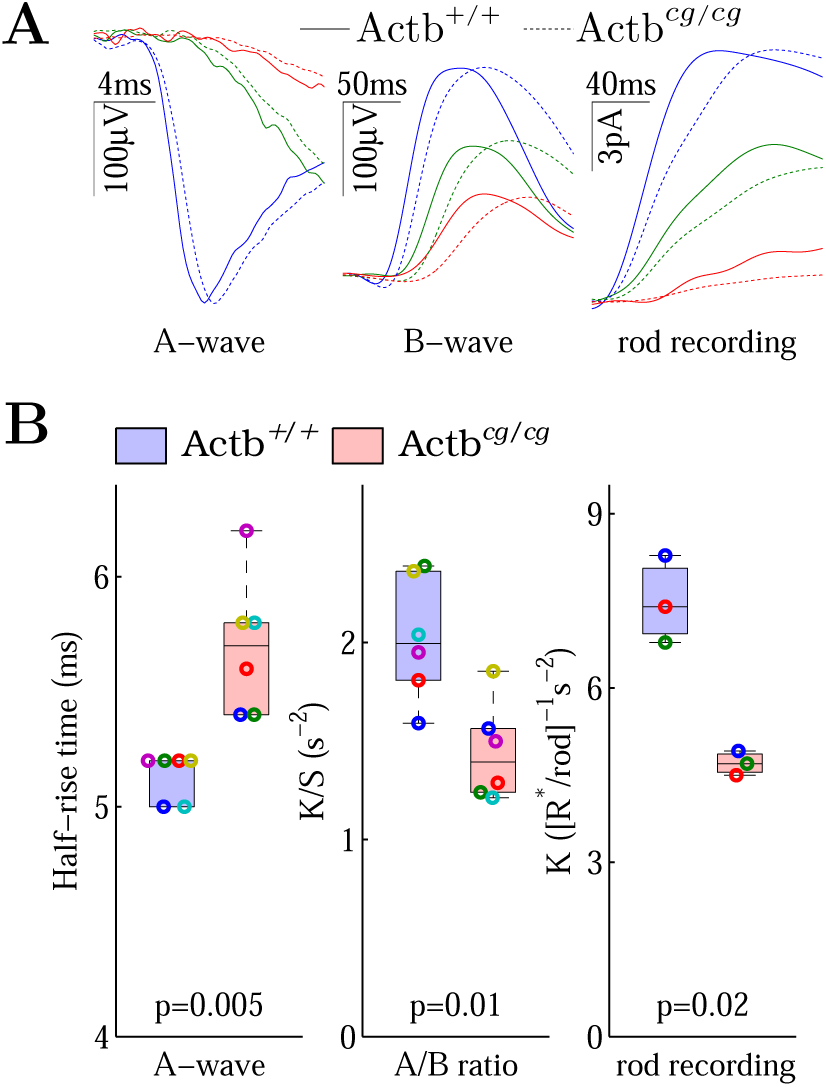
Slower light responses and reduced sensitivity of Actb*^cg/cg^* retina. (A) From left to right, rod ERG A-wave, B-wave and suction-pipette recording, from wildtype (solid line) and Actb*^cg/cg^* (dashed line). The red, green, and blue colored curves are the responses to the light stimulation of increasing intensities: 1, 4, 300 *cd·s/m*^2^ for A-wave, 0.00065, 0.00215, 0.01076 *cd·s/m*^2^ for B-wave, and 4, 16, 72 *R^∗^/rod* for rod recording. *R*∗*/rod*: rhodopsin photoisomerization calculated for 500nm green light and 0.5*µm*^2^ absorption cross-section per rod. (B, left) The half-rise times are significantly longer for Actb*^cg/cg^* retina. (B, center) The ratio of A-wave and B-wave response sensitivities. Each pair of data points of the same color represents a wildtype/Actb*^cg/cg^* littermate pair. The p-values are from paired Student’s t-test with *n* = 6. (B, right) The sensitivities of rod recording from retinal slices of 3 wildtype/Actb*^cg/cg^* pairs. The blue and green pairs were recorded with suction pipettes and the red pair with multi-electrode array. The p-values are from paired Student’s t-test with *n* = 3.

Several light flashes of different intensities were used to generate a family of responses for each of the A-wave and B-wave, including a flash intensity which generates a maximum amplitude response. The maximum amplitudes of responses were similar between wildtype and Actb*^cg/cg^* mice (Figure S5A), however the Actb*^cg/cg^* mice exhibited these responses with a slower time course (Figure 5A). To quantify the time differences, we determined the half-rise time, from the baseline to the valley for A-wave. The half-rise time was significantly longer for the Actb*^cg/cg^* mice (Figure 5B, left), indicating that the light responses were slower in their rod outer segments. To further determine the cause of the slower light responses, we fitted the A-wave and B-wave response families to the established models of rod phototransduction and rod bipolar responses. The resulting A-wave sensitivity *K* was lower in the Actb*^cg/cg^* while the resulting

B-wave sensitivity *S* did not change (Figure S5B). The ratio of those two (*K/S*) was significantly smaller for the Actb*^cg/cg^* (Figure 5B, center). Since both *K* and *S* are proportional to the actual rate of rhodopsin photoisomerization during the same experimental session, the ratio eliminates any variation due to light illumination, and, more importantly, excludes the possibility that the observed slower A-wave responses were solely caused by a reduction of rhodopsin photoisomerization in the Actb*^cg/cg^* rods (in which case, the *K/S* ratio would not change). Such differences were not present in mice less than 4 month old (Figure S5C), indicating an age-related development of the defect in the Actb*^cg/cg^* rods.

To make sure that the observed differences were caused by rods, we directly measured the photocurrent of dark-adapted rods (Figure 5A, right) in retinal slices as well as A-wave responses from flat mounted retinal patches at very low levels of light stimulation. We found that the rod sensitivity *K* was significantly lower in the Actb*^cg/cg^* than the wildtype (Figure 5B, left). Thus, replacement of *β* actin with *γ* actin in mice leads to slower light responses in the outer segments of the Actb*^cg/cg^* rods compared to the wildtype.

### Replacing β-actin with γ-actin significantly changes the interaction between actin and actin-binding proteins

To test the molecular mechanisms underlying the retina defects in the Actb*^cg/cg^* mice, we analyzed protein composition of the actin cytoskeleton and actinbinding proteins by performing a pulldown from total retina lysates using biotin-phalloidin, which interacts selectively with F-actin, and analyzed the composition of the actin-binding proteins in these pulldowns using mass spectrometry.

We specifically tested 10 actin-binding proteins known to be important for RPE microvilli and OS phagocytosis: annexin A2 and A5 [31, 32], ezrin [33, 34], non-muscle myosin-IIa and -IIb [35, 36], unconventional myosin-VI and -VIIa [37, 38, 39, 40], talin and vinculin [41, 42]. We were able to reliably quantify 9 of them from mass spectrometry data and found that all except one (myosin-IIa, encoded by Myh9) differentially bind to F-actin in wildtype and Actb*^cg/cg^* retina

(Figure 6A). Since the actin cytoskeleton in the Actb*^cg/cg^* retina is composed of 100% *γ* actin, while in wildtype it contains a mixture of *β* and *γ*-actin, this change likely reflects differential interaction of these proteins for *γ*-actin and *β*-actin (in Figure 6A, protein/actin ratio increase and decrease, respectively; also see Table 1 and 2 for details). None of these proteins showed a change in level in the retina lysates (Figure S6), confirming that these changes in the pulldown were a result of altered F-actin interaction.

**Figure 6.**
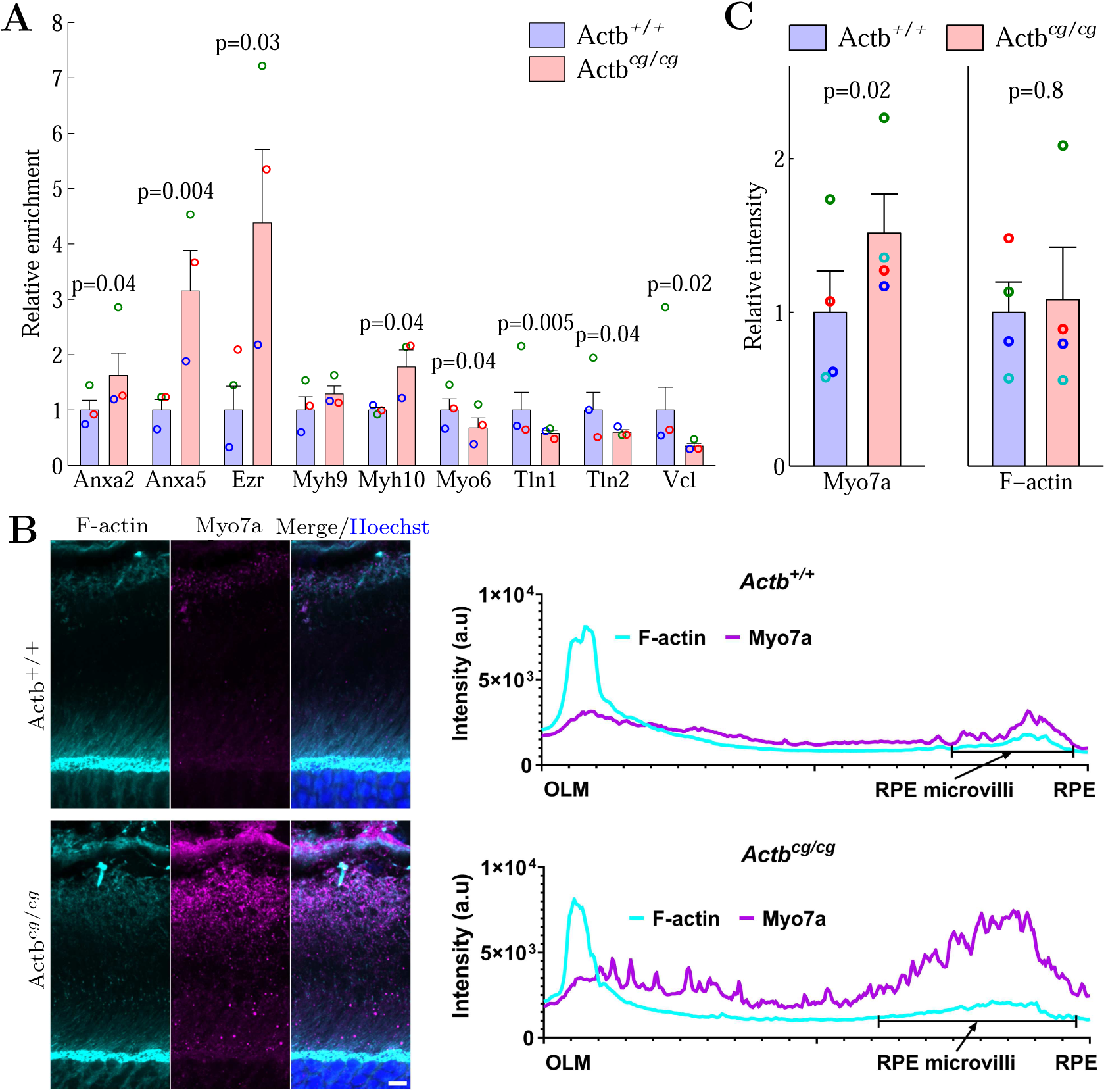
Differential interaction of cytoplasmic actins with actin associated proteins. (A) Different interaction with *β*- and *γ*-actin was shown as the relative enrichment of a protein in phalloidin-pulldown of the lysates of wildtype and Actb*^cg/cg^* retina. The p-values are from Table 1 and 2. (B) The immunostaining of myosin-VIIa indicated the increased interaction of myosin-VIIa with *γ* actin (C). Scale bar: 5*µ*m. The p-values are from paired Student’s t-test with *n* =4 pairs. Myh9/Myo7a: gene symbol of myosin-IIa/myosin-VIIa protein. For other gene symbols in (A), Table 1 and 2 show the protein names.

**Table 1:** Actin associated proteins with significantly increased protein to β- and γ-actin ratio in Phaloidin pulldown of retinal lysates. The p-value is from Actb^cg/cg^ and wildtype paired Student’s t-test of the ratio of a protein to β- and γ-actin (n=3 pairs), except for Wdr1 which is from Spearman’s correlation coefficient of a protein with γ-actin (n=6 samples).

| Gene symbol | Fold change | p-value | MS/MS count | Unique peptide | Score | Protein name |
| --- | --- | --- | --- | --- | --- | --- |
| Anxa2 | 0.70 | 4.4e-02 | 177 | 28 | 323 | Annexin A2 |
| Anxa5 | 1.66 | 4.5e-03 | 87 | 21 | 323 | Annexin A5 |
| Anxa6 | 1.16 | 2.9e-02 | 120 | 25 | 164 | Annexin A6 |
| Ezr | 2.13 | 3.5e-02 | 18 | 8 | 36 | Ezrin |
| Msn | 1.50 | 3.5e-02 | 53 | 24 | 93 | Moesin |
| Myh10 | 0.83 | 4.2e-02 | 31 | 27 | 196 | Myosin-10, non-muscle myosin-IIb |
| Myh11 | 1.37 | 1.1e-02 | 52 | 43 | 323 | Myosin-11 |
| Wdr1 | 2.67 | 4.2e-02 | 26 | 13 | 59 | WD repeat-containing protein 1 |

**Table 2:** Actin associated proteins with significantly decreased protein to β- and γ-actin ratio in Phaloidin pulldown of retinal lysates. The p-value is from Spearman’s correlation coefficient of a protein with γ-actin (n=6 samples), except for Myo6 which is from Actb^cg/cg^ and wildtype paired Student’s t-test of the ratio of a protein to β- and γ-actin (n=3 pairs).

| Gene symbol | fold change | p-value | MS/MS count | Unique peptide | Score | Protein name |
| --- | --- | --- | --- | --- | --- | --- |
| Cct5 | -0.81 | 1.9e-02 | 222 | 34 | 323 | T-complex protein 1 subunit epsilon |
| Iqgap1 | -0.85 | 1.9e-02 | 251 | 68 | 323 | Ras GTPase-activating-like protein |
| Myo6 | -0.56 | 4.1e-02 | 37 | 22 | 61 | Unconventional myosin-VI |
| Tln1 | -0.78 | 4.8e-03 | 341 | 89 | 323 | Talin-1 |
| Tln2 | -0.74 | 4.2e-02 | 235 | 67 | 323 | Talin-2 |
| Utrn | -0.56 | 1.9e-02 | 37 | 18 | 54 | Utrophin |
| Vcl | -1.52 | 1.9e-02 | 122 | 54 | 323 | Vinculin |

The only one left of the 10 proteins was myosin-VIIa. Only a small amount of myosin-VIIa was detected in our F-actin pulldowns, preventing proper quantification. This low yield is likely due to the fact that the major myosin-VIIa structures in the retina are large, and thus end up in the insoluble fraction during the preparation of the cell extract. However, immunostaining showed a significantly higher intensity level of myosin-VIIa in the Actb*^cg/cg^* RPE compared to wildtype. This higher intensity myosin-VIIa staining also occupied a broader zone within the retina layers, consistent with the increased length of the RPE microvilli in Actb*^cg/cg^* retina (Figure 2D). Factoring this increase in both length and intensity of myosin-VIIa, these data indicate that myosin-VIIa is enriched in Actb*^cg/cg^* retina due to its increased interaction with *γ*-actin (Figure 6B, C). Thus replacing *β*-actin with *γ*-actin significantly changed the interaction between the actin cytoskeleton and proteins involved in phagocytosis of the OS by the RPE. We propose that a combined change in the interaction of these proteins with actin leads to weaker and more extended contacts of RPE microvilli along rod OS, which accounts for the major retina defects we observe in Actb*^cg/cg^* mice.

## Discussion

Our study demonstrates that editing of the *β*-actin gene by introducing five point mutations that convert the protein product from this gene to non-muscle *γ*-actin (*β*-coded *γ*-actin or Actb*^cg/cg^* ) results in a major impact on the retina, the visual input part of the brain. Previous studies showed that nucleotide- and protein-level changes contribute to the unique functions of non-muscle actin isoforms in vivo, but none ever showed that the four amino acids difference between the two actin isoforms are important to the structural integrity and physiological function of the retina.

Our data show that Actb*^cg/cg^* retina exhibit accelated age-related degeneration and impairments in light responses. These are major physiological consequences of a substitution in four conservative amino acid residues between the two closely related non-muscle actin isoforms. While Actb*^cg/cg^* mice, unlike *β*-actin knockout mice, are viable and relatively healthy at young age [3, 4],our present results show that these mice have major defects at the tissue level during aging, including both structural and functional defects. The outer segments (OS) of mutant rod photoreceptors have a bigger volume, accompanied by a reduced concentration of transducin and a slower light response. The Actb*^cg/cg^* retinal pigment epithelium (RPE) has accelerated lipofuscin accumulation and mitochondrial dysfunction (Figure 2C, 2D, and 3), two defects normally associated with retina aging and suggested to synergistically impair the OS phagocytosis by the RPE [43]. These OS and RPE defects are intimately detrimental to a crucial function of the actin dynamics, the continued renewal of the OS disk membranes to maintain photoreceptor viability [8, 9]. The RPE microvilli in Actb*^cg/cg^* mice ensheathe the OS tips further along the OS under dark adaptation (Figure 1C, 1D), reminiscent of the wildtype under light adaptation [28]. While the OS expansion under light adaptation is retarded in Actb*^cg/cg^* (Figure 2B, 5D), the OS tips in Actb*^cg/cg^* are bulging and often convoluted (Figure 2E, 2F). Most of these defects become observable in Actb*^cg/cg^* mouse retina at ages between 6 to 12 months, while similar defects in wildtype develop much later.

Such isoform-specific defects in structure and function at the tissue level are the likely reason that the difference between the N-terminal sequences of *β*- and *γ*-actin is preserved throughout the evolution of the amniotes. An important difference between the *β*- and *γ*-actin isoforms is their differential interaction with other proteins important to the actin dynamics. Such differential interaction is expected due to differences in the sequences of the *β*- and *γ*-actin N-termini, which contain conserved stretch of either aspartic or glutamic acid, resulting in predicted alterations in the strength of the negative charge. In addition, *β*-actin undergoes a more extensive N-terminal processing compared to*γ*-actin, and this processing is amino acid dependent [44].

To find out the differential interaction between the *β*- and *γ*-actin interaction with other proteins, we did the pulldown experiment. The relative amount of each actin-binding protein in retinal lysates in comparison to the amount of the same protein in phalloidin pulldown from these lysates is an indicator of the interaction of this protein with F-actin. By doing this comparison using retinas with and without *β*-actin, we detected several proteins that exhibit different interaction with the actin filaments in wildtype and Actb*^cg/cg^* retina (Figure 6), suggesting a differential interaction of these proteins with *β*- and *γ*-actin and pointing to possible molecular mechanisms of the observed phenotypes. First, through *α*v*β*5 integrin on apical RPE membrane [45, 46, 47], talin and vinculin connect OS with RPE actin cytoskeleton [42], therefore the weak-ened binding of talin and vinculin with F-actin in the Actb*^cg/cg^* likely interferes with the “molecular clutch function” [48, 49] of this complex. Second, because ezrin promotes morphogenesis of apical microvilli of RPE [33], the increased interaction of ezrin with F-actin in the Actb*^cg/cg^* can lead to increased length of the microvilli. Third, the overall increase (with longer microvilli) of the phosphorylated ERM (p-ERM shown in Figure 1D) in the Actb*^cg/cg^* can lead to decreased activation of non-muscle myosin II [50]. Decreased activation of non-muscle myosin II hampers the clustering of integrin, talin, vinculin, and F-actin at phagocytic cups [42] and causes significant phagocytic defects [35]. Thus in the Actb*^cg/cg^* retina, RPE microvilli have more freedom to extend further along OS and phagocytosis of OS tips by RPE is retarded (Figure 7).

**Figure 7.**
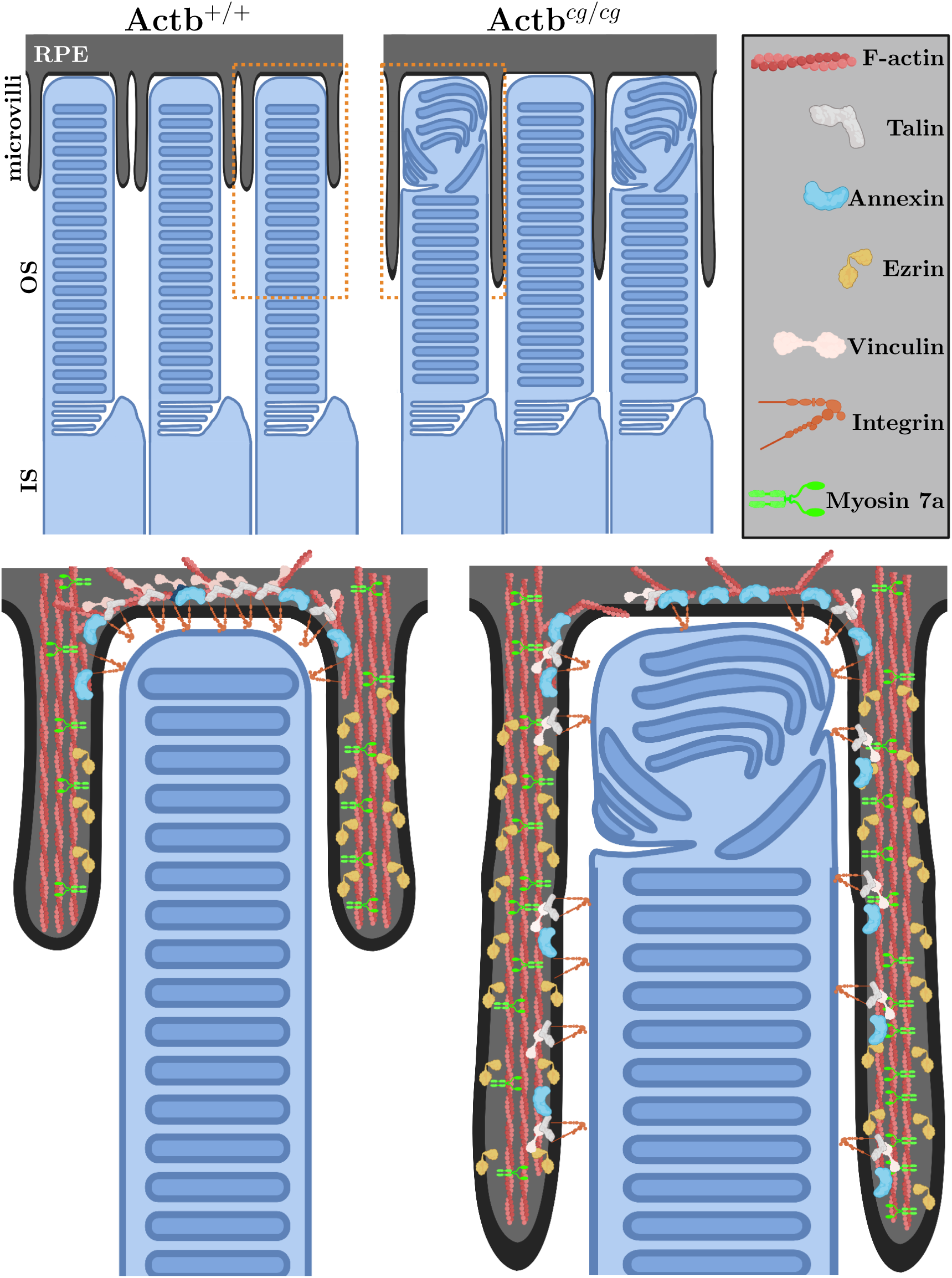
The molecular mechanism for the Actb*^cg/cg^* changes at the interface of RPE and OS.

Annexin A2 and A5 both show higher binding to the *γ*-only Actb*^cg/cg^* actin. Annexin A2 regulates the internalization of newly phagocytosed outer segments [31]. Annexin A5 regulates the binding of microvilli membrane to the OS tips destined for phagocytosis [32]. Their preferred interaction with *γ*-over *β*-actin in the retina is reason enough to change the phagocytosis in the mutant without *β*-actin. While the preference of annexin A2 of *γ*-over *β*-actin is a new discovery, the fact that annexin A5 prefers to bind *γ*-over *β*-actin was previously shown in platelets [51].

The decreased interaction of myosin-VI with *γ*-actin is likely to negatively affect phagocytosis, since myosin-VI is the only retrograde actin motor involved in OS phagosome transport and maturation in RPE and its downregulation was previously shown to suppress OS phagosome degradation [37, 38]. Interestingly this is opposite to myosin-VIIa, known to prefer interaction with *γ*-actin [52]. In the RPE, myosin-VIIa participates in the melanosome localization apically A. [39] and in the phagosome delivery basally [40, 53]. Our phalloidin pulldown only yielded a few MS/MS counts of myosin-VIIa; however, the significantly higher level of myosin-VIIa staining in RPE from Actb*^cg/cg^* retina is consistent with the increased myosin-VIIa interaction with F-actin made entirely of *γ*-actin (Figure 6B, 6C).

A number of proteins identified in our pulldowns as differential interactors with *β*- and *γ*-actin are known to affect mitochondria. Non-muscle myosin IIb and *β*-actin connect mitochondria to actin cytoskeleton and contribute to mitochondrial DNA maintenance and segregation [54], Annexin A6 regulates mitochondria morphogenesis [55], while myosin-VI is required for degradation of damaged mitochondria [56]. In addition, some *β*-actin might reside within mitochondria [54] and regulate mitochondrial DNA transcription [57]. Changes in these proteins constitute possible causes of the mitochondrial phenotype in the mutant RPE (Figure 3E).

One of the proteins differentially interacting with F-actin in our pulldowns is especially intriguing: ezrin is more abundant in the pulldowns from Actb*^cg/cg^* retina and thus shows preference for *γ*-actin. Since ezrin promotes morphogenesis of the apical microvilli of the RPE [33], this differential interaction constitutes a possible reason for the increased width of the ezrin-positive zone in Actb*^cg/cg^* compared to wildtype, seen in the immunostaining of ezrin as well as other associated proteins (Figure 2). This may seem to contradict a previous finding that ezrin prefers *β*-actin [58]. Our explanation of this apparent contradiction is that the preference seen in this previous study is actually a reflection of the activation location of ezrin. It is known that ezrin activation requires phosphorylation and local phosphocycling restricts ezrin function to the apical aspect of epithelial cells [59]. In gastric epithelial cells, *β*-actin happens to localize near the apical membrane and thus the apparent preference of ezrin for *β*-actin [58] is likely due to this preferential localization rather than the actual binding affinity. In our case, only *γ*-actin is present in the mutant while both *β*- and *γ*-actin are present in wildtype. Thus, the preference of ezrin for *γ*-actin in our pulldown experiment could be the result of ezrin binding better with *γ*-actin and/or ezrin in Actb*^cg/cg^* being more phosphorylated at apical RPE. It is possible that the ezrin homolog moesin prefers *γ*-actin for the same reason(s) (Table 1), since moesin is also found in apical microvilli [11].

It has been previously found that Actb*^cg/cg^* mice experience progressive hearing loss and progressive disorganization of the microvilli in the inner ear [4]. In the retina, in addition to the microvilli elongation at the RPE and OS interface, we found that the microvilli at the IS and MC interface, appear shorter in Actb*^cg/cg^* (Figure 1C, inset). It is possible that the mechanisms underlying those phenotypes are driven by similar mechanisms and ultimately related to differential interactions of key binding proteins with *β*- and *γ*-actin.

Surprisingly, among all the actin binding proteins (see Supplement 1), we only found 15 with significant change in the F-actin pulldown experiment (Table 1 and 2). 10 of these are discussed above as likely related to the phenotypes observed in the RPE—in addition to myosin-VIIa. It is possible that some of those other proteins also contribute to the RPE dysfunction —e.g., changes in calcium concentration and phosphorylation which in turn impact the interaction of the others with the actin cytoskeleton. Those are exciting future directions to explore.

## Methods

### Ethical approval and accordance

Procedures involving animals were performed in accordance with National Institute of Health guidelines and the protocol was reviewed and approved by the Institutional Animal Care and Use Committee of the University of Pennsylvania. The total number of animals used in the present study was 24 wildtype (Actb^+*/*+^, WT) mice and 24 *β*-actin mutant (Actb*^cg/cg^* , Mut) mice. The earliest age used for the experiments was 3 months old while the majority was older than 6 months; both male and female mice were used.

### Generation of β-actin mutant mice

*β*-actin mutant mice were generated as described in [3] and maintained inC57BL/6J (Jackson Laboratory) background. Mice were bred and maintained according to approved animal protocols.

### Immunofluorescence and electron microscopy

Mice were euthanized with a mixture containing ketamine/xylazine (300*µ*g each per g body-weight). Immediately after the euthanasia, the eye was enucleated and the connective tissues were removed. For transmission electron microscopy, the eyeball was fixed overnight at 4*°*C in 2% w/v paraformaldehyde, 2.5% v/v glutaraldehyde in 0.1M sodium cacodylate buffer, pH 7.4. After subsequent buffer washes, the samples were post-fixed in 2% w/v osmium tetroxide for 1 hour at room temperature, and rinsed in dH2O prior to en bloc staining with 2% uranyl acetate. After dehydration through a graded ethanol series, the tissue was infiltrated and embedded in EMbed-812 (Electron Microscopy Sciences, Fort Washington, PA). Thin sections were stained with uranyl acetate and lead citrate and examined with a JEOL 1010 electron microscope fitted with a Hamamatsu digital camera and AMT Advantage image capture software.

For the scanning electron microscopy, the eye was dissected as described in the “Rod recording”. The neural retina and the RPE were separated and small pieces of each were fixed overnight at 4*°*C in 2% w/v paraformaldehyde, 2.5% v/v glutaraldehyde in 0.1M sodium cacodylate buffer, pH 7.4. The samples were then dehydrated in a graded series of ethanol concentrations through 100% over a period of 1.5 hour. Dehydration in 100% ethanol was done three times. After 100% ethanol step dehydrated samples were incubated for 20min in 50% HMDS in ethanol followed by three changes of 100% HMDS (Sigma-Aldrich Co.) and followed by overnight air-drying as described previously [60]. Then samples were mounted on stubs and sputter coated with gold palladium. Specimens were observed and photographed using a Quanta 250 FEG scanning electron microscope (FEI, Hillsboro, OR, USA) at 10 kV accelerating voltage.

For immunofluorescence, the eyeball, with the cornea cut open, was fixed in 4% w/v paraformaldehyde for 60 minutes, rinsed in phosphate buffer (PB), cryoprotected overnight at 4*°*C in 0.1M PB containing 30% sucrose. The cornea and the lens were removed and the eyecup was embedded in a mixture of two parts 20% sucrose in PB and one part optimal cutting temperature compound (Tissue Tek, Electron Microscopy Sciences, Hateld, PA, USA) and frozen at -80*°* with the dorsal/superior side up. Radial sections (16*µ*m) were cut on a cryostat (Leica Biosystems, Buffalo Grove, IL, USA) and were collected on a superfrost plus glass slide (Fisher Scientific, Pittsburgh, PA, USA). Retinal cryosections were permeabilized in 0.1M PB containing 0.5% Triton X-100 and blocked in 0.1M PB containing 5% normal goat serum for 60 minutes at room temperature. For staining cryosections with antibodies against actin isoforms, the sections were first treated with 100% Methanol at -20*°*C before proceeding with the permeabilization. Sections were incubated with primary antibodies diluted in the blocking buffer at 4*°*C overnight, followed by three 10 minute washes with 0.1M PB. Sections were then incubated with fluorescently tagged secondary antibodies and/or fluorescently tagged Phalloidin (ThermoFisher Scientific, Waltham, MA) diluted in the blocking buffer at room temperature for 1 hour, followed by three 10 minute washes with 0.1M PB. Finally the slides were mounted using ProLong Glass mounting medium (ThermoFisher Scientific, Waltham, MA, USA) and imaged using a Nikon Ti microsopce fitted with a Yokogawa CSUX1 spinning disc confocal system. Using a 100X 1.49NA TIRF oil-immersion objective, images were captured on an Andor Ultra iXon 897 EM-CCD camera. Optical sectioning at 0.3*µ*m for 15 sections were used to produce a Z-stack.

Each pair of retinas from WT and Mut were dark-adapted overnight, collected at the same time of a day, and imaged in parallel under the same conditions and settings. Consecutive outer retinal images of immunostaining were taken along temporal, central, and nasal direction on each retinal cryosection. Maximum intensity projections of the Z-stack were used to generate a 2D XY image, and the intensity was averaged over the surrounding 1.7*µm×*1.7*µm* square. For antibodies which mark predominately RPE microvilli or photoreceptor OS (ezrin, myosin VIIa, transducin G*β*1, and rhodopsin), the top 960 intensities of each image were pooled to represent the average staining intensity for the retinal section. We also manually sampled 3 representative spots of each image to make sure that the automatic approach worked and to quantify staining intensities in the region of interest which were not predominately stained. Wildtype retinas stained with antibodies against actin isoforms were also imaged using a Lecia SP5 laser scanning confocal to produce a higher resolution image of actin isoform localization. To achieve this, a 100X 1.46NA oil-immersion objective was used and the pin-hole was closed to .8 Airy Units. Z-stacks were acquired and the images were deconvolved using Huygens software (Imaris) to obtained high-resolution images of the localization of actin isoforms in the retina.

### Antibodies

The following primary antibodies were used: mouse anti-*β*-actin (1:100 dilution, clone 4C2, MABT825, Millipore-Sigma), mouse anti-*γ*-actin (1:100 dilution, clone 2A3, MABT824, Millipore-Sigma), mouse anti-Ezrin (1:100 dilution, CPTC-ezrin-1, AB 2100318, Developmental Studies Hybridoma Bank, a gift from Dr. A. Bretscher, Cornell University), rabbit anti-p-ERM (1:100 dilution, a gift from Dr. A. Bretscher, Cornell University [50]), mouse anti-Myosin VIIa (1:50 dilution, clone C-5, sc-74516, Santa Cruz Biotechnology Inc.), rat anti-HCAM (CD44) (1:10 dilution, clone IM7, sc-18849, Santa Cruz Biotechnology Inc.), rabbit anti-G*β*1 (1:100 dilution, clone, C-16, sc-379, Santa Cruz Biotechnology Inc.), mouse anti-rhodopsin (1:100 dilution, clone 1D4, sc-57432, Santa Cruz Biotechnology Inc.).

### Mass spectrometry of retinal lysates and phalloidin pulldowns

Freshly excised mouse retinas from light adapted mice, with the cornea and the lens removed, were flash-frozen in liquid nitrogen and homogenized by grinding in liquid nitrogen with mortar and pestle. The powder was transferred into pre-chilled tube on ice and mixed with ice-cold lysis buffer (100mM HEPES pH 7.4, 50mM NaCl, 2mM MgCl2, 2mM ATP, and protease inhibitors cocktail (Sigma)) and passed through a 211/2 G syringe on ice. The lysate was clarified at 6,000 g, 10 min, 4*°*C. A portion of the lysate was mixed with Phalloidin-Biotin and incubated on a rocker for 2 hours at 4*°*C. Pulldowns were collected by addition of magnetic streptavidin beads, washed 3 times with lysis buffer, eluted into SDS sample buffer, and fractionated by a short run on SDS PAGE. The entire protein zone after Coomassie blue staining was excised and processed for mass spectrometry.

The mass spectrometry and its data analysis were performed as described previously [61]. The retinal protein samples of the supernatants and the corre-sponding pulldowns with phalloidin-Biotin were reduced with tris(2-carboxyethyl)-phosphine (TCEP), alkylated with iodoacetamide, and digested with trypsin. Tryptic digests were analyzed using a 1.5 h LC gradient on the Thermo Q Exactive Plus mass spectrometer. The raw data files from the samples described above were processed using MaxQuant software (Version 1.6.17.0) from Max-Planck Institute for Biochemistry, Martinsried, Germany [62], searching against the mouse protein reference UP000000589 and a locally compiled list of contaminants. The options of “Match between runs” and “Second peptide search” were enabled. Specific tryptic search was performed with the fixed modification of “Carbamidomethyl (C)” and the variable modifications of “Oxidation (M)”, “Acetyl (Protein N-term)”, “Deamidation (N)”, and “Phospho (STY)”. Consensus identification lists were generated with false discovery rates of 1% at protein, peptide and site levels. The locally compiled files of contaminants and search parameters were uploaded with the raw data files (ProteomeXchange Consortium accession number PXD031465).

This experiment was performed in triplicates, and proteins identified in the pulldowns were filtered against the total list of actin binding proteins [63], which was further narrowed down to manually exclude proteins that are not directly related to actin filament organization (Supplement 1). The iBAQ values of a protein of interest and the iBAQ values of *β* and *γ* actin were used as therelative amounts of the protein and the actin isoforms in the pulldown of each sample. Two methods were used to calculate p-values: (1) Mut and WT paired Student’s t-test of the iBAQ ratio of a protein to *β*- and *γ*-actin (n=3 pairs); (2) Spearman’s correlation coefficient of a protein iBAQ with *γ*-actin iBAQ (n=6 samples). A protein is considered to have a significant differential interaction with *β* and *γ* actin if a p-value *≤* 0.05 (from either calculations), the fold change *log*2(Mut/WT) *≥* 0.5, the fold change of the pulldown relative to the input *≥* 0.5, and the MS/MS count *≥* 9 in either Mut or WT.

### In vivo imaging of the retina

Mice were adapted for at least one week of 7am/7pm light/dark cycle. On the day of imaging, mice were taken directly from the housing facility, light adapted to the room light or further dark adapted after 7am for another 3 to 8 hours in their cage (with food and water), which was placed in a specially aerated black box; under room light or dim red light, respectively, their pupils were dilated with 1% tropicamide (Mydriacyl, Alconox, New York, NY, USA) and they were anesthetized by intraperitoneal injection of a mixture containing ketamine/xylazine/acepromazine (90*µ*g/10*µ*g/2*µ*g per g body-weight) and placed onto a platform maintained at 38*°*C. Depth of anesthesia was monitored by toe pinching and eye reflex. Spectral domain optical coherence tomography (OCT) and confocal scanning laser ophthalmoscope (SLO) were were performed in sequence. OCT was performed using a Bioptigen Envisu (R2200; Bioptigen Inc.) coupled to a broadband LED light source (T870-HP; Superlum Diodes, Ltd.). SLO was performed using a Spectralis HRA (Heidelberg Engineering) with blue AF (488 nm excitation) imaging mode. At the end of the imaging session, animals were returned to the cage on a body-temperature heating pad, observed until they fully recovered, and then brought back to the facility. These animals were checked daily and stayed in 7am/7pm light/dark cycle of the facility for a week before an additional imaging session with dark/light adaptation condition reversed for each mouse. Data analysis was performed with custom programs (MATLAB software, MathWorks, Natick, MA, USA). OCT images were automatically segmented, with manual verification, into reflective bands; the location and the intensity of each band were averaged along the line of the best linear fit of a section of the band between 280 *µm* and 420 *µm* away from the center of the optic disk. SLO images (768*×*768 pixels and 105*°* degrees of view) were filtered with a balanced positive center and negative surround filter with the center radius of 5 pixels (*∼*20*µm*, given 55-65 pixels of the imaged optic disks which presumably were 0.2-0.25 mm in diameter [64]) and the surround 3 times of that. The filtered images were then thresholded at zero and the number of the local maximum (visually corresponding to the autofluorescent spots) were counted.

### Electroretinograph

The electroretinograph (ERG) recording were performed as described previously [26, 27]. Briefly, mice were dark adapted for more than 12 hours. Before ERG, mice were anesthetized under dim red light by intraperitoneal injection of a mixture containing ketamine/xylazine/acepromazine (90*µ*g/10*µ*g/2*µ*g per g body-weight) and were placed on a platform maintained at 38*°*C. Pupils were dilated with 1% tropicamide saline solution. A platinum electrode was inserted into the mouth to serve as the reference and the ground electrode, another platinum electrode was placed on the cornea of each eye, and the platform was moved inside a light-proof Ganzfeld Faraday cage during ERG. ERG recordings from Mut and their WT littermates were performed on the consecutive hours of the same day or the same hour of the consecutive days under the same settings and conditions. Light stimuli were either 4 ms flashes produced by a green light-emitting diode (LED) or *<*1 ms flashes produced by a Xenon tube delivered in the Ganzfeld (Espion Electrophysiology System; Diagnosys, Lowell, MA, USA).

### Rod recording

Due to the elongated microvilli of mutant RPE, which extended further along rod outer segments, it was difficult to have clean separation between their RPE and neural retina. We used three methods to record photoresponses under rod stimulating light intensities in the experimental order: (1) suction pipette outer-segment-in, (2) suction pipette inner-segment-in, and (3) multielectrode array (MEA), which were performed as described previously [27], [65], and [66], respectively, with some modifications. Before each experiment, mice were dark adapted for more than 12 hours; they were then intraperitoneally injected with a lethal dose of ketamine/xylazine mixture (300*µ*g each per g body-weight) under dim red light. Eyes were quickly enucleated and dissected under infrared light or dim red light in a Petri dish filled with Lockes solution and bovine serum albumin. Small pieces of retina were transferred to a Petri dish with constant perfusion of oxygenated (95% O2/5% CO2) Ames media for MEA recording. Additional small pieces of retina were excised, sliced, and transferred to a chamber for suction pipette recording. The chamber was perfused with Lockes solution (pH 7.4, no BSA added) and maintained at 35-37*°*C with a heating system designed for microscopy (Warner Instruments, Hamden, CT, USA). The Lockes solution was continuously bubbled with a gas mixture containing 95% O2/5% CO2 before it was directed into the chamber. Freshly pulled glass pipettes with a tip internal diameter of 4 *µ*m were used to suck in 1 to 3 rod outer segments (outer-segment-in) or several photoreceptor cell bodies/inner segments (inner-segment-in) for recording. The pipette, pipette holder, and a tube connecting pipette holder to the pressure control system were filled with Lockes solution. A small air bubble separated Lockes solution from the oil-filled pressure control system. For MEA recordings, the retina was placed ganglion cell side down in the recording chamber (pMEA 100/30iR-Tpr; Multi Channel Systems) of a 60-channel MEA system with a peristaltic pump. A custom-made dialysis membrane weight was placed on the retina, adding positive pressure from above. Additionally, a vacuum was applied to the retina using the constant vacuum pump, adding negative pressure and improving electrode-to-tissue contact and the signal-to-noise ratio. During recording, constant perfusion of oxygenated Ames media (34 *°*C) was provided to the recording chamber. Light stimuli were 5 ms flashes produced by the green LEDs in the recording chambers, which were calibrated before the experiments with a calibrated photodiode (OSI Optoelectronics, Hawthorne, CA, USA). The light intensity was converted to photoisomerizations per rod using 0.5 and 1.0 *µ*m^2^ collecting area per rod for suction pipette recording and MEA recording, due to the perpendicular and parallel stimulating light paths to OS, respectively.

### Analysis of response sensitivities

The ERG B-wave response sensitivity *S* and the maximum amplitude *Bmax* were calculated by fitting the B-wave amplitudes in response to four light flashes of different intensities using the following equation:

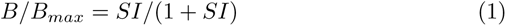

in which *B* is the amplitude of B-wave response to a light flash of intensity *I* and *S* is the inverse of half saturating intensity, *I*0.5, e.g., in [67].

The ERG A-wave sensitivity *K* and the maximum amplitude *Amax* were calculated by fitting A-wave rising phase time courses, *A*(*I, t*), using the equation of [30]:

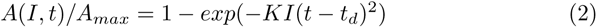

in which *t_d_* is a constant time delay. For each flash of intensity *I*, the rising phase of the corresponding A-wave was chosen to be the time period from *t_d_* after the light flash until reaching the maximum (or 5 msec before—if the period is longer than 5 msec). The value of *t_d_* was fixed at 3.7 msec.

The rod photocurrent sensitivity was calculated by fitting the same equation as ERG A-wave, except that the rising phase was chosen to be fixed at 60 msec (flashes used in these experiments were much dimmer than ERG) and the value of *td* was fitted for each recording, varying between 14 to 17 msec due to small variations of capacitance and/or conductance between the rods and the suction pipettes. The values of *SA* and *Amax* were averaged over the 3 to 6 recorded rods for each mouse. The MEA field potential sensitivity was calculated similarly, except that the rising phase was chosen to be fixed at 30 msec (after that the recording had ganglion cell spikes superimposed on rod response traces) and the value of *td* was 0.

For the numerical analysis, the original response data were digitally filtered by a 6th order Butterworth filter with a cutoff frequency of 22.5*Hz*, 1*kHz*, and 22.5*Hz* to remove high frequency components in B-wave, A-wave, and rod recording, respectively.

### Experimental design and statistical analysis

Most of the experiments were performed in Mut and WT pairs, for which the p-values were calculated from paired Student’s t-test on the *n*-number of experiments of Mut/WT pairs. For Pearson/Spearman correlation coefficients, the p-values were calculated by transforming the correlation between *n* pairs of concerned variables to create a t-statistic having *n −* 2 degrees of freedom. Everything was computed using MATLAB software.

For experiments not performed in pairs, the p-values were calculated from Welch’s t-test by

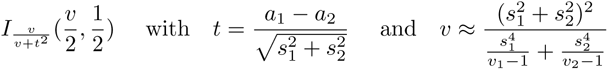

where I is the regularized incomplete beta function with the statistic t and the degrees of freedom *v* in which *a*_1_, *s*_1_, and *v*_1_ are the mean, the SEM, and the size of the 1*^st^* sample, and *a*_2_, *s*_2_, and *v*_2_ of the 2*^nd^* sample. This can be generalized to any linear combination of independent random variables [68, 69], following the formulation of [68]:

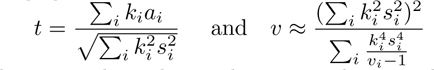

in which k_i_ is a known real number and a_i_, s_i_, and v_i_ are the mean, the SEM, and the sample size of the *i^th^* variable. We used this as the “Welch’s t-test” in the quantification of the OS diameter and volume as well as the transducin concentration in the OS, with all variables in *log*() expression.

## Supporting information

Vedula et al 2023 selected list of actin binding proteins

## Acknowledgments

This work was supported by NIH/NIGMS R35GM122505, NIH/NINDS R01NS102435, NIH/NEI P30EY001583, NIH/ORIP S10OD026860, F.M. Kirby Foundation, the Office of Dean at School of Veterinary Medicines and the Office of the Vice Provost for Research at the University of Pennsylvania. We thank Dr. Noga Vardi for helpful discussions. We thank Stephanie Sterling and Nicolae A Leu for help with mouse colony maintenance and breeding. We thank Dr. Gordon Ruthel and the PennVet Imaging core for laser scanning confocal imaging. We thank Inna Martynyuk and Dr. Yuri Veklich and the Microscopy Core in the Department of Cell and Developmental Biology for TEM and SEM imaging. We thank the Bollinger Digital Fabrication Lab at the University of Pennsylvania Library Biotech Commons for 3D printing imaging apparatus. We thank Dr. Hsin Yao Tang and the Wistar Proteomics Core facility for mass spectrometry. We thank Dr. Noga Vardi at University of Pennsylvania for donating G*β*1 antibody and Dr. Anthony Bretscher and Dr. Anthony Lombardo at Cornell University for donating Ezrin and p-ERM antibodies used in this study.

---

**Figure S1.**
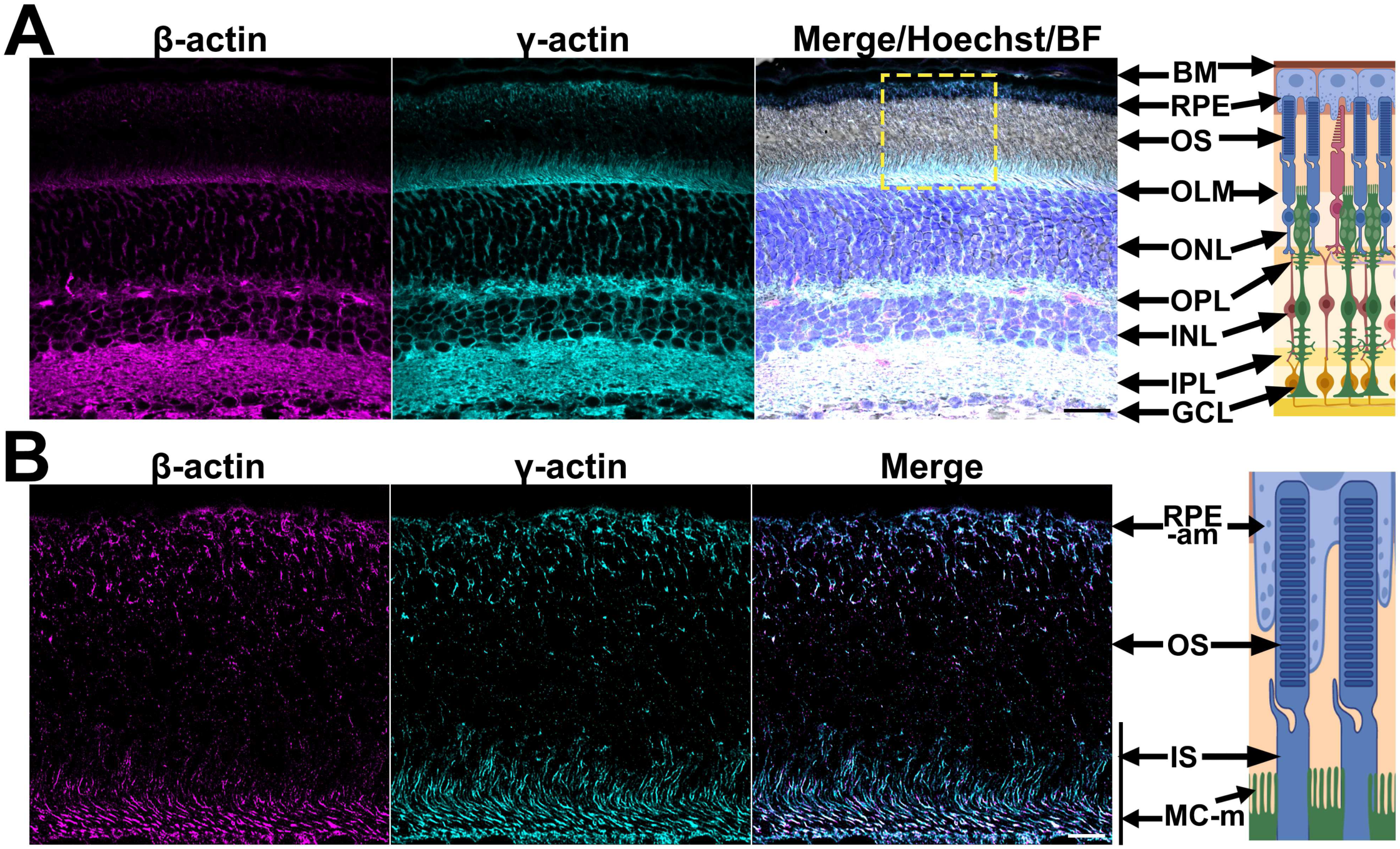
(A) Same as Figure 1A, the localization of non-muscle actin isoforms in the wildtype retina. Scale bar: 20*µ*m. Pink: *β*-actin. Cyan: *γ*-actin. Blue: DAPI. BM: Bruch’s membrane. RPE: retinal pigment epithelium. OS/IS: outer/inner segment. OLM: outer limiting membrane. ONL/INL: outer/inner nuclear layer. OPL/IPL: outer/inner plexiform layer. GCL: ganglion cell layer. (B) The selected region of the retina from (A) containing both the RPE and Müller cell microvilli was imaged with a laser scanning confocal setting the pinhole to less than 1 airy unit and by carrying out deconvolution to obtain a super-resolution image. *β*- and *γ*-actin were co-localized in the cellular protrusions of the RPE intersecting the OS, as well as in some of the processes where the Müller cells intersect the IS. Scale bar: 5*µ*m. RPE-am: RPE apical microvilli. MC-m: Müller cell microvilli.

**Figure S2.**
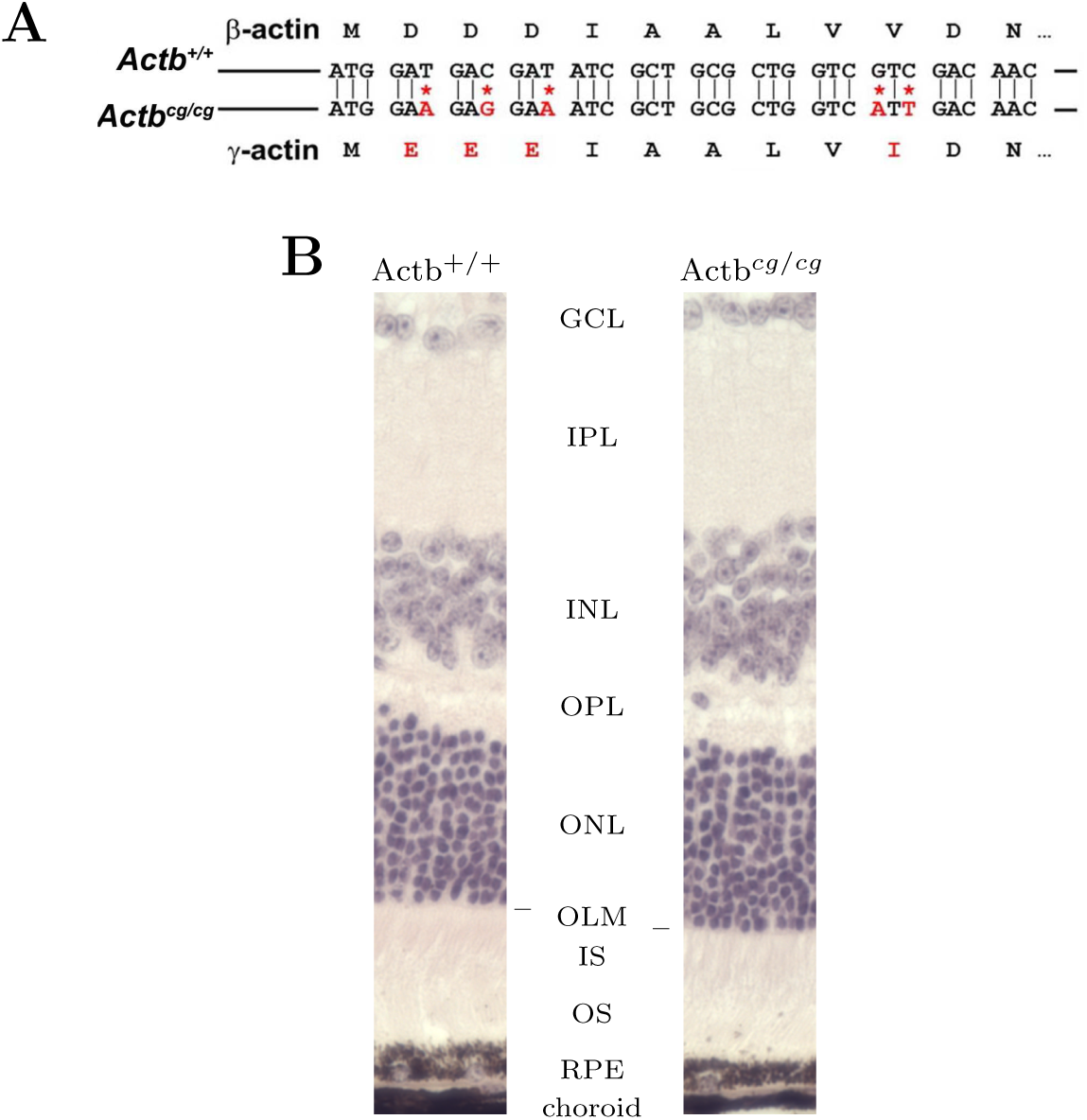
(A) Actb*^cg/cg^* is generated with editing 5 Actb nuclei acids to produce *γ* instead of *β* actin protein. (B) Retina layers of the Actb*^cg/cg^* mouse look the same as the wildtype.

**Figure S3.**
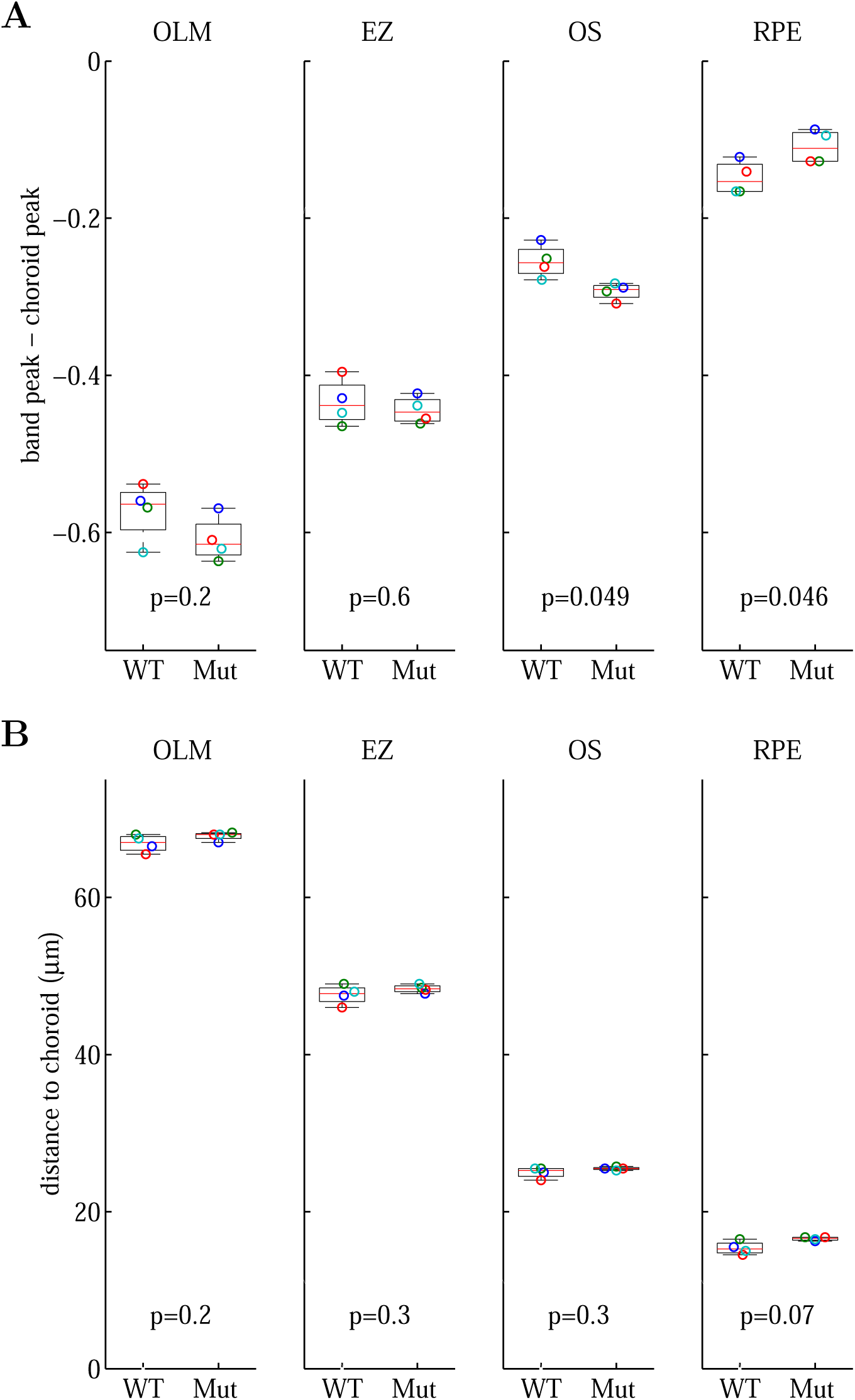
OCT bands relative to the choroid band. (A) Contrast. (B) Distance.

**Figure S4.**
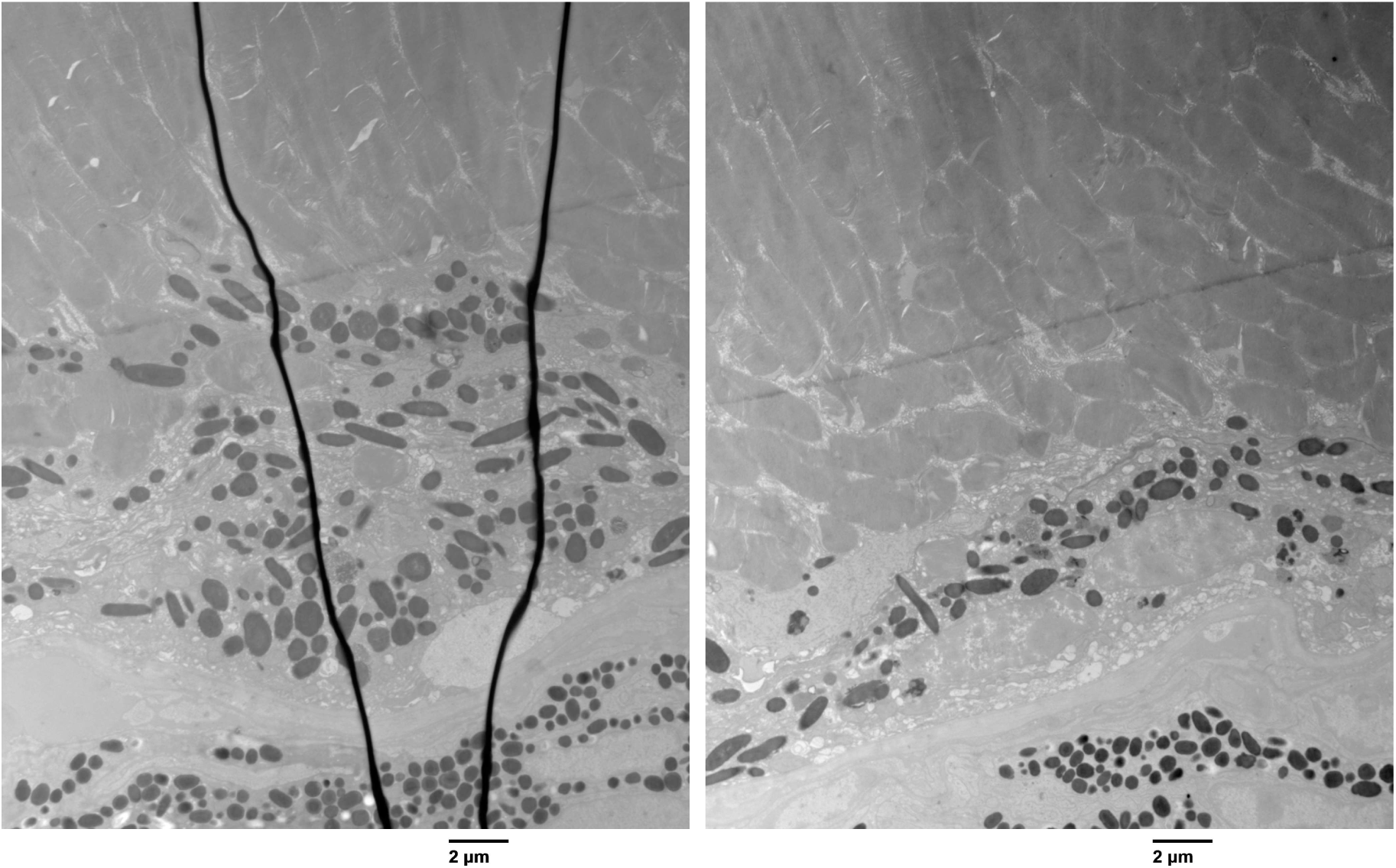
The high resolution TEM images containing the microglia cell shown in Figure 3d.

**Figure S5.**
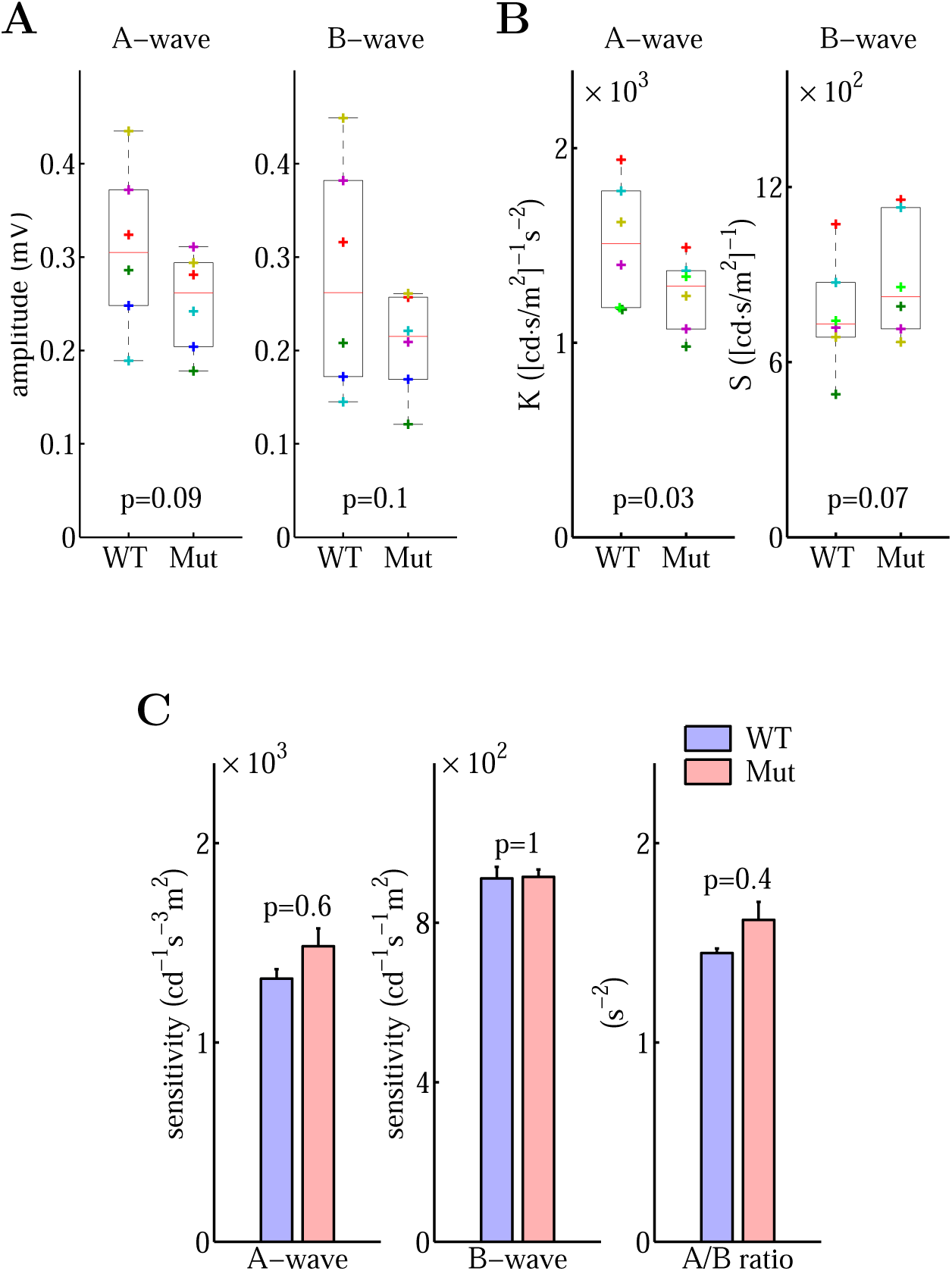
(A) Response amplitudes of ERG A-wave and B-wave. (B) Response sensitivities of ERG A-wave and B-wave. (C) Response sensitivities of ERG A-wave, B-wave, and their ratio for yound mice aged between 3 to 4 months. WT: Actb^+*/*+^. Mut: Actb*^cg/cg^* . The error bars represent SEM. The p-values are from paired Student’s t-test with *n* = 6.

**Figure S6.**
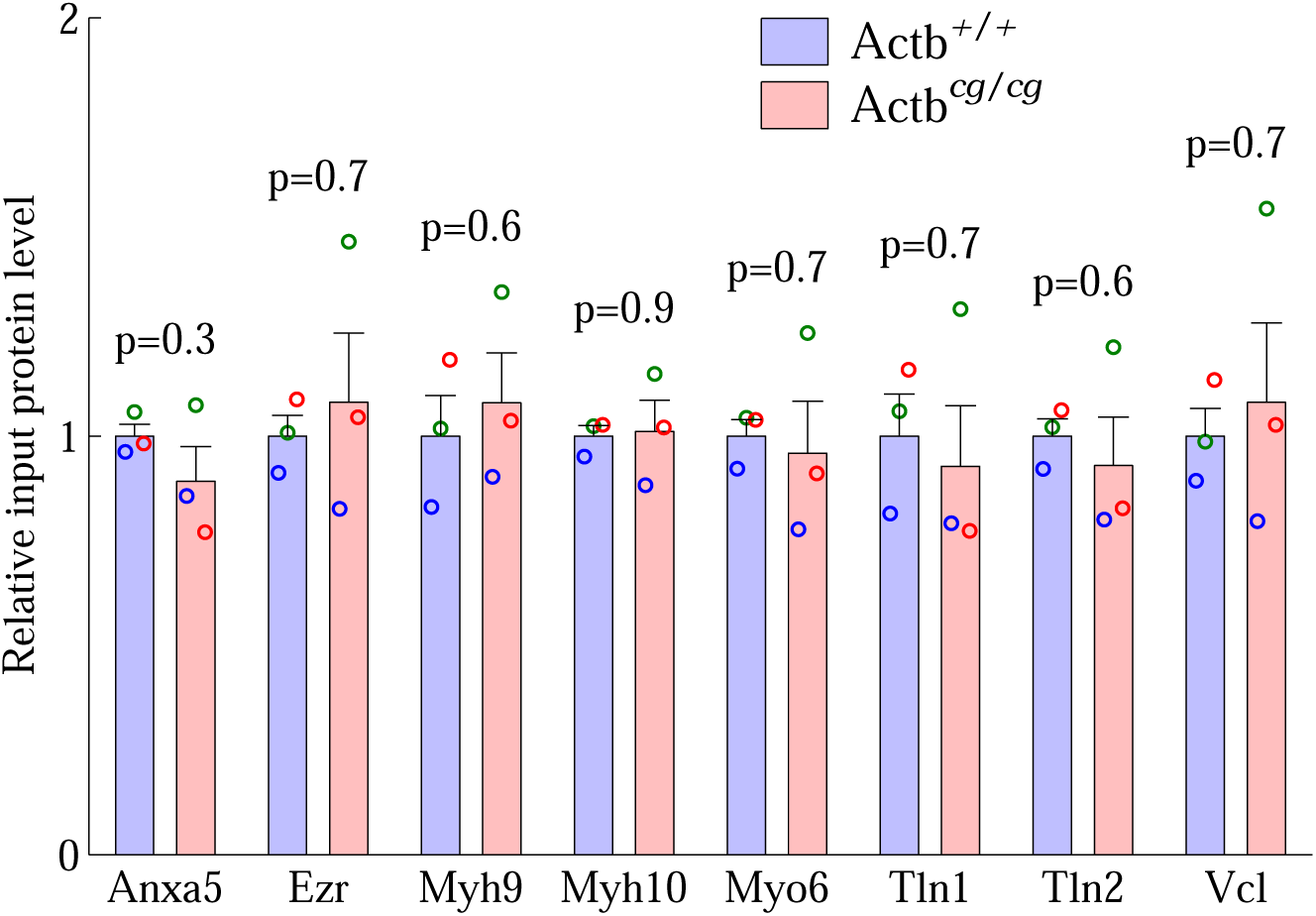
The relative levels of actin-binding proteins did not differ between the Actb*^cg/cg^* and the wildtype retinal lysates. The LFQ (load-free quantification) value of each protein quantified from the mass-spec data was used as the protein level. The error bars represent SEM. The p-values are from paired Student’s t-test with *n* = 3.

