## Supplementary material for "*β*-actin is essential for structural integrity and physiological function of the retina": Vedula et al 2023 selected list of actin binding proteins

Vedula et al 2023 selected list of actin binding proteins (adapted from Gao and Nakamua 2022)

| Protein name | Gene name | Chromosome | Size, aa | Binding mode(s)* | Function | Disease | Actin Binding domain | Human Protein Atlas |
|---|---|---|---|---|---|---|---|---|
| Unconventional myosin-Ia, Brush border myosin I (BBM1), Myosin I heavy chain (MIHC) | MYO1A, MYHL | 12 | 1043 | Motor | Regulate movement of organelles along actin filaments. | Deafness. | ABD (571-593) within motor domain | https://www.protein |
| Unconventional myosin-Ib, MYH-1c, Myosin I alpha (MMIa) | MYO1B | 2 | 1136 | Motor | Regulate cell migration, neurite outgrowth and vesicular transport. | Expression is increased in cancers. | ABD (578-600) within motor domain | https://www.protein |
| Unconventional myosin-Ic, Myosin I beta (MMIb) | MYO1C | 17 | 1063 | Motor | Regulate transforming growth factor-β-signaling and fibrosis in podocytes. | Impaired visual function. | ABD (608-630) within motor domain | https://www.protein |
| Unconventional myosin-Id | MYO1D, KIAA0727 | 17 | 1006 | Motor | Regulate endosomal protein trafficking. | Regulate regeneration process after demyelination. Laterality defects. | ABD (572-594) within motor domain | https://www.protein |
| Unconventional myosin-Ie, Myosin-Ic | MYO1E, MYO1C | 15 | 1108 | Motor | Required for normal kidney function. | Focal segmental glomerulosclerosis. | ABD (581-591) within motor domain | https://www.protein |
| Unconventional myosin-If, Myosin -Ie | MYO1F | 19 | 1098 | Motor | Regulate neutrophil migration and immune signaling. | Thyroid cancer. | ABD (579-589) within motor domain | https://www.protein |
| Unconventional myosin-Ig, Minor histocompatibility antigen HA-2 (mHag-HA-2) | MYO1G, HA2 | 7 | 1018 | Motor | Regulate T-cell migration. | Acute Lymphoblastic Leukemia. | ABD (584-606) within motor domain | https://www.protein |
| Unconventional myosin-Ih, Myosin-1H | MYO1H | 12 | 1032 | Motor | Regulate $CO_2$ sensitivity and respiratory control. | Congenital central hypoventilation. | ABD (578-600) within motor domain | https://www.protein |
| Myosin-1, Myosin heavy chain 1, Myosin heavy chain 2x (MyHC-2x), Myosin heavy chain IIx/d (MyHC-IIx/d), Myosin heavy chain, skeletal muscle, adult 1 | MYH1 | 17 | 1939 | Motor | Muscle contraction. | Myopathies. | ABD (659-681, 761-775) within motor domain | https://www.protein |

| | | | | | | | | |
|---|---|---|---|---|---|---|---|---|
| Myosin-2, Myosin heavy chain 2, yosin heavy chain 2a (MyHC-2a, MyHC-IIa), Myosin heavy chain, skeletal muscle, adult 2 | MYH2, MYHSA2 | 17 | 1941 | Motor | Muscle contraction. | Myopathies. | ABD (661-683, 763-777) within motor domain | https://www.protein |
| Myosin-3, Muscle embryonic myosin heavy chain, Myosin heavy chain 3, Myosin heavy chain, fast skeletal muscle, embryonic, SMHCE | MYH3 | 17 | 1940 | Motor | Muscle contraction. | Myopathies. Freeman-Sheldon syndrome and Sheldon-Hall syndrome. | ABD (656-678, 758-772) within motor domain | https://www.protein |
| Myosin-4, Myosin heavy chain 2b (MyHC-2b), Myosin heavy chain, skeletal muscle, fetal | MYH4 | 17 | 1939 | Motor | Muscle contraction. | Myopathies. | ABD (659-681, 761-775) within motor domain | https://www.protein |
| Myosin-6, Myosin heavy chain 6, Myosin heavy chain, cardiac muscle alpha isoform (MyHC-alpha) | MYH6, MYHCA | 14 | 1939 | Motor | Muscle contraction. | Cardiomyopathy. Sick sinus syndrome. | ABD (657-679, 759-773) within motor domain | https://www.protein |
| Myosin-7, Myosin heavy chain 7, Myosin heavy chain slow isoform (MyHC-slow), Myosin heavy chain, cardiac muscle beta (MyHC-beta) | MYH7, MYHCB | 14 | 1935 | Motor | Muscle contraction. | Cardiomyopathy. | ABD (655-677, 757-771) within motor domain | https://www.protein |
| Myosin-7B, Antigen MLAA-21, Myosin cardiac muscle beta chain, Myosin heavy chain 7B, cardiac muscle beta isoform, Slow A MYH14 | MYH7B, KIAA1512 | 20 | 1983 | Motor | Muscle contraction. | Cardiomyopathy. | ABD (704-726, 806-820) within motor domain | https://www.protein |

| | | | | | | | | |
|---|---|---|---|---|---|---|---|---|
| Myosin-8, Myosin heavy chain 8, Myosin heavy chain, skeletal muscle, perinatal (MyHC-perinatal) | MYH8 | 17 | 1937 | Motor | Muscle contraction. | Myopathies. | ABD (658-680, 760-774) within motor domain | https://www.protein |
| Myosin-9, Myosin heavy chain 9, Cellular myosin heavy chain, type A, Myosin heavy chain, non-muscle IIa, Non-muscle myosin heavy chain A (NMMHC-A), Non-muscle myosin heavy chain IIa (NMMHC II-a) | MYH9 | 22 | 1960 | Motor | Regulate cytokinesis, cell shape, and migration. | Alport syndrome, Cataract, Deafness. | ABD (654-676) within motor domain | https://www.protein |
| Myosin-10, Cellular myosin heavy chain, type B, Myosin heavy chain 10, Myosin heavy chain, non-muscle IIb, Non-muscle myosin heavy chain B (NMMHC-B), Non-muscle myosin heavy chain IIb (NMMHC II-b) | MYH10 | 17 | 1976 | Motor | Regulate cytokinesis, cell shape, and migration. | Intellectual disability, microcephaly, and feeding difficulties as well as cerebral atrophy. | ABD (661-683) within motor domain | https://www.protein |
| Myosin-11, Myosin heavy chain 11, Myosin heavy chain, smooth muscle isoform, SMMHC | MYH11, KIAA0866 | 16 | 1972 | Motor | Muscle contraction. | Acute myeloid leukemia, aortic aneurysm. | ABD (661-683, 762-776) within motor domain | https://www.protein |

| | | | | | | | | |
|---|---|---|---|---|---|---|---|---|
| Myosin-13, Myosin heavy chain 13, Myosin heavy chain, skeletal muscle, extraocular (MyHC-EO), Myosin heavy chain, skeletal muscle, laryngeal (MyHC-IIL), Superfast myosin | MYH13 | 17 | 1938 | Motor | Muscle contraction. | | ABD (659-681, 761-775) within motor domain | https://www.protein |
| Myosin-14, Myosin heavy chain 14, Myosin heavy chain, non-muscle IIc, Non-muscle myosin heavy chain IIc (NMHC II-C) | MYH14, KIAA2034, FP17425 | 19 | 1995 | Motor | Regulate cytokinesis, cell shape, and migration. | Deafness, neuropathy. | ABD (678-700) within motor domain | https://www.protein |
| Myosin-15, Myosin heavy chain 15 | MYH15, KIAA1000 | 3 | 1946 | Motor | Muscle contraction. | Deafness. Amyotrophic lateral sclerosis. | ABD (667-689, 769-783) within motor domain | https://www.protein |
| Myosin-IIIa | MYO3A | 10 | 1616 | Motor | Regulate auditory hair bundle. | Deafness. Bardet-Biedl syndrome. | ABD (934-956) within motor domain | https://www.protein |
| Myosin-IIIb | MYO3B | 2 | 1341 | Motor | Regulate auditory hair bundle. | Deafness. Bardet-Biedl syndrome. | ABD (939-961) within motor domain | https://www.protein |
| Unconventional myosin-Va, Dilute myosin heavy chain, non-muscle, Myosin heavy chain 12, Myosin-12, Myoxin | MYO5A, MYH12 | 15 | 1855 | Motor | Transport melanosome and vesicles. | Griscelli syndrome type 1 and Elejalde syndrome. | ABD (643-665) within motor domain | https://www.protein |
| Unconventional myosin-Vb | MYO5B, KIAA1119 | 18 | 1848 | Motor | Regulate vesicle transport and trafficking. | Microvillus inclusion disease. | ABD (640-662) within motor domain | https://www.protein |
| Unconventional myosin-Vc | MYO5C | 15 | 1742 | Motor | Regulate transferrin trafficking. | Diabetic retinopathy. | ABD (632-654) within motor domain | https://www.protein |
| Unconventional myosin-VI | MYO6, KIAA0389 | 6 | 1294 | Motor | Reverse-direction motor protein that moves towards the minus-end of actin filament. Vesicular membrane trafficking and cell migration. | Deafness. | ABD (665-672) within motor domain | https://www.protein |
| Unconventional myosin-VIIa | MYO7A, USH1B | 11 | 2215 | Motor | Regulate migration of retinal pigment epithelial melanosomes and phagosomes. | Usher syndrome type III (USH1B). | ABD (632-639) within motor domain | https://www.protein |

| | | | | | | | | |
|---|---|---|---|---|---|---|---|---|
| Unconventional myosin-VIIb | MYO7B | 2 | 2116 | Motor | Regulate microvilli organization. | | ABD (637-659) within motor domain | https://www.protein |
| Unconventional myosin-IXa | MYO9A, MYR7 | 15 | 2548 | Motor | Regulate motor neuron axon guidance. | Bardet-Biedl Syndrome. Congenital myasthenic syndrom. | ABD (898-920) within motor domain | https://www.protein |
| Unconventional myosin-IXb | MYO9B, MYR5 | 19 | 2157 | Motor | Regulate cell migration through RHOA. | Celiac disease, lcerative colitis and Crohn's disease. | ABD (844-855) within motor domain | https://www.protein |
| Unconventional myosin-X | MYO10, KIAA0799 | 5 | 2058 | Motor | Mediatescargo transport. Regulate cell shape, cell spreading and cell adhesion. | Cancer metastasis and pathogen infection. | ABD (619-641) within motor domain | https://www.protein |
| Unconventional myosin-XV, Unconventional myosin-15 | MYO15A, MYO15 | 17 | 3530 | Motor | Transport vesicle. | Deafness | ABD (1792-1799) within motor domain | https://www.protein |
| Unconventional myosin-XVI, Neuronal tyrosine-phosphorylated phosphoinositide-3-kinase adapter 3 | MYO16, KIAA0865, NYAP3, MYO16B | 13 | 1858 | Motor | Transport vesicle. | Neurological disorders. | Motor domain (401-1145) | https://www.protein |
| Unconventional myosin-XVIIIa, Molecule associated with JAK3 N-terminus (MAJN), Myosin containing a PDZ domain, Surfactant protein receptor SP-R210 (SP-R210) | MYO18A, CD245, KIAA0216, MYSPDZ | 17 | 2054 | Motor | Intracellular trafficking. | Myopathies. | Motor domain (405-1185) | https://www.protein |
| Unconventional myosin-XVIIIb | MYO18B | 22 | 2567 | Motor | Intracellular trafficking.Regulate tumor development and progression. | Myopathies. | Motor domain (571-1333) within motor domain | https://www.protein |
| Unconventional myosin-XIX, Myosin head domain-containing protein 1 | MYO19, MYOHD1 | 17 | 970 | Motor | Regulate mitochondrial transport or positioning. | | ABD (602-624) within motor domain | https://www.protein |

| | | | | | | | | |
|---|---|---|---|---|---|---|---|---|
| Myosin light chain kinase, smooth muscle (MLCK, smMLCK), Kinase-related protein (KRP), Telokin | MYLK, MLCK, MLCK1, MYLK1 | 3 | 1914 | | Regulate smooth muscle contraction via phosphorylation of myosin light chains. | Aortic aneurysm, familial thoracic 7 (AAT7), Megacystis-microcolon-intestinal hypoperistalsis syndrome (MMIHS) | DFRxxL motifs of N-terminal | https://www.protein |
| Formin-1, Limb deformity protein homolog | FMN1, FMN, LD | 15 | 1419 | Nucleation, polymerization, bundling, Severing | Cell movement, shape change | X-linked Alport syndrome, autism spectrum disorder, schizophrenia, Cenani–Lenz-like non-syndromic, hearing loss, congenital anomalies of the kidney and urinary, melanoma, pancreatic, prostate cancer. | FH2<br>PDB: 1UX5 | https://www.protein |
| Formin-2 | FMN2 | 1 | 1722 | Nucleation, polymerization, bundling | Cell movement, shape change | Intellectual disability and neurodevelopmental disorders, premature ovarian failure and infertility, Alzheimer disease, colorectal, brain, pancreatic, glioma and testicular cancer. | FH2 | https://www.protein |
| Formin-like protein 1, CLL-associated antigen KW-13, Leukocyte formin | FMNL1, C17orf1, C17orf1B, FMNL, FRL1 | 17 | 1100 | Nucleation, polymerization, severing | Cell movement, shape change | Osteocarcinoma, lymphoma, leukemia, brain, head and neck, lung, ovarian, pancreatic, gastric cancer, renal carcinoma. | FH2 | https://www.protein |

| | | | | | | | | |
|---|---|---|---|---|---|---|---|---|
| Formin-like protein 2, Formin homology 2 domain-containing protein 2 | FMNL2, FHOD2, KIAA1902 | 2 | 1086 | Nucleation, polymerization | Cell movement, shape change, Proliferation, regulate Golgi architecture. | Adenomyosis, mental retardation, POEMS syndrome, Crohn’s disease, brain, colorectal, ovarian, gastric cancer, gallbladder carcinoma, hepatocellular carcinoma, melanoma, early-onset stroke, glaucoma. | FH2 | https://www.protei |
| Formin-like protein 3, Formin homology 2 domain-containing protein 3, WW domain-binding protein 3 | FMNL3, FHOD3, FRL2, KIAA2014, WBP3 | 12 | 1028 | Nucleation, polymerization | Cell movement, shape change, Proliferation, regulate Golgi architecture. | Gastric, pancreatic, colorectal, head and neck, ovarian, prostate cancer, dermatophyma, nasopharyngeal carcinoma, peripheral neurodegenerative disorder. | FH2 | https://www.protei |
| FH1/FH2 domain-containing protein 1, Formin homolog overexpressed in spleen 1, Formin homology 2 domain-containing protein 1 | FHOD1, FHOS, FHOS1 | 16 | 1164 | Polymerization | Organize actin filaments | Gastric, brain, breast, head and neck, renal cancer, dermatophyma, lung, oral squamous carcinoma, impaired cardiac. | FH2 | https://www.protei |
| FH1/FH2 domain-containing protein 3, Formactin-2, Formin homolog overexpressed in spleen 2 | FHOD3, FHOS2, KIAA1695 | 18 | 1422 | Polymerization | Organize actin filaments | Brain, ovarian, thyroid cancer, leukemia, hypertrophic and dilated cardiomyopathies, type I diabetes, periodontal disease. | FH2 | https://www.protei |
| Disheveled-associated activator of morphogenesis 1 | DAAM1, KIAA0666 | 14 | 1078 | Nucleation, polymerization | Cell movement, shape change | Brain, breast, colorectal, esophagus, osteocarcinoma, dermatophyma, congenital heart defects, anomalies of the kidney and urinary tract, cerebral palsy. | FH2 PDB: 2Z6E | https://www.protei |

| | | | | | | | | |
|---|---|---|---|---|---|---|---|---|
| Dishevelled associated activator of morphogenesis 2 | DAAM2, KIAA0381 | 6 | 1068 | Nucleation, polymerization | Cell movement, shape change | Brain, Cervix, colorectal, lung cancer, renal carcinoma, hepatocellular carcinoma, schizophrenia, Guillain–Barre syndrome, diffuse pulmonary ossification, diabetic nephropathy, osteoporosis, melanoma. | FH2 | https://www.protein |
| Protein diaphanous homolog 1, Diaphanous-related formin-1, mammalian Diaphanous-related formin (mDia1) | DIAPH1, DIAP1 | 5 | 1272 | Nucleation | Actin polymerization. Regulate spindle formation and cell division. | Deafness, autosomal dominant 1 (DFNA1), Seizures, cortical blindness, and microcephaly syndrome (SCBMS), neuroinflammation and neurodegenerative diseases, brain, breast, colorectal, esophagus, head and neck, ovarian cancer, glioma, leukemia, dermatophyma, Mendelian disorder, thrombocytopenia, Alzheimer disease, diabetes-associated nephropathy, ischemic stroke, Moyamoya disease. | FH2, FH1-FH2 domain | https://www.protein |
| Protein diaphanous homolog 2, Diaphanous-related formin-2 | DIAPH2, DIA | X | 1101 | Nucleation | Actin polymerization. Regulate formation of lamellipodia and filopodia. | Premature ovarian failure 2A, breast, colorectal, head and neck, renal, lung cancer; age-related macular degeneration, premature ovarian failure. | FH2 | https://www.protein |

| | | | | | | | | |
|---|---|---|---|---|---|---|---|---|
| Protein diaphanous homolog 3, Diaphanous-related formin-3 | DIAPH3, DIAP3 | 13 | 1193 | Nucleation | Actin polymerization. Regulate cytokinesis and microtubule dynamics. | Hepatocellular carcinoma, pancreatic, brain, breast, colorectal, head and neck, lung, ovarian, prostate, auditory neuropathy spectrum disorders, neurodevelopmental disorders, infantile spasm, epileptic encephalopathy. | FH2 | https://www.protein |
| Inverted formin-2, HBEBP2-binding protein C | INF2, C14orf151, C14orf173 | 14 | 1249 | Severing | Essential for proper placentation. Accelerates actin polymerization and depolymerization. | Neurodegeneration, Neuropathy, Focal segmental glomeruloesclerosis (FSGS), Charcot–Marie–Tooth (CMT) disease, intellectual disability and neurodevelopmental disorders, brain, breast, prostate, colorectal, thyoid cancer, thrombocytopenia, auditory nerve damage, Focal segmental glomerulosclerosis (FSGS), steroid-resistant nephrotic syndrome. | WH2, FH2 | https://www.protein |

| | | | | | | | | |
|---|---|---|---|---|---|---|---|---|
| Protein cordon-bleu | COBL, KIAA0633 | 7 | 1261 | Nucleation | Crucial for neuromorphogenesis processes | Acute lymphoblastic leukemia, Alzheimer's disease, pulmonary neuroendocrine carcinomas, lung cancer, lung squamous cell carcinoma, type 1 diabetes, head and neck squamous cell carcinoma, Huntington's disease, renal cell carcinoma. | Three WH2 PDB: 4JHD | https://www.protein |
| Junction-mediating and -regulatory protein | JMY | 5 | 988 | Nucleation | A regulator of both transcription and actin polymerization | Cardiovascular Disease, ankylosing spondylitis, glioblastoma, pancreatic cancer. | Tandem WH2 | https://www.protein |
| Leiomodin-1, 64 kDa autoantigen 1D, 64 kDa autoantigen 1D3, 64 kDa autoantigen D1, Leiomodin, muscle form, Smooth muscle leiomodin (SM-Lmod), Thyroid-associated ophthalmopathy autoantigen | LMOD1 | 1 | 600 | Nucleation | Bind along F-actin, next to the pointed end. Involved in sarcomere assembly and organization. | Visceral myopathy, megacystis microcolon intestinal hypoperistalsis syndrome. | ABS1, ABS2 (LRR domain), ABS3 (WH2 domain) | https://www.protein |
| Leiomodin-2, Cardiac leiomodin (C-LMOD), Leiomodin | LMOD2 | 7 | 547 | Nucleation | Bind along F-actin, next to the pointed end. Involved in sarcomere assembly and organization. | Neonatal dilated cardiomyopathy. | ABS1, ABS2 (LRR domain), ABS3 (WH2 domain) | https://www.protein |
| Leiomodin-3, Leiomodin, fetal form | LMOD3 | 3 | 560 | Nucleation | Bind along F-actin, next to the pointed end. Involved in sarcomere assembly and organization. | Nemaline myopathy | ABS1, ABS2 (LRR domain), ABS3 (WH2 domain) | https://www.protein |

| | | | | | | | | |
|---|---|---|---|---|---|---|---|---|
| Arp2/3 complex (Actin-related protein 2, Actin-related protein 3, Actin-related protein 3B, Actin-related protein 2/3 complex subunit 1A, Actin-related protein 2/3 complex subunit 1B, Actin-related protein 2/3 complex subunit 2, Actin-related protein 2/3 complex subunit 3, Actin-related protein 2/3 complex subunit 4, Actin-related protein 2/3 complex subunit 5) | ACTR2, ACTR3, ACTR3B, ARPC1A, ARPC1B, ARPC2, ARPC3, ARPC4, ARPC5 | 2<br>2<br>7<br>7<br>7<br>2<br>12<br>3<br>1 | 394<br>418<br>418<br>370<br>372<br>300<br>178<br>168<br>151 | Nucleation, branching, capping | Generates branched actin networks. Regulate actin-based cell motility. | Lung, breast, gliomas, gastric, pancreatic and colorectal cancers, psoriasis-like disease, psoriasis-like disease, intervertebral disc and cartilage degeneration, chronic obstructive pulmonary disease, Hirschsprung disease. | | https://www.proteinatlas.org/ENSG00000138071-ACTR2<br>https://www.proteinatlas.org/ENSG00000115091-ACTR3<br>https://www.proteinatlas.org/ENSG00000133627-ACTR3B<br>https://www.proteinatlas.org/ENSG00000241685-ARPC1A<br>https://www.proteinatlas.org/ENSG00000130429-ARPC1B<br>https://www.proteinatlas.org/ENSG00000163466-ARPC2<br>https://www.proteinatlas.org/ENSG00000111229-ARPC3<br>https://www.proteinatlas.org/ENSG00000241553-ARPC4<br>https://www.proteinatlas.org/ENSG00000162704-ARPC5 |
| Actin-related protein 3C, Actin-related protein 11 | ACTR3C, ARP11 | 7 | 210 | Capping | Decreases the formation of the Arp1 assemblies. Cap pointed end. | | | https://www.protein |
| Wiskott-Aldrich syndrome protein, WASp | WAS, IMD2 | X | 502 | Polymerization, cross-linking | Effector protein for Rho-type GTPases, interaction with the Arp2/3 complex. Promote actin polymerization in the nucleus. | Wiskott-Aldrich syndrome, Thrombocytopenia 1, Neutropenia, severe congenital, X-linked (XLN). | WH1, WH2 | https://www.protein |
| Wiskott-Aldrich syndrome protein family member 1 (WASP family protein member 1), Protein WAVE-1, Verprolin homology domain-containing protein 1 | WASF1, KIAA0269, SCAR1, WAVE1 | 6 | 559 | Monomer binding | The WAVE complex regulates actin filament reorganization via its interaction with the Arp2/3 complex. It restricts actin network extension at the leading edge. | Alzheimer's disease, acute myeloid leukemia, oral squamous cell carcinoma, breast, prostate, ovarian, melanoma cancer. | WH2 | https://www.protein |

| | | | | | | | | |
|---|---|---|---|---|---|---|---|---|
| WAS protein family member 2, Verprolin homology domain-containing protein 2 | WASF2, WAVE2 | 1 | 498 | Monomer binding | The WAVE complex regulates actin filament reorganization via its interaction with the Arp2/3 complex. It promotes actin network extension at the leading edge. | Lung, liver, pancreatic, prostate, colorectal and breast, cervical cancers, left-sided obstructive heart defects, osteosarcoma, melanoma, Parkinson's disease. | WH2 | https://www.protein |
| Neural Wiskott-Aldrich syndrome protein, N-WASP | WASL | 7 | 505 | Monomer binding, Depolymerization | Stimulating the actin-nucleating activity of the Arp2/3 complex; bind with CDC42, involved in the extension and maintenance of the formation filopodia. | Parkinson's disease, Wiskott-Aldrich syndrome, hepatocellular carcinoma, polycystic kidney disease, esophageal cancer, gastric cancer, proteinuria kidney disease, breast cancer. | WH1, WH2 | https://www.protein |
| WAS/WASL-interacting protein family member 1, Protein PRPL-2, Wiskott-Aldrich syndrome protein-interacting protein | WIPF1, WASPIP, WIP | 2 | 503 | Monomer binding | Activation of N-WASP; activator of Arp2/3. | Wiskott-Aldrich syndrome 2. | WH2 | https://www.protein |
| WAS/WASL-interacting protein family member 2, WASP-interacting protein-related protein, WIP- and CR16-homologous protein, WIP-related protein | WIPF2, WICH, WIRE | 17 | 440 | Stabilization, Bunding | Cooperate with WASP and N-WASP to induce mobilization and reorganization of the actin filament system. | Parkinson's disease, hepatocellular carcinoma, breast cancer. | WH2 | https://www.protein |
| WAS/WASL-interacting protein family member 3, Corticosteroids and regional expression protein 16 homolog | WIPF3, CR16 | 7 | 483 | Monomer binding | Cooperate with N-WASP to regulate actin polymerization. | Abdominal aortic aneurysm. | WH2 | https://www.protein |

| | | | | | | | | |
|---|---|---|---|---|---|---|---|---|
| Tropomodulin 1, Erythrocyte tropomodulin | TMOD1, D9S57E, ETMOD, TMOD | 9 | 359 | Capping | Capping the pointed end. Interaction with tropomyosin stabilizes thin filaments in cardiac myocytes. | Breast cancer, glaucomatous retina, neurological diseases, diabetic kidney disease, chronic obstructive pulmonary disease, lung adenocarcinoma, congenital myopathy, meningiomas, acute lymphoblastic leukemia, ulcerative colitis (UC), Duchenne muscular dystrophy (DMD) | ABS1, ABS2 | https://www.protein |
| Tropomodulin 2, Neuronal tropomodulin | TMOD2, NTMOD | 15 | 351 | Capping | Capping the pointed end. Regulate behavior, learning, memory, and synaptic plasticity. | Neurological diseases, Idiopathic pulmonary fibrosis (IPF), first-episode schizophrenia. | ABS1, ABS2 | https://www.protein |
| Tropomodulin 3, Ubiquitous tropomodulin | TMOD3, UTMOD | 15 | 352 | Capping | Capping the pointed end. Regulate cell morphology, motility, and division. | Liver cancer, neurological diseases. | ABS1, ABS2 | https://www.protein |
| Tropomodulin 4, Skeletal muscle tropomodulin | TMOD4, Sk-Tmod | 1 | 345 | Capping | Capping the pointed end. Skeletal muscle development. | Neurological diseases, myocardial infarction, early-onset dyslipidemia, primary auditory neurons degeneration (PAND) | ABS1, ABS2 | https://www.protein |
| Adseverin, Scinderin | SCIN, KIAA1905 | 7 | 715 | Severing, Capping, Nucleation | F-actin depolymerization. Regulate cell proliferation. Gelsolin superfamily protein. | periodontal disease, bladder cancer, hepatocellular carcinoma, gastric cancer, Colorectal Cancer, breast cancer, acute myeloid leukemia, prostate cancer, melanoma, multiple sclerosis, head and neck cancer. | Gelsolin-like domain | https://www.protein |

Vedula et al 2023 selected list of actin binding proteins (adapted from Gao and Nakamua 2022)

| | | | | | | | | |
|---|---|---|---|---|---|---|---|---|
| Advillin, p92 | AVIL | 12 | 819 | Bunding | Regulate morphogenesis of neuronal cells. Gelsolin superfamily protein | Steroid-resistant nephrotic syndrome, glioblastoma. | Headpiece domain | https://www.protein |
| Supervillin, Archvillin, p205/p250 | SVIL | 10 | 2214 | Bunding | Regulate actin dynamics. Gelsolin superfamily protein. | Liver cancer, myopathy, prostate cancer, Lymphangiol eiomyomatosi s. | Gelsolin-like domain | https://www.protein |
| Twinfilin-1, protein A6, protein tyrosine kinase 9 | TWF1, PTK9 | 12 | 350 | Monomer binding, Severing | Inhibit actin polymerization. Regulate motility. | Ocular coloboma, pancreatic cancer, breast cancer, lung adenocarcino ma, hepatocellular carcinoma, coronary artery disease. | ADF-H | https://www.protein |
| Twinfilin-2, A6-related protein, Twinfilin-1-like protein | TWF2, PTK9L, MSTP011 | 3 | 349 | Monomer binding | Inhibit actin polymerization by sequestering actin monomers and capping filament barbed ends. | Pancreatic cancer. | ADF-H | https://www.protein |
| Drebrin-like protein, Cervical SH3P7, Cervical mucin-associated protein, Drebrin-F, HPK1-interacting protein of 55 kDa | DBNL, CMAP, SH3P7 | 7 | 430 | Depolymerization | Reorganize the actin cytoskeleton. | | ADF-H | https://www.protein |
| Adenylyl cyclase-associated protein 1 | CAP1, CAP | 1 | 475 | Monomer binding | Rapid actin filament depolymerization. | Head and neck squamous cell carcinomas. Chronic kidney disease. | C-CAP | https://www.protein |
| Adenylyl cyclase-associated protein 2 | CAP2 | 6 | 477 | Monomer binding | Blocking G-actin, rapid actin filament depolymerization. | Gliomas. | C-CAP | https://www.protein |
| Erythroid spectrin Spectrin (Spectrin alpha chain, erythrocytic 1, Erythroid alpha-spectrin; Spectrin beta chain, erythrocytic; Beta-I spectrin) | SPTA1, SPTA; SPTB1 | 1<br>14 | 2419<br>2137 | Bundling, Anchoring, Nucleation | Maintains cell membrane integrity and its mechanical properties. involve in cell adhesion and spreading, form lamellipodia, and also participate in morphogenetic processes. | Duchenne Muscular Dystrophy, Hereditary Spherocytosis, hereditary elliptocytosis, hereditary pyropoikilocytosis, | CH1, CH2 | https://www.proteinatlas.org/ENSG00000163554-SPTA1<br>https://www.proteinatlas.org/ENSG00000070182-SPTB |

| | | | | | | | | |
|---|---|---|---|---|---|---|---|---|
| Non-erythrocytic spectrin, fodrin (Spectrin alpha chain, non-erythrocytic 1, Alpha-II spectrin, Fodrin alpha chain, Spectrin, non-erythroid alpha subunit; Spectrin beta chain, non-erythrocytic 1; Beta-II spectrin; Fodrin beta chain; Spectrin, non-erythroid beta chain 1) | SPTAN1, NEAS, SPTA2; SPTBN1, SPTB2 | 9<br>2 | 2472<br>2364 | Bundling, Anchoring, Nucleation | Maintenance of structural integrity in mammalian cells, which is necessary for proper cell function. | Moyamoya disease (MMD), Colorectal, gastric, lung, breast, prostate, ovatian cancers, cutaneous, soft tissue tumors, non-Hodgkin lymphoma, acute lymphocytic leukemia. | CH1, CH2 | https://www.protein |
| Ezrin, Cytovillin, Villin-2, p81 | EZR, VIL2 | 6 | 586 | Anchoring | Crosslinker of actin with membrane; stimulate the actin polymerization. | Breast, lung and prostate cancers, oral squamous cell carcinomas (OSCCS). | C-terminal ABD | https://www.protein |
| Radixin | RDX | 11 | 583 | Anchoring, Capping | Capping the barbed end; inhibit the actin polymerization. | Prostate cancer. | C-terminal ABD | https://www.protein |
| Moesin, Membrane-organizing extension spike protein | MSN | X | 577 | Anchoring | Crosslinker of actin with membrane. | Oral squamous cell carcinomas (OSCCS). | C-terminal ABD | https://www.protein |
| Merlin, Moesin-ezrin-radixin-like protein, Neurofibromin-2, Schwannomerlin, Schwannomin | NF2, SCH | 22 | 595 | Anchoring, Stabilization | Tumor suppressor. | Neurofibromatosis type 2 (NF2), multiple nervous system tumors, vestibular and spinal schwannomas, meningiomas and ependymomas, mesotheliomas, breast, prostate, colorectal, hepatic, clear cell renal cell carcinoma, and melanomas. | N-terminal ABD (178-367) | https://www.protein |

| | | | | | | | | |
|---|---|---|---|---|---|---|---|---|
| Coronin-1A, Coronin-like protein A, Tryptophan aspartate-containing coat protein | CORO1A, CORO1 | 16 | 461 | Stabilization, Bundling | Disassemble actin filament branches. Regulate phagocytosis, locomotion, and cytokinesis. | Immunodeficiency 8 (IMD8), severe combined immunodeficiency (SCID), systemic lupus erythematosus (SLE), multiple sclerosis (MS), neurocognitive and behavioral abnormal, pathogenic infection. | N- and C-terminal | https://www.protein |
| Coronin-1B, Coronin-2 | CORO1B | 11 | 489 | Stabilization, Bundling | Controls actin networks at classical lamellipodia. | | N-terminal (Arg30 is crucial) | https://www.protein |
| Coronin 1C, Coronin-3, hCRNN4 | CORO1C, HCRNN4 | 12 | 474 | Stabilization, Bundling | Regulate cell migration and metastasis. | Gastric, colorectal, breast cancers, lung squamous cell carcinoma, renal cell cancer, Diffuse Gliomas, Hepatocellular Carcinoma (HCC). | C-terminal | https://www.protein |
| Coronin-2A, IR10, WD repeat-containing protein 2, Coronin-4 | CORO2A, IR10, WDR2 | 9 | 525 | Stabilization, Bundling | Regulate focal adhesion turnover. Bind to the promoter of NCoR target gene. | Inflammatory. | | https://www.protein |
| Coronin-2B, Coronin-like protein C (Clipin-C), Protein FC96, Coronin-5 | CORO2B | 15 | 480 | Stabilization, Bundling | Limiting the speed of actin polymerization. | Diabetic nephropathy (DN). | N-terminal | https://www.protein |
| Coronin-6, Coronin-like protein E (Clipin-E) | CORO6 | 17 | 472 | Anchoring | Anchor acetylcholine receptors to actin. | Congenital myasthenic syndrome (CMS). | N-terminal | https://www.protein |
| Coronin-7 (Crn7), 70 kDa WD repeat tumor rejection antigen homolog | CORO7 | 16 | 925 | Stabilization | Facilitate vesicular trafficking. Regulate Golgi structure. | Obesity. | | https://www.protein |

| | | | | | | | | |
|---|---|---|---|---|---|---|---|---|
| Src substrate cortactin, Amplaxin, Oncogene EMS1 | CTTN, EMS1 | 11 | 550 | Stabilization | Stabilize new filament branch points. directly activate Arp2/3, form microspikes. | Head and neck squamous cell carcinoma (HNSCC), oral squamous cell carcinoma, lung squamous cell carcinoma, gliosarcoma, breast cancer, colorectal cancer and melanoma, leukemia, cardiovascular diseases (CVDs), cerebral cavernous malformations (CCM), inflammatory Bowel diseases, acute lung injury, Alzheimer’s disease (AD), hypogonadatropic hypogonadism (HH), myositis. | 6.5 tandem repeats | https://www.protein |
| Hematopoietic lineage cell-specific protein, Hematopoietic cell-specific LYN substrate 1, LckBP1, p75 | HCLS1, HS1 | 3 | 486 | Stabilization | Stabilizing newly formed branched actin networks. | Chronic lymphocytic leukemia (CLL), Systemic lupus erythematosus (SLE), severe congenital neutropenia (SCN), Alzheimer’s disease (AD). | Repetitive tandem repeats and the coiled-coil (CC) domain | https://www.protein |
| F-actin-capping protein subunit alpha-1, CapZ alpha-1 | CAPZA1 | 1 | 286 | Capping | Blocking the exchange of subunits at barbed end. | Gastric cancer, asthma, chronic obstructive pulmonary disease (COPD), pancreatic cancer | | https://www.protein |
| F-actin-capping protein subunit alpha-2, CapZ alpha-2 | CAPZA2 | 7 | 286 | Capping | Blocking the exchange of subunits. | Non-syndromic neurodevelopmental disorder in children. | | https://www.protein |
| F-actin-capping protein subunit alpha-3, CapZ alpha-3 | CAPZA3, CAPPA3, GSG3 | 12 | 299 | Capping | Controlling actin polymerization during spermiogenesis. | Male infertility | | https://www.protein |

Vedula et al 2023 selected list of actin binding proteins (adapted from Gao and Nakamua 2022)

| | | | | | | | | |
|---|---|---|---|---|---|---|---|---|
| F-actin-capping protein subunit beta, CapZ beta | CAPZB | 1 | 277 | Capping | Blocks actin polymerizationand depolymerization at the fast growing (barbed) filament ends. | Alzheimer's disease; ovarian cancer; sporadic amyotrophic lateral sclerosis and frontotemporal lobar degeneration comorbidity (ALS/FTD). | | https://www.protein |
| Destrin, Actin-depolymerizing factor (ADF) | DSTN, ACTDP, DSN | 20 | 165 | Severing | Actin depolymerizing. Sever actin filaments. | Lung adenocarcinoma, Alzheimer's disease and ischemic kidney disease. | ADF-H | https://www.protein |
| Gelsolin, AGEL, Actin-depolymerizing factor | GSN | 9 | 782 | Capping, Severing, Nucleation | Prevents further monomer binding and eventually depolymerizes actin filament; promote the polymerization of monomers into filaments (nucleation) as well as sever filaments. | Cancer, amyloidosis, rheumatoid arthritis, AD, autoimmune diseases, chronic kidney disease, diabetes type 2. | G1, G2 and G4 | https://www.protein |
| Villin-1 | VIL1, VIL | 2 | 827 | Capping, Severing, Nucleation, Bundling | Actin nucleation, actin filament bundling, actin filament capping and severing. Gelsolin superfamily protein. | Biliary atresia. | Core fragment in N terminal; headpiece domain in C terminal | https://www.protein |
| Alpha-actinin-1, Alpha-actinin cytoskeletal isoform, F-actin cross-linking protein, Non-muscle alpha-actinin-1 | ACTN1 | 14 | 892 | Bundling | Anchor actin to a variety of intracellular structures. | Bleeding disorder, platelet-type 15 (BDPLT15), congenital macrothrombocytopenia. | CH1, CH2 | https://www.protein |

| | | | | | | | | |
|---|---|---|---|---|---|---|---|---|
| Alpha-actinin-2, Alpha-actinin skeletal muscle isoform 2 | ACTN2 | 1 | 894 | Bundling | Anchor actin to a variety of intracellular structures. | Cardiomyopathy, familial hypertrophic 23, with or without left ventricular non-compaction (CMH23); Cardiomyopathy, dilated 1AA, with or without left ventricular non-compaction (CMD1AA); Myopathy, congenital, with structured cores and Z-line abnormalities (MYOCOZ); Myopathy, distal, 6, adult onset, autosomal dominant (MPD6). | CH1, CH2 | https://www.protein |
| Alpha-actinin-3, Alpha-actinin skeletal muscle isoform 3 | ACTN3 | 11 | 901 | Bundling | Skeletal muscle growth. | Sarcopenia, bone loss. | CH1, CH2 | https://www.protein |
| Alpha-actinin-4, Non- muscle alpha-actinin 4 | ACTN4 | 19 | 911 | Bundling | Regulate cell motility and invasion. | Focal segmental glomerulosclerosis 1 (FSGS1). | CH1, CH2 | https://www.protein |
| Cofilin-1 | CFL1, CFL | 11 | 166 | Severing | pH-sensitive F-actin depolymerizing activity. | Alzheimer disease; Parkinson's disease; ischemic kidney disease; colorectal cancer; Dent's disease; urothelial cancer; autoinflammatory disease and macrothrombocytopenia; neurodegenerative disease. | ADF-H | https://www.protein |
| Cofilin-2 | CFL2, NEM7 | 14 | 166 | Severing | Controls reversibly actin polymerization and depolymerization in a Ph-sensitive manner. Its F-actin depolymerization activity is regulated by association with CSPR3. | Human muscle disorder; Duchenne muscular dystrophy; nemaline myopathy 7. | ADF-H | https://www.protein |

Vedula et al 2023 selected list of actin binding proteins (adapted from Gao and Nakamua 2022)

| | | | | | | | | |
|---|---|---|---|---|---|---|---|---|
| Dematin, Dematin actin- binding protein, Erythrocyte membrane Protein band 4.9 | DMTN, DMT, EPB49 | 8 | 405 | Bundling, Stabilization, Anchoring | Induces F-actin bundles formation and stabilization; attaches the spectrin-actin network to the erythrocytic plasma membrane. | Autosomal dominant Marie Unna hereditary hypotrichosis disease; prostate cancer. | N-terminal core domain and the C-terminal headpiece domain | https://www.protein |
| Filamin-A, Actin-binding protein 280, Endothelial actin-binding protein, Filamin-1, Non- muscle filamin | FLNA, FLN, FLN1 | X | 2647 | Cross-linking, Scaffolding | Scaffolding | Periventricular nodular heterotopia 1 (PVNH1), Otopalatodigital syndrome 2 (OPD2), Frontometaphyseal dysplasia 1 (FMD1), Frontometaphyseal dysplasia 1 (FMD1), Intestinal pseudoobstruction, neuronal, chronic idiopathic, X-linked (IPOX), FG syndrome 2 (FGS2), Terminal osseous dysplasia (TOD), Cardiac valvular dysplasia, X-linked (CVD1), Cardiac valvular dysplasia, X-linked (CVD1) | N-terminal ABD (CH1, CH2), R10 | https://www.protein |
| Filamin-B, ABP-278, ABP-280 homolog, Actin-binding-like protein, Beta-filamin, Filamin homolog 1, Thyroid autoantigen, Truncated actin-binding protein | FLNB, FH1, FLN1L, TABP, TAP | 3 | 2602 | Cross-linking, Scaffolding | Scaffolding | Spondylocarpotarsal synostosis (SCT), Larsen syndrome (LS), atelosteogenesis (AO), boomerang dysplasia (BD), and isolated congenital talipes equinovarus. | N-terminal ABD (CH1, CH2) | https://www.protein |
| Filamin-C, ABP-280-like protein, Actin-binding-like protein, Filamin-2, Gamma-filamin | FLNC, ABPL, FLN2 | 7 | 2725 | Cross-linking, Scaffolding | Cross-links F-actin in the Z-disc. | Myopathies, Distal and Myofibrillar Skeletal Myopathy, Cardiomyopathy, Limb-girdle muscular dystrophy | N-terminal ABD (CH1, CH2) | https://www.protein |

| | | | | | | | | |
|---|---|---|---|---|---|---|---|---|
| Macrophage-capping protein, Actin regulatory protein CAP-G | CAPG, AFCP, MCP | 2 | 348 | Capping | Block the barbed ends of actin filament. | Breast cancer, bladder cancer, Atherosclerosis, Malignant mesothelioma, ovarian carcinoma, clear cell renal cell carcinoma (ccRCC), lung adenocarcinoma, oral squamous-cell carcinoma, carotid atherosclerosis, prostate cancer, Rheumatoidarthritis, osteoarthritis. | Gelsolin-like 1, Gelsolin-like 2, Gelsolin-like 3 | https://www.protein |
| Calponin-1, Basic calponin, Calponin H1, smooth muscle | CNN1 | 19 | 297 | Cross-linking, polymerization | Inhibit the actomyosin Mg-ATPase activity. Specific to smooth muscle cells. | Hepatocellular carcinoma, renal angiomyolipoma, papillary carcinomas, basal cell-like breast carcinoma, metastatic basal cell carcinomas, and prostate cancer, fibrosarcoma, leiomyosarcoma, synovial sarcoma and osteosarcoma. | CH, two actin-binding sites in the middle region. | https://www.protein |
| Calponin-2, Calponin H2, smooth muscle, Neutral calponin | CNN2 | 19 | 309 | Cross-linking, polymerization | Regulate multiple actin cytoskeleton-based functions. | Prostate and breast cancer. | CH, two actin-binding sites in the middle region. | https://www.protein |
| Calponin-3, Calponin, acidic isoform | CNN3 | 1 | 329 | Cross-linking, polymerization | Participates in actin cytoskeleton-based activities in embryonic development and myogenesis. | Seizure | CH, two actin-binding sites in the middle region. | https://www.protein |
| Transgelin, 22 kDa actin-binding protein, Protein WS3-10, Smooth muscle protein 22-alpha | TAGLN, SM22, WS3-10 | 11 | 201 | Bundling | Gelation and stabilization. Organization of the actin cytoskeleton, cell migration and response to stress. | Breast, prostate and colon cancers. | Actin-binding motif (ABM) | https://www.protein |

| | | | | | | | | |
|---|---|---|---|---|---|---|---|---|
| Transgelin-2, Epididymis tissue protein Li 7e, SM22-alpha homolog | TAGLN2, KIAA0120 | 1 | 199 | Polymerization, Bundling | Blocks Arp2/3-nucleated actin branching. | Gliomas, infertility, lupus erythematosus, bladder, colorectal, hepatocellular and lung cancers, maxillary sinus squamous cell carcinoma, uterine cervical squamous cell carcinoma, Asthma. | Actin-binding motif (ABM) | https://www.protein |
| Transgelin 3, Neuronal protein 22 (NP22) | TAGLN3, NP25 | 3 | 199 | Bundling | Regulate actin stabilization and actomyosin contractility. | | Actin-binding motif (ABM) | https://www.protein |
| Nesprin-1, Enaptin, KASH domain-containing protein 1 (KASH1, Myne-1), Synaptic nuclear envelope protein 1, Synaptic nuclear envelope protein 1 (Syne-1) | SYNE1, C6orf98, KIAA0796, KIAA1262, KIAA1756, MYNE1 | 6 | 8797 | Anchoring | Anchor the F-actin cytoskeleton to the nuclear envelope. | Autosomal recessive cerebellar ataxia (ARCA), schizophrenia, Disrupted-In-Schizophrenia 1 (DISC1), autism spectrum disorder (ASD), Emery Dreifuss muscular dystrophy (EDMD), Cerebellar ataxia accompanying motor neuron disease (juvenile-onset amyotrophic lateral sclerosis mimic), cardiac disease, arthrogryposis multiplex congenita (AMC), breast, lung, ovarian, pancreatic, head and neck and colorectal cancers. | ABD (1-289) | https://www.protein |

Vedula et al 2023 selected list of actin binding proteins (adapted from Gao and Nakamua 2022)

| | | | | | | | | |
|---|---|---|---|---|---|---|---|---|
| Nesprin-2, KASH domain-containing protein 2 (KASH2), Nucleus and actin connecting element protein (NUANCE, Synaptic nuclear envelope protein 2 (Syne-2) | SYNE2, KIAA1011, NUA | 14 | 6885 | Anchoring | Anchor the F-actin cytoskeleton to the nuclear envelope. | Emery Dreifuss muscular dystrophy (EDMD), breast, lung, ovarian, pancreatic, head and neck and colorectal cancers. | ABD (1-286) | https://www.protein |
| Brain-specific angiogenesis inhibitor 1-associated protein 2-like protein 1, Insulin receptor tyrosine kinase substrate | BAIAP2L1, IRTKS | 7 | 511 | Bundling | Induce actin microspikes. | Ovarian cancer, lung adenocarcinoma, bladder cancer, clear cell renal cell carcinoma, pancreatic cancer, lung cancer, prostate cancer, spitzoid melanoma, gastric cancer, rheumatoid arthritis, hepatocellular carcinoma. | IRSp53/MIM homology domain (IMD, 1-249 | https://www.protein |

Vedula et al 2023 selected list of actin binding proteins (adapted from Gao and Nakamua 2022)

| | | | | | | | | | |
|---|---|---|---|---|---|---|---|---|---|
| Fascin, 55 kDa actin-bundling protein, Singed-like protein, p55 | FSCN1, FAN1, HSN, SNL | 7 | | 493 | Bundling | Organizes F-actin into parallel bundles. Promote metastasis. | Colorectal cancer, ovarian cancer, renal cell carcinoma, prostate cancer, adrenocortical carcinoma, thyroid carcinoma, squamous cell carcinoma, cervical carcinoma, pituitary adenoma, breast cancer, non-small cell lung cancer, esophageal cancer, hepatocellular carcinoma, gastric cancer, bladder cancer, pancreatic cancer, nasopharyngeal carcinoma, adrenocortical cancer, metastatic melanoma, chondrosarcoma. | Actin-binding sites (ABS) 1-3. | https://www.protein |
| Fascin-2, Retinal fascin | FSCN2, | 17 | | 492 | Bundling | Play a pivotal role in photoreceptor cell-specific events, such as disk morphogenesis. | Retinopathy, Hearing Loss, autosomal dominant retinitis pigmentosa. | ABS | https://www.protein |
| Fascin-3, Testis fascin | FSCN3 | 7 | | 498 | Bundling | Testis-specific. | Nonobstructive azoospermia. | | https://www.protein |
| LIM domain and actin-binding protein 1, Epithelial protein lost in neoplasm | LIMA1, EPLIN, SREBP3 | 12 | | 759 | Bundling, Stabilization | Formation of stress fibers. Inhibit membrane ruffling. | Oral cancer, breast cancer, prostate cancer, squamous cell carcinoma of head and neck (SCCHN), lung cancer, oesophageal cancer, ovarian cancer, colorectal cancer (CRC), gastric cancer. | Actin binding region | https://www.protein |

Vedula et al 2023 selected list of actin binding proteins (adapted from Gao and Nakamua 2022)

| | | | | | | | | |
|---|---|---|---|---|---|---|---|---|
| Protein flightless-1 homolog | FLII, FLIL | 17 | 1269 | Monomer binding, Capping, Severing | Regulate actin dynamics. Gelsolin superfamily protein. | OVA-Induced Atopic Dermatitis Skin-Like Disease, epidermolysis bullosa acquisita (EBA), prostate cancer, epithelial ovarian cancer, breast cancer, endometrial cancer, alcoholic liver disease, Ulcerative colitis, psoriasiform dermatitis, cutaneous leishmaniasis, Systemic lupus erythematosus (SLE), melanoma. | gelsolin-like domain | https://www.protein |
| Four and a half LIM domains protein 3 (FHL-3), Skeletal muscle LIM-protein 2 (SLIM-2) | FHL3, SLIM2 | 1 | 280 | ? | Inhibit alpha-actinin-mediated actin bundling. Enhance cell spreading and stress fiber disassembly. | Pancreatic, gastric breast cancers, glioma. | LIM domains | https://www.protein |
| WD repeat-containing protein 1 , Actin-interacting protein 1 (AIP1, NORI-1) | WDR1 | 4 | 606 | Depolymerization | Enhances the filament disassembly activity of cofilin and restricts cofilin localization to cortical actin patches. Regulate cytokinesis and migration. | Gout, pancreatitis, and primary glioblastoma. | | https://www.protein |
| Plastin-1, Intestine-specific plastin (I-plastin), fimbrin | PLS1 | 3 | 629 | Bundling | Required for stereocilia formation. | Deafness, aut | CH (108-380, 381-625) | https://www.protein |
| Plastin-2, Lymphocyte cytosolic protein 1 (LCP-1), L-plastin, LC64P | LCP1, PLS2 | 13 | 627 | Bundling | Regulate T-cell activation. | B-cell non-Hodgkin lymphomas, coloboma, melanoma, breast, prostate, colorectal cancer | CH (106-379, 380-624) | https://www.protein |

| | | | | | | | | |
|---|---|---|---|---|---|---|---|---|
| Plastin-3, T-plastin | PLS3 | X | 630 | Bundling | Regulate actin-based cellular processes. Regulate bone development. | Acute myeloid leukemia (AML), Sézary syndrome (SS), Osteoporosis, osteoarthritis, colorectal, prostate, breast, gastric, and lung cancer, thoracic aortic dissection, Spinal muscular atrophy (SMA), ataxia, amyotrophic lateral sclerosis (ALS), and Charcot–Marie–Tooth (CMT), Infection and pathogen entry. | CH (1069-382, 383-627) | https://www.protein |
| KICSTOR complex protein kaptin, Actin-associated protein 2E4 | KPTN, 2E4 | 19 | 436 | ? | Localized in lamellipodia. | Deafness, KPTN-related syndrome. | Bind to F-actin column | https://www.protein |
| TRIO and F-actin-binding protein, Protein Tara, Trio-associated repeat on actin | TRIOBP, KIAA1662, TARA | 22 | 2365 | Stabilization | Regulate cell spreading and contraction. | Deafness, autosomal recessive 28 (DFNB28), Schizophrenia, gastric, rectal, pancreatic and brain cancer | R1 motif | https://www.protein |
| Tyrosine-protein kinase ABL1, Abelson murine leukemia viral oncogene homolog 1, Abelson tyrosine-protein kinase 1, Proto-oncogene c-Abl, p150 | ABL1, ABL, JTK7 | 9 | 1130 | Bundling | Membrane ruffling, cell spreading, cell migration, and neurite extension in response to growth factor and extracellular matrix signals. | Alzheimer's Disease, Parkinson's disease (PD), Lewy body dementia, neuro-inflammation, breast, prostate and renal cancer, leukemia. | C-terminal | https://www.protein |

| | | | | | | | | |
|---|---|---|---|---|---|---|---|---|
| Neurabin-1, Protein phosphatase 1 regulatory subunit 9A, Neural tissue-specific F-actin-binding protein I | PPP1R9A, KIAA1222 | 7 | 1098 | Cross-linking | Regulate cell morphology. Inhibit protein phosphatase 1-alpha activity. | Prostate cancer, schizophrenia and bipolar disorder, Hepatosplenic T-cell lymphoma (HSTL), Huntington disease, squamous cell carcinoma of head and neck (SCCHN), papillary thyroid cancer, breast cancer, restless legs syndrome (RLS). | ABD (1-144) | https://www.protein |
| Neurabin-2, Neurabin-II, Protein phosphatase 1 regulatory subunit 9B, Spinophilin | PPP1R9B, PPP1R6 | 17 | 817 | Cross-linking | Scaffolding | Alzheimer's disease, hypertension, hepatocellular carcinoma, Parkinson disease, breast, colorectal, lung, head and neck, colon, prostate cancers. | ABD (1-154, 164-283) | https://www.protein |
| Protein spire homolog 1 (Spir-1) | SPIRE1, KIAA1135, SPIR1 | 18 | 756 | Nucleation | Mediate asymmetric spindle positioning by assembling an actin network. Caps the pointed end. | | WH2 | https://www.protein |
| Protein spire homolog 2 (Spir-2) | SPIRE2, KIAA1832, SPIR2 | 16 | 714 | Nucleation | Mediate asymmetric spindle positioning by assembling an actin network. | | WH2 | https://www.protein |
| Drebrin, Developmentally-regulated brain protein | DBN1, D0S117E | 5 | 649 | Stabilization | The stability, dynamics, and organizations of actin structures in neuronal cells. | Alzheimer's disease, hereditary and acquired glomerulopathies, Down Syndrome, Seizures and Suspected Encephalitis, prostate cancer, lung adenocarcinoma,.glaucoma. | ADF-H | https://www.protein |

Vedula et al 2023 selected list of actin binding proteins (adapted from Gao and Nakamua 2022)

| | | | | | | | | |
|---|---|---|---|---|---|---|---|---|
| Tropomyosin alpha-1 chain, Alpha-tropomyosin, Tropomyosin-1 | TPM1, C15orf13, TMSA | 15 | 284 | Stabilization | Regulate contractile systems and cytoskeleton, modulates actin-myosin interaction. | Hypertrophic cardiomyopathy (HCM), Dilated cardiomyopathy (DCM), left ventricular noncompaction (LVNC), arrthythmogenic right ventricular cardiomyopathy. | ABS | https://www.protein |
| Tropomyosin beta chain, Beta-tropomyosin, Tropomyosin-2 | TPM2, TMSB | 9 | 284 | Stabilization | Regulates contractile systems and cytoskeleton, modulates actin-myosin interaction. | Myopathies and distal arthrogryposis. | ABS | https://www.protein |
| Tropomyosin alpha-3 chain, Gamma-tropomyosin, Tropomyosin-3, Tropomyosin-5 (Htm5) | TPM3 | 1 | 285 | Stabilization | Regulates contractile systems and cytoskeleton, modulates actin-myosin interaction. | Myopathies and distal arthrogryposis. | ABS | https://www.protein |
| Tropomyosin alpha-4 chain, TM30p1, Tropomyosin-4 | TPM4 | 19 | 248 | Stabilization | Regulates contractile systems and cytoskeleton, modulates actin-myosin interaction. | Macrothrombocytopenia, Muscle diseases. | ABS | https://www.protein |
| Xin actin-binding repeat-containing protein 1, Cardiomyopathy-associated protein 1 | XIRP1, CMYA1, XIN | 3 | 1843 | Stablization | Required for cardiac development and cardiac function. Protects actin filaments from depolymerization. | Cardiomyopathy, arrhythmias, heart disease, intercalated disc (ICD). | Xin-repeats | https://www.protein |
| Xin actin-binding repeat-containing 2, Beta-xin, Cardiomyopathy-associated protein 3, Xeplin | XIRP2, CMYA3 | 2 | 3374 | Stablization | Required for long-term maintenance of hair cell stereocilia. | Hearing loss | Xin-repeats | https://www.protein |

| | | | | | | | | |
|---|---|---|---|---|---|---|---|---|
| Microtubule-actin cross-linking factor 1, isoforms 1/2/3/5, 620 kDa actin-binding protein (ABP620), Actin cross-linking family protein 7, Macrophin-1, Trabeculin-alpha | MACF1, ABP620, ACF7, KIAA0465, KIAA1251 | 1 | 7388 | Cross-linking | Modulate actin and microtubule cytoskeletal networks. | Spectraplakinopathy type I, schizophrenia, Parkinson's disease (PD), breast, lung, colorectal, liver cancer, gliomas and glioblastoma, renal cell carcinoma, steoporosis. Lissencephaly 9 with complex brainstem malformation (LIS9) | CH1, CH2 | https://www.protein |
| Allograft inflammatory factor 1 (AIF-1), Ionized calcium-binding adapter molecule 1, Protein G1 | AIF1, G1, IBA1 | 6 | 147 | Polymerization, Bundling | Enhance the actin-bundling activity of LCP1. Promote membrane ruffling and the phagocytosis of macrophages. Involved in Rac and calcium signaling pathways. | Experimental autoimmune neuritis. | | https://www.protein |
| Allograft inflammatory factor 1-like, Ionized calcium-binding adapter molecule 2 | AIF1L, C9orf58, IBA2 | 9 | 150 | Bundling | Regulates actomyosin contractility and filopodial extensions. | Breast cancer. | | https://www.protein |
| EF-hand domain-containing protein D1, EF-hand domain-containing protein 1, Swiprosin-2 | EFHD1, SWS2 | 2 | 239 | Bundling | Bind to β-actin in the mitochondrial matrix. | Colorectal cancer | | https://www.protein |
| EF-hand domain-containing protein D2 (EFhd2), Swiprosin-1 | EFHD2, SWS1 | 1 | 240 | Bundling | Regulate cell spreading and migration. | Lung adenocarcinoma, acute myeloid leukemia (AML), idiopathic cardiomyopathy, Schizophrenia, Parkinson’s disease (PD), Alzheimer’s disease (AD), Huntington’s disease (HD), amyotrophic lateral sclerosis (ALS). | | https://www.protein |

| | | | | | | | | |
|---|---|---|---|---|---|---|---|---|
| CD2-associated protein, Adapter protein CMS, Cas ligand with multiple SH3 domains | CD2AP | 6 | 639 | Capping, Anchoring | Actin barbed-end capping protein. May anchor the podocyte slit diaphragm to the actin cytoskeleton in renal glomerolus. | Focal segmental glomerulosclerosis 3 (FSGS3), Congenital nephrotic syndromes, Alzheimer's Disease | C-terminal | https://www.protein |
| Coactosin-like protein | COTL1, CLP | 16 | 142 | Polymerization | Regulate lamellipodia dynamics in part by protecting F-actin from cofilin-mediated disassembly. | Rheumatoid arthritis | ADF-H | https://www.protein |
| Brain-specific angiogenesis inhibitor 1-associated protein 2 (BAI-associated protein 2, BAI1-associated protein 2, Protein BAP2 Alternative), as ligand-associated factor 3 (FLAF3), Insulin receptor substrate p53/p58 (IRS-58, IRSp53/58), Insulin receptor substrate protein of 53 kDa (RSp53) | BAIAP2 | 17 | 552 | Bundling | Induce filopodia and the formation of tightly packed parallel F-actin bundles. | Schizophrenia, autism spectrum disorders (ASDs) and attention deficit/hyperactivity disorder (ADHD). | IRSp53/MIM homology domain (IMD) | https://www.protein |
| Actin filament-associated protein 1, 110 kDa actin filament-associated protein (AFAP-110) | AFAP1, AFAP | 4 | 730 | Cross-linking, Scaffolding | Act as an adaptor protein that links signaling molecules to actin filaments. Serve as a platform for the construction of larger signaling complexes. Serve as an activator of Src family kinases in response to cellular signals that alter its conformation. Effect actin organization, | Glaucoma | C-terminal ABD | https://www.protein |

| | | | | | | | | |
|---|---|---|---|---|---|---|---|---|
| Alpha-adducin, Erythrocyte adducin subunit alpha | ADD1, ADDA | 4 | 737 | Capping, Bundling | Recruite spectrin to actin filaments, bundling actin filaments and capping the barbed ends of actin filaments. Stabilize membrane cytoskeleton and cell-cell junctions. | Amyotrophic Lateral Sclerosis (ALS), ovarian cancer, non-cardia gastric cancer, colorectal cancer, lung cancer, small cell lung cancer, coronary heart disease, renal failure. | MARCKS-related domain | https://www.protein |
| Beta-adducin, Erythrocyte adducin subunit beta | ADD2, ADDB | 2 | 726 | Capping, Bundling | Recruite spectrin to actin filaments, bundling actin filaments and capping the barbed ends of actin filaments. Stabilize membrane cytoskeleton and cell-cell junctions. | Basal cell carcinoma and squamous cell carcinoma. | MARCKS-related domain | https://www.protein |
| Gamma-adducin, Adducin-like protein 70 | ADD3, ADDL | 10 | 706 | Capping, Bundling | Recruite spectrin to actin filaments, bundling actin filaments and capping the barbed ends of actin filaments. Stabilize membrane cytoskeleton and cell-cell junctions. | Small cell lung cancer, murine breast tumor, non-small cell lung cancer, colorectal cancer, glioblastoma, T-lymphoblastic leukemia. | MARCKS-related domain | https://www.protein |
| Anillin | ANLN | 7 | 1124 | Bundling | Organize the cortical actomyosin cytoskeleton in syncytial structures, involved in cell division. | Lung, Breast, pancreatic, colorectal, liver, bladder, urothelial, renal, nasopharyngeal, ovarian, hormone resistant prostate cancer. | ABD | https://www.protein |
| Espin, Autosomal recessive deafness type 36 protein, Ectoplasmic specialization protein | ESPN, DFNB36, LP2654 | 1 | 854 | Bundling | Connect to hair cell stereocilia and microvillar specializations of sensory cells in the inner ear. | Melanoma | Actin-bundling module (ABM) | https://www.protein |

| | | | | | | | | |
|---|---|---|---|---|---|---|---|---|
| Transcription activator BRG1, ATP-dependent helicase SMARCA4, BRG1-associated factor 190A (BAF190A), Mitotic growth and transcription activator, Protein BRG-1, Protein brahma homolog 1SWI/SNF related, matrix-associated actin-dependent regulator of chromatin subfamily A, member 4, SNF2-beta | SMARCA4, BAF190A, BRG1, SNF2B, SNF2L4 | 19 | 1647 | Capping | Induce the formation of thick actin filament bundles resembling stress-fibers. Change the morphology and alterations in actin cytoskeletal organization. | Non-small cell lung carcinomas, lung adenocarcinoma, lung large-cell carcinoma, lung squamous cell carcinoma, prostate, breast, pancreas, colon cancers. | C-terminal | https://www.protein |
| Palladin, SIH002, Sarcoma antigen NY-SAR-77 | PALLD, CGI-151, KIAA0992 | 4 | 1383 | Bundling, Scaffolding | Regulate cell morphology, motility, cell adhesion. | Myocardial infarction (MI), pancreatic ductal adenocarcinoma (PDA), colorectal cancer, pancreatic cancer, breast cancer. | Immunoglobulin-like domain (Ig-like domain 3) | https://www.protein |
| Myotilin, 57 kDa cytoskeletal protein, Myofibrillar titin-like Ig domains protein, Titin immunoglobulin domain protein | MYOT, TTID | 5 | 498 | Bundling | Regulate myofibril assembly and stability at the Z lines in muscle cells. | Late onset autosomal dominant distal limb girdle muscular dystrophy, spheroid body myopathy, myofibrillar myopathy. | C-terminal Ig-like domains | https://www.protein |
| Myopalladin, 145 kDa sarcomeric protein | MYPN, MYOP | 10 | 1320 | Bundling, Scaffolding | Component of Z-lines. | Dilated cardiomyopathy, Familial restrictive cardiomyopathy (FRCM). | Ig-like domain | https://www.protein |
| Alpha-parvin, Actopaxin, Calponin-like integrin-linked kinase-binding protein (CH-ILKBP), Matrix-remodeling-associated protein 2 | PARVA, MXRA2 | 11 | 372 | Scaffolding | Involved in integrin-mediated cell adhesion. | Lobular breast carcinoma, diabetic nephropathy. | CH1, CH2 | https://www.protein |

| | | | | | | | | |
|---|---|---|---|---|---|---|---|---|
| Caldesmon (CDM) | CALD1, CAD, CDM | 7 | 793 | Stabilization | Capable of stabilizing actin filaments against actin-severing proteins, inhibiting actomyosin ATPase activity, and inhibiting Arp2/3-mediated actin polymerization. Involved in smooth muscle contraction, cell motility and secretion. | Atherosclerosis, restenosis, glioma, Gastrointestinal stromal tumor, Ovarian adult granulosa cell tumor, Epithelioid pleural mesothelioma, Oral cavity squamous cell carcinoma, colorectal cancer, bladder cancer, Melanoma, Leiomyosarcoma, Fibroxanthoma. | 653-686, 768-793 | https://www.protein |
| Switch-associated protein 70 (SWAP-70) | SWAP70, KIAA0640 | 11 | 585 | Bundling | Alter the actin organization and lamellipodial morphology. | Prostate cancer | ABD | https://www.protein |
| Afadin, ALL1-fused gene from chromosome 6 protein (Protein AF-6), Afadin adherens junction formation factor | AFDN, AF6, MLLT4 | 6 | 1824 | Scaffolding | Enhance the formation of adherens and tight junctions. Effect on cell adhesion, polarization, migration, differentiation, and survival. | Breast cancer, Parkinson's Disease. | C-terminal (1631-1829) | https://www.protein |
| Dystrophin | DMD | X | 3685 | Stablization | Link the actin cytoskeleton to the dystroglycan complex in the plasma membrane. Bind to the intracellular actin network to link the cytoskeleton to dystrophin glycoprotein complex | Duchenne and Becker muscular dystrophies (DMD and BMD), X-linked dilated cardiomyopathy, | CH1 (15-119), CH2 (134-240) | https://www.protein |
| Dystroglycan 1, Dystrophin-associated glycoprotein 1, Dystroglycan | DAG1 | 3 | 895 | Bundling | Maintain sarcolemmal integrity. | Muscular dystrophy-dystroglycanopathy limb-girdle C9 (MDDGC9), Muscular dystrophy-dystroglycanopathy congenital with brain and eye anomalies A9 (MDDGA9) | The cytoplasmic tail | https://www.protein |
| Vasodilator-stimulated phosphoprotein | VASP | 19 | 380 | Polymerization, Bundling | Stimulate actin polymerization by promoting the transfer of profilin-bound actin monomers onto the barbed end of growing F-actin. | Listeria monocytogenes | Enabled/VASP homology (EVH2) contains a G-actin-binding site (GAB), an F-actin-binding site (FAB). | https://www.protein |

Vedula et al 2023 selected list of actin binding proteins (adapted from Gao and Nakamua 2022)

| | | | | | | | | |
|---|---|---|---|---|---|---|---|---|
| Protein enabled homolog | ENAH, MENA | 1 | 591 | Polymerization, Bundling | Regulate actin remodeling. Involved in carcinoma cell invasion and metastasis. | Carcinaoma. | EVH2 contains GAB and FAB. | https://www.protein |
| Ena/VASP-like protein, Ena/vasodilator-stimulated phosphoprotein-like protein | EVL, RNB6 | 14 | 416 | Polymerization, Bundling | Implicate in the regulation of axon guidance, platelet aggregation, cell motility, and cell adhesion. | Breast cancer | EVH2 contains GAB and FAB. | https://www.protein |
| Kelch-like protein 1 | KLHL1, KIAA1490 | 13 | 748 | Scaffolding | Bind to F-actin. Modulate neuronal structure and function. | Neurodegenerative diseases. | C-terminal Kelch β-propeller region | https://www.protein |
| Kelch-like protein 17, Actinfilin | KLHL17, AF | 1 | 642 | Scaffolding | Bind to F-actin. Form circular puncta in dendritic spines and are surrounded by or adjacent to F-actin. | Infantile spasms and autism. | C-terminal Kelch domain (289 – 641) | https://www.protein |
| Protein 4.1 (P4.1), Erythrocyte membrane protein band 4.1, 4.1R, Band 4.1, EPB4.1 | EPB41, E41P | 1 | 864 | Anchoring, Cross-linking | Involved in cytoskeletal rearrangements, intracellular transport and signal transduction. | Hereditary elliptocytosis (HE), bradycardia and/or Long QT syndrome, | Spectrin-actin-binding domain (SABD, 615-713) | https://www.protein |
| Band 4.1-like protein 2, Erythrocyte membrane protein band 4.1-like 2, Generally expressed protein 4.1 (4.1G) | EPB41L2 | 6 | 1005 | Anchoring, Cross-linking | Regulates cell adhesion, spreading, and migration. | | Spectrin-actin-binding domain (SABD, 611-676) | https://www.protein |
| Band 4.1-like protein 1, Erythrocyte membrane protein band 4.1-like 1, Neuronal protein 4.1, 4.1N. | EPB41L1 | 20 | 881 | Anchoring, Cross-linking | Regulate stability and plasticity of neuronal membrane. | Mental retardation, autosomal dominant 11 (MRD11). | Spectrin-actin-binding domain (SABD, 483-541) | https://www.protein |

| | | | | | | | | |
|---|---|---|---|---|---|---|---|---|
| Band 4.1-like protein 3 (4.1B), Differentially expressed in adenocarcinoma of the lung protein 1 (DAL-1), Erythrocyte membrane protein band 4.1-like 3 | EPB41L3, DAL1, KIAA0987 | 18 | 1087 | Anchoring, Cross-linking | Regulate cytoskeletal organization and a number of processes through multiple interactions. | Esophageal squamous cell carcinoma (ESCC), lung adenocarcinoma, meningiomas, breast cancer, ovarian cancer, prostate cancer, cervical cancer, gastric cancer, intestinal carcinoma, colorectal cancer, hepatocellular carcinoma and pancreatic carcinoma, esophageal carcinoma, renal clear cell carcinoma (RCCC). | Spectrin-actin-binding domain (SABD, 514-860) | https://www.protein |
| Talin-1 | TLN1, KIAA1027, TLN | 9 | 2541 | Cross-linking, Anchoring | Stabilize focal adhesion, involve in cellular mechanotransduction, provide a connection between the cytoskeleton and the ECM. | Myelodysplastic syndromes, prostate cancer, colon cancer, hepatocellular carcinoma, breast cancer, endometrioid carcinoma, glioblastoma, oral squamous cell carcinoma. | Actin-binding sites 1-3 | https://www.protein |
| Talin-2 | TLN2, KIAA0320 | 15 | 2542 | Cross-linking, Anchoring | Stabilize focal adhesion, involve in cellular mechanotransduction, provide a connection between the cytoskeleton and the ECM. | Breast cancer, hepatocellular carcinoma. | Actin-binding sites 1-3 | https://www.protein |
| Talin rod domain-containing protein 1, Mesoderm development candidate 1 | TLNRD1, MESDC1 | 15 | 362 | Bundling | Enhance filopodia formation and cell migration. | | Four-helix domain. | https://www.protein |

| | | | | | | | | |
|---|---|---|---|---|---|---|---|---|
| Nexilin F-actin binding protein, Nelin | NEXN | 1 | 675 | Polymerization, Cross-linking | Stimulate cell migration and adhesion. Maintenan Z line and sarcomere integrity. | Coronary artery disease, congenital heart disease, atrial septal defect, hypertrophic cardiomyopathy. | Central ABD | https://www.protein |
| Actin binding Rho-activating protein, Striated muscle activator of Rho-dependent signaling (STARS) | ABRA | 8 | 381 | Anchoring | Specifically expressed in cardiac and skeletal muscle cells. Bind to the I-band of the sarcomere. Act as a mechanosensor that translates skeletal muscle-specific stimuli into intracellular signals to promote serum-response factor -dependent gene transcription. | Cardiac hypertrophy and myopathy, ageing, type 2 diabetic muscle, skeletal muscle hypertrophy and atrophy. | 199-299, 300-381 | https://www.protein |
| Actin-binding LIM protein 1 (abLIM-1), Actin-binding LIM protein family member 1, Actin-binding double zinc finger protein, LIMAB1, Limatin | ABLIM1, ABLIM, KIAA0059, LIMAB1 | 10 | 778 | Scaffolding | Connect actin filaments and cytoplasmic targets. | Nasopharyngeal Carcinoma, Hepatocellular Carcinoma, Myotonic dystrophy type 1 (DM1). | dematin-like domain | https://www.protein |
| Actin-binding LIM protein 2 (abLIM-2), Actin-binding LIM protein family member 2 | ABLIM2, KIAA1808 | 4 | 611 | Scaffolding | Enhance STARS-dependent activation of serum-response factor. | Breast carcinogenesis, periodontitis, Alzheimer's Disease. | villin domain | https://www.protein |
| Actin-binding LIM protein 3 (abLIM-3), Actin-binding LIM protein family member 3 | ABLIM3, KIAA0843 | 5 | 683 | Scaffolding | Enhance STARS-dependent activation of serum-response factor.Involved in anchoring LIM domain-binding components of adherens junctions to circumferential actin bundles. | | villin domain | https://www.protein |

| | | | | | | | | |
|---|---|---|---|---|---|---|---|---|
| Tight junction protein ZO-1, Tight junction protein 1, Zona occludens protein 1, Zonula occludens protein 1 | TJP1, ZO1 | 15 | 1748 | Anchoring, Scaffolding | Link tight junction transmembrane proteins such as claudins and occludin to the actin cytoskeleton. | Inflammatory bowel disease, Kawasaki disease, Parkinson's disease, liver cancer, squamous cell carcinoma, Bowen's disease, celiac disease, gastrointestinal stromal tumor, endometrial carcinoma. | 1151-1371 | https://www.protein |
| Tight junction protein ZO-2, Zonula occludens protein 2 | TJP2, ZO2 | 9 | 1190 | Anchoring, Scaffolding | Link tight junction transmembrane proteins such as claudins and occludin to the actin cytoskeleton. | Familial hypercholanemia (FHCA), Cholestasis, progressive familial intrahepatic, 4 (PFIC4). | | https://www.protein |
| Tight junction protein ZO-3, onula occludens protein 3 | TJP3, ZO3 | 19 | 919 | Anchoring, Scaffolding | Link tight junction transmembrane proteins such as claudins and occludin to the actin cytoskeleton. | | | https://www.protein |
| Occludin | OCLN | 5 | 522 | Anchoring, Scaffolding | Transmembrane protein that forms and regulate tight junction. Maintain blood-brain barrier. | Pseudo-TORCH syndrome 1 (PTORCH1) | | https://www.protein |

Vedula et al 2023 selected list of actin binding proteins (adapted from Gao and Nakamua 2022)

| | | | | | | | | |
|---|---|---|---|---|---|---|---|---|
| LIM and SH3 domain protein 1 (LASP1), Metastatic lymph node gene 50 protein (MLN 50) | LASP1, MLN50 | 17 | 261 | Scaffolding | Enhance cancer cell migration and cell invasion. Involved in the differentiation and development of neurons. Involve in vesicular secretion. | Gastric carcinoma (GC), bladder cancer, esophageal squamous cell carcinoma, Breast carcinoma, Colorectal carcinoma, Ovarian carcinoma, Hepatocellular carcinoma, Renal cell carcinoma, Prostate carcinoma (PC), Medulloblastoma, Nasopharyngeal carcinoma (NC), Non-small lung cancer, Lung Adenocarcinoma, Choriocarcinoma, Gall bladder carcinoma, Thyroid carcinoma, Pancreatic carcinoma. | Nebulin-like repeat (NR1, NR2) | https://www.protein |
| Nebulette, Actin-binding Z-disk protein | NEBL, C10orf113, LNEBL | 10 | 1014 | Bundling | Enhances cancer cell migration but reduces cell invasion. Involve in cell spreading | Colorectal cancer, non-small cell lung cancer (NSCLC). | Nebulin-like repeat (NR1, NR2, NR3) | https://www.protein |
| Actin-histidine N-methyltransferase, Protein-L-histidine N-tele-methyltransferase, SET domain-containing protein 3 (hSETD3) | SETD3, C14orf154 | 14 | 594 | Polymerization | Mediate histidine methylation on β-actin, and accelerate the assembly of actin filaments. | Muscle loading and hypertrophy | SET domain (94 – 314) | https://www.protein |
| Synaptopodin | SYNPO, KIAA1029 | 5 | 929 | Scaffolding | Regulate the integrity of the podocyte actin cytoskeleton and for the regulation of podocyte cell migration. | Alport syndrome (AS), eosinophilic esophagitis, autism, schizophrenia | | https://www.protein |
| Synaptopodin-2, Genethonin-2, Myopodin | SYNPO2 | 4 | 1093 | Bundling | Participate in signaling pathways between the Z-disc and the nucleus. | Urothelial cancer, prostate cancer, bladder cancer. | 410-563 | https://www.protein |

| | | | | | | | | |
|---|---|---|---|---|---|---|---|---|
| Synaptopodin 2-like protein | SYNPO2L | 10 | 977 | Bundling | Required for cardiac and skeletal muscle development. | Atrial fibrillation, myofibrillar myopathies (MFMs), cardiomyopathy and contractile dysfunction | | https://www.protein |
| Smoothelin | SMTN | 22 | 917 | Bundling, Scaffolding | Implicated in cell contraction and mediate the interaction between actin filaments and other components of the cytoskeleton. | Atherosclerosis and restenosis, cardiac hypertrophy, essential hypertension, cerebral infarction, smooth muscle hamartoma, prostate cancer, glomus tumor, urinary bladder carcinoma, colorectal adeno-carcinoma, smooth muscle myopathy | Multiple domains | https://www.protein |
| Smoothelin-like protein 2 | SMTNL2 | 17 | 461 | Stabilization | Regulate actin dynamics during epithelial morphogenesis and the developmental control of the cellular cortex. | Rheumatoid arthritis, diabetes, middle cerebral artery occlusion, non-small cell lung cancer, Brachmann-Cornelia de Lange syndrome. | | https://www.protein |
| Myc box-dependent-interacting protein 1, Amphiphysin II, Amphiphysin-like protein, Box-dependent myc-interacting protein 1, Bridging integrator 1 | BIN1, AMPHL | 2 | 593 | Stabilization | Involved in endocytosis, actin cytoskeletal organization, transcription, and stress responses. | Alzheimer's disease, heart failure (HF), malignant arrhythmia, breast, colon, prostate and lung cancers, hepatocarcinoma, neuroblastoma, centronuclear myopathy (CNM) and myotonic dystrophy (DM), ventricular arrhythmia | Bin/Amphiphysin/Rvs (BAR) domain | https://www.protein |

| | | | | | | | | |
|---|---|---|---|---|---|---|---|---|
| Catenin alpha-1, Alpha E-catenin, Cadherin-associated protein, Renal carcinoma antigen NY-REN-13 | CTNNA1 | 5 | 906 | Bundling, Anchoring | Guide the establishment of classical epithelial cell polarity and contribute to the control of migration, growth, and differentiation. | Gastrointestinal cancer, colon cancer, dilated cardiomyopathy, cardiac injury, hereditary diffuse gastric cancer (HDGC), invasive lobular breast cancer (LBC), colitis. | C-terminal | https://www.protein |
| EH domain-binding protein 1 | EHBP1, KIAA0903, NACSIN | 2 | 1231 | Anchoring | Link endosomes to the actin cytoskeleton. | Prostate cancer, colorectal cancer, pulmonary arterial hypertension, rectal cancer. | Calponin homology (CH) domain | https://www.protein |
| Ras GTPase-activating-like protein IQGAP1, p195 | IQGAP1, KIAA0051 | 15 | 1657 | Cross-linking | Regulate mitogen-activated protein kinase (MAPK) signaling, Ca2+/calmodulin signaling, cell-cell adhesion, β-catenin-mediated transcription and microbial invasion. | Colorectal cancer, glioma, lung cancer, head and neck squamous cell, astrocytoma, breast cancer, gastric cancer, ovarian cancer, metastatic melanoma. | Calponin homology (CH) domain | https://www.protein |
| Ras GTPase-activating-like protein IQGAP2 | IQGAP2 | 5 | 1575 | Cross-linking | Predicted to bind to F-actin due to sequence similarity. | Carcinoma. | Calponin homology (CH) domain | https://www.protein |
| Ras GTPase-activating-like protein IQGAP3 | IQGAP3 | 1 | 1631 | Cross-linking | Link the activation of Rac1 and Cdc42 with the cytoskeletal architectures during neuronal morphogenesis. | Colorectal cancer, ovarian cancer, lung cancer, gastric cancer, colorectal cancer, hepatocellular carcinoma, breast cancer, clear cell renal cell carcinoma, bladder cancer, pancreatic cancer. | Calponin homology (CH) domain | https://www.protein |

| | | | | | | | | |
|---|---|---|---|---|---|---|---|---|
| LIM domain-binding protein 3， Protein cypher, Z-band alternatively spliced PDZ-motif protein | LDB3, KIAA0613, ZASP | 10 | 727 | Monomer binding, F-actin binding | Effect on the core structure of the Z-discs in skeletal muscle. | Markesbery disease, myofibrillar myopathies (MFM), hypertrophic cardiomyopathy (HCM), dilated cardiomyopathy (DCM), arrhythmogenic right ventricular cardiomyopathy (ARVC), left ventricular noncompaction (LVNC). | ABD between PDZ and LIM | https://www.protein |
| Myocardin-related transcription factor A (MRTF-A), MKL/myocardin-like protein 1, Megakaryoblastic leukemia 1 protein, Megakaryocytic acute leukemia protein | MRTFA, KIAA1438, MAL, MKL1 | 22 | 931 | Monomer binding | Transcription coactivator that associates with the serum response factor. | Acute megakaryoblastic leukemia, Immunodeficiency 66 (IMD66) | RPEL motif | https://www.protein |
| Protein MTSS1, Metastasis suppressor YGL-1, Metastasis suppressor protein 1, Missing in metastasis protein | MTSS1, KIAA0429, MIM | 8 | 755 | Monomer binding, Bundling, Scaffolding | Regulate cell morphology, motility, metastasis. Act as a scaffold protein that interacts with Rac, actin and actin-associated proteins to modulate lamellipodia formation. | Prostate cancer, breast cancer, acute myeloid leukemia, pancreatic cancer, intrahepatic cholangiocarcinoma, pancreatic ductal adenocarcinoma, bladder uroepithelium cell carcinoma, hepatocellular carcinoma, colorectal cancer, esophageal cancer, tongue squamous cellular carcinoma, gastric cancer, melanomas. | WH2 domain, IRSp53/MIM domain | https://www.protein |

| | | | | | | | | |
|---|---|---|---|---|---|---|---|---|
| Rab effector MyRIP, Exophilin-8, Myosin-VIIa- and Rab-interacting protein, Synaptotagmin-like protein lacking C2 domains C (SlaC2-c) | MYRIP, SLAC2C | 3 | 859 | Anchoring | Regulate Weibel-Palade body trafficking and exocytosis. | Left ventricular hypertrophy, hepatocellular carcinoma. | Slac2-a (400-590) and Slac2-c (670-856) | https://www.protein |
| Nebulin | NEB | 2 | 6669 | Nucleation, Stabilization | Form composite thin filaments in the skeletal muscle sarcomere. | Core-rod myopathy, distal myopathy | Nebulin repeats | https://www.protein |
| Nebulin-related-anchoring protein (N-RAP) | NRAP | 10 | 1730 | Anchoring | Act as an organizing center for the initial recruitment and assembly of sarcomeric actin filaments and Z-discs. | Dilated cardiomyopathy (DCM), | Nebulin repeats | https://www.protein |
| Vinculin, Metavinculin (MV) | VCL | 10 | 1134 | Capping, Anchoring | Interact with F-actin both in recruitment of actin filaments to the growing focal adhesions and also in capping of actin filaments to prevent actin polymerization. | Chagas cardiomyopathy, dilated cardiomyopathy, hypertrophic cardiomyopathy, left ventricular assist device (LVAD). | Vinculin tail domain (Vt) | https://www.protein |
| Cysteine and glycine-rich protein 3, Cardiac LIM protein, Cysteine-rich protein 3 (CRP3), LIM domain protein, cardiac, Muscle LIM protein | CSRP3, CLP, MLP | 11 | 194 | Bundling | Facilitate filopodia formation and increasing growth cone motility. Act as a mechanical stress sensor, entering in nuclei and modulating gene expression. | Dilated cardiomyopathy, hypertrophic cardiomyopathy, ischemic cardiomyopathy, Nemaline myopathy (NM), neuromuscular disorder facioscapulohumeral muscular dystrophy (FSHD). | LIM motif | https://www.protein |
| Utrophin, Dystrophin-related protein 1 (DRP-1) | UTRN, DMDL, DRP1 | 6 | 3433 | Anchoring | Perform 'spacer' or 'shock absorber' role as dystrophin in mature muscle tissues. | Duchenne muscular dystrophy (DMD), Becher muscular dystrophy. | CH1(31-135), CH2(150-255) | https://www.protein |
| Growth arrest-specific 7 (GAS-7) | GAS7, KIAA0394 | 17 | 476 | Polymerization, Cross-linking | Mediate reorganization of microfilaments and induce the formation of extended cellular processes. | Glaucoma, Alzheimer disease, schizophrenia. | C terminal | https://www.protein |

### Vedula et al 2023 selected list of actin binding proteins (adapted from Gao and Nakamua 2022)

| | | | | | | | | |
|---|---|---|---|---|---|---|---|---|
| Calicin | CCIN | 9 | 588 | F-actin binding | Target of calicin at the subacrosomal space of round spermatids, and that its ability to form homomultimers contributes to the formation of a rigid calyx. | | | https://www.protein |
| Ermin, Juxtanodin (JN) | ERMN, KIAA1189 | 2 | 284 | | Induce the formation of numerous cell protrusions and a pronounced change in cell morphology. | Multiple Sclerosis (MS) | 265-284 | https://www.protein |
| FK506-binding protein 15 (FKBP-15), 133 kDa FK506-binding protein (133 kDa FKBP, FKBP-133), WASP- and FKBP-like protein (WAFL) | FKBP15, KIAA0674 | 9 | 1219 | | Involve in the transport of early endosomes at the level of transition between microfilament-based and microtubule-based movement. | Inflammatory bowel disease (IBD). | | https://www.protein |
| Growth arrest-specific protein 2 (GAS-2) | GAS2 | 11 | 313 | Cross-linking | Cross-link microtubule and actin cytoskeletons. | Liver cancer, leukemia, recurrent colorectal cancer, prostate cancer, breast cancer, lung adenocarcinoma. | CH (34-156) | https://www.protein |
| GAS2-like protein 1, GAS2-related protein on chromosome 22, Growth arrest-specific protein 2-like 1 | GAS2L1, GAR22 | 22 | 681 | Cross-linking | Cross-link microtubule and actin cytoskeletons. | Acute myeloid leukemia, meningioma. | CH (27-148) | https://www.protein |
| GAS2-like protein 2, GAS2-related protein on chromosome 17, Growth arrest-specific 2-like 2 | GAS2L2, GAR17 | 17 | 880 | Cross-linking | Cross-link microtubule and actin cytoskeletons. | Cancer with painful bone metastases, Ciliary dyskinesia, primary, 41 (CILD41). | CH (32-159) | https://www.protein |
| GAS2-like protein 3, Growth arrest-specific 2-like 3 | GAS2L3 | 12 | 694 | Cross-linking | Cross-link microtubule and actin cytoskeletons. | Glioma | CH (48-168) | https://www.protein |

| | | | | | | | | |
|---|---|---|---|---|---|---|---|---|
| Kelch-like protein 2, Actin-binding protein Mayven | KLHL2 | 4 | 593 | Scaffolding | Translocate along axonal processes and involved in the dynamic organization of the actin cytoskeleton in brain cells. | Renal cell carcinoma, acute myeloid leukemia (AML), ovarian cancer, familial hyperkalemic hypertension, acute peanut allergic reaction, septic-shock-associated acute kidney injury, multiple myeloma. | Kelch repeats | https://www.protein |
| Kelch-like protein 20, Kelch-like ECT2-interacting protein, Kelch-like protein X | KLHL20, KHLHX, KLEIP | 1 | 609 | Scaffolding | Involved in anterograde Golgi to endosome transport. Regulate corneal epithelial integrity. | Colon cancer | Kelch repeats | https://www.protein |
| Actin-binding protein IPP, Intracisternal A particle-promoted polypeptide (IPP), Kelch-like protein 27 | IPP, KLHL27 | 1 | 584 | Anchoring | Regulate cell adhesion. | Breast cancer | Kelch repeats | https://www.protein |
| Protein ITPRID2, Cleavage signal-1 protein (CS-1), ITPR interacting domain-containing 2, Ki-ras-induced actin-interacting protein, Sperm-specific antigen 2 | ITPRID2, CS1, KIAA1927, KRAP, SSFA2 | 2 | 1259 | Scafolding | Regulate energy metabolism. | Colorectal cancer, pancreatic cancer, small cell lung cancer, obesity and diabetes, lung squamous cell cancer, chronic lymphocytic leukemia | | https://www.protein |
| Melanophilin, Exophilin-3, Slp homolog lacking C2 domains a (SlaC2-a), Synaptotagmin-like protein 2a | MLPH, SLAC2A | 2 | 600 | Scafolding | Involved in in actin-based melanosome transport. | Rectal cancer, prostate cancer, allergic and inflammatory disease, Inherited diseases of pigmentation. | 401-590 | https://www.protein |
| Pleckstrin homology domain-containing family H member 2 | PLEKHH2, KIAA2028 | 2 | 1493 | Stabilization | Slow down actin depolymerization. | Diabetic nephropathy, schizophrenia. | FERM domain | https://www.protein |

| | | | | | | | | |
|---|---|---|---|---|---|---|---|---|
| SH2B adaptor protein 1, Pro-rich, PH and SH2 domain-containing signaling mediator (PSM), SH2 domain-containing protein 1B | SH2B1, KIAA1299, SH2B | 16 | 756 | Cross-linking | Regulate cultured cell morphology, motility and adhesion. | Obesity, leptin and insulin resistance, non-small cell lung cancer (NSCLC), esophageal cancer, gastric cancer, colorectal cancer, oropharyngeal cancer | 150–200 and 615–670. | https://www.protein |
| Annexin A1, Annexin I, Annexin-1, Calpactin II, Calpactin-2, Chromobindin-9, Lipocortin I, Phospholipase A2 inhibitory protein | ANXA1, ANX1, LPC1 | 9 | 346 | Bundling, Anchoring | Mediate the Ca2+-dependent interaction between phagosomes and the actin cytoskeleton. | Hepatocellular carcinoma, lung cancer, melanoma colorectal cancer, and pancreatic cancer, esophageal cancer, prostate cancer, cervical cancer, B-cell non-Hodgkin's lymphomas, larynx cancer, nasopharyngeal carcinoma (NPC) and oral squamous cell carcinoma, multiple sclerosis, Alzheimer's disease, atherosclerosis, coronary artery disease, myocardial infarction, diabetic nephrophthy, stroke. | IRI motif | https://www.protein |
| Annexin A2, Annexin II, Annexin-2, Calpactin I heavy chain, Calpactin-1 heavy chain, Chromobindin-8, Lipocortin II, Placental anticoagulant protein IV (PAP-IV, Protein I), p36 | ANXA2, ANX2, ANX2L4, CAL1H, LPC2D | 15 | 339 | Bundling, Anchoring | Maintain the plasticity of the dynamic membrane-associated actin cytoskeleton. | Breast cancer, glioblastoma, renal cell carcinoma, hepatocellular carcinoma, colorectal cancer, lung cancer, bacteria, fungus, virus infection, acute and chronic inflammatory disorder. | IRI motif | https://www.protein |

Vedula et al 2023 selected list of actin binding proteins (adapted from Gao and Nakamua 2022)

| | | | | | | | | |
|---|---|---|---|---|---|---|---|---|
| Annexin A5, Anchorin CII, Annexin V, Annexin-5, Calphobindin I (CBP-I), Endonexin II, Lipocortin V, Placental anticoagulant protein 4 (PP4), Placental anticoagulant protein I (PAP-I), Thromboplastin inhibitor, Vascular anticoagulant-alpha (VAC-alpha) | ANXA5, ANX5, ENX2, PP4 | 4 | 320 | Anchoring | | Hepatocarcinoma, breast cancer, cervical carcinoma, gastric cancer, nasopharyngeal carcinoma, colorectal cancer, pancreatic cancer, bladder cancer, prostate cancer, squamous cell carcinoma, glioma, sarcoma, thyroid cancer. | | https://www.protein |
| Annexin A6, 67 kDa calelectrin, Annexin VI, Annexin-6, Calphobindin-II (CPB-II), Chromobindin-20, Lipocortin VI, Protein III, p68, p70 | ANXA6, ANX6 | 5 | 673 | Anchoring | Regulate microfilament architecture, in response to calcium signals in neurites. | Breast cancer, esophageal adenocarcinoma. | | https://www.protein |
| Annexin A8, Annexin VIII, Annexin-8, Vascular anticoagulant-beta, VAC-beta. | ANXA8, ANX8 | 10 | 327 | Anchoring | Regulate late endosomes organization. | Gastric carcinoma. | | https://www.protein |
| Girdin, Akt phosphorylation enhancer (APE), Coiled-coil domain containing protein 88A, G alpha-interacting vesicle-associated protein (GIV), Girders of actin filament, Hook-related protein 1 (HkRP1) | CCDC88A, APE, GRDN, KIAA1212 | 2 | 1871 | Cross-linking, Bundling, Anchoring | Form actin bundles, and anchor the bundles to the plasma membrane at the cortical region of the cells. Facilitate lamellar protrusion, cell migration, adhesion, and invasion. | Pancreatic cancer, breast cancer, lung cancer, cervical carcinoma, colorectal cancer, gastric cancer, hepatocellular carcinoma, acute myocardial infarction, Alzheimer's disease (AD). | C-terminal 2 | https://www.protein |

| | | | | | | | | |
|---|---|---|---|---|---|---|---|---|
| Lymphocyte-specific protein 1, 47 kDa actin-binding protein, 52 kDa phosphoprotein (pp52), Lymphocyte-specific antigen WP34 | LSP1, WP34 | 11 | 339 | Bundling | Regulate focal adhesion dynamics, cell migration and phagocytosis. | Hodgkin's disease, breast cancer, Sjögren's syndrome, acute lung inflammation, rheumatoid arthritis, hepatocellular carcinoma, schizophrenia, carotid paragangliomas. | C-terminal basic domain | https://www.protein |
| [F-actin]-monooxygenase MICAL1, Molecule interacting with CasL protein 1 (MICAL1), NEDD9-interacting protein with calponin homology and LIM domains | MICAL1, MICAL, NICAL | 6 | 1067 | PT modification, Depolymerization | Directly oxidizes Met44 and Met47 of F-actin into methionine-R-sulfoxides resulting in F-actin disassembly. | Different cancers, diabetic nephropathy, blood brain barrier dysfunction, muscular dystrophy, liver disease, susceptibility to infection, epilepsy, Alzheimer’s disease, aging, deafness, wide-ranging neurological disorders, skeletal abnormalities, and obesity. | FAD | https://www.protein |
| [F-actin]-monooxygenase MICAL2, Molecule interacting with CasL protein 2 (MICAL-2) | MICAL2, KIAA0750, MICAL2PV1, MICAL2PV2 | 11 | 1124 | PT modificatio, Depolymerization | Induce loss of actin stress fibers and the generation of actin-rich protrusions. | Colorectal, breast, gastric, prostate, bladder Cancer, lung adenocarcinoma, laryngeal squamous cell carcinoma. | FAD | https://www.protein |
| [F-actin]-monooxygenase MICAL3, Molecule interacting with CasL protein 3, MICAL-3. | MICAL3, KIAA0819, KIAA1364 | 22 | 2002 | PT modificatio, Depolymerization | Directly oxidizes Met44 and Met47 of F-actin into methionine-R-sulfoxides resulting in F-actin disassembly. | | | https://www.protein |
| Methionine-R-sulfoxide reductase B1 (MsrB1), Selenoprotein X (SelX) | MSRB1, SEPX1 | 16 | 116 | PT modificatio, Polymerization | Regulate actin assembly by reducing methionine (R)-sulfoxide mediated by MICALs on actin, thereby promoting polymerization. | | | https://www.protein |

| | | | | | | | | |
|---|---|---|---|---|---|---|---|---|
| Methionine-R-sulfoxide reductase B2, mitochondrial (MsrB2) | MSRB2, CBS-1, MSRB | 10 | 182 | PT modification, Polymerization | Regulate actin assembly by reducing methionine (R)-sulfoxide mediated by MICALs on actin, thereby promoting polymerization. | Parkinson's disease, Alzheimer's disease. | | https://www.protein |
| N-alpha-acetyltransferase 80 (HsNAAA80) N-acetyltransferase 6, Protein fusion-2 (Protein fus-2) | NAA80, FUS2, NAT6 | 3 | 286 | PT modification, Polymerization | Regulate actin assembly and cell motility. | Glioblastoma, low-grade gliomas (LGG), kidney cancers. | | https://www.protein |
| Prolactin-inducible protein, gross cystic disease fluid protein 15, secretory actin-binding protein (SABP, gp17) | PIP, GCDFP15, GPIP4 | 7 | 146 | Scaffolding | Regulate antitumor immunity and metastasis. | Lung cancer, breast cancer, Sjogren's Syndrome, metastatic carcinoma, mucinous carcinoma of skin, uterine leiomyomas. | | https://www.protein |
| Plectin (PCN, PLTN), Hemidesmosomal protein 1 (HD1), Plectin-1 | PLEC, PLEC1 | 8 | 4684 | Cross-linking | Connect intermediate filaments with microtubules and actin filaments. Form desmosomes and hemidesmosomes. | Pancreatic ductal carcinoma (PDAC), pancreatic, lung, esophageal, stomach, ovarian, and breast cancer, head and neck squamous cell carcinoma (HNSCC), oral squamous cell carcinoma (OSCC), hepatocellular carcinoma (HCC), epidermolysis bullosa simplex (EBS). | CH1 (179 – 282), CH2 (295 – 400) | https://www.protein |
| Shootin-1 | SHTN1, KIAA1598 | 10 | 631 | Cross-linking | Mediate actin filament retrograde flow and L1-CAM in axonal growth cones. | Age-related macular degeneration (AMD) | WH2 | https://www.protein |

| | | | | | | | | |
|---|---|---|---|---|---|---|---|---|
| Angiomotin | AMOT, KIAA1071 | X | 1084 | Anchoring, Scaffolding | Regulate tube formation and migration of endothelial cells and the regulation of tight junctions, polarity, and epithelial-mesenchymal transition in epithelial cells. | Breast cancer, osteosarcoma, prostate cancer, head and neck squamous cell carcinoma (HNSCC), hepatic carcinoma, renal cell cancer, ovarian cancer, lung cancer. | N-terminal | https://www.protein |
| CLIP-associating protein 1, Cytoplasmic linker-associated protein 1, Multiple asters homolog 1, Protein Orbit homolog 1 (hOrbit1) | CLASP1, KIAA0622, MAST1 | 2 | 1538 | Cross-linking | Link plus ends of growing microtubules and actin filaments. | Prostate cancer. | Middle serine-arginine rich motif, dis1/TOG motif | https://www.protein |
| CLIP-associating protein 2, Cytoplasmic linker-associated protein 2, Protein Orbit homolog 2 (hOrbit2) | CLASP2, KIAA0627 | 3 | 1294 | Cross-linking | Link plus ends of growing microtubules and actin filaments. | Bladder tumor, endothelial inflammation and lung injury. | Middle serine-arginine rich motif, dis1/TOG motif | https://www.protein |
| Dixin, Coiled-coil protein DIX1 (Coiled-coil-DIX1), DIX domain-containing protein 1 | DIXDC1, CCD1, KIAA1735 | 11 | 683 | Cross-linking | Interact with gamma-tubulin. Regulate excitatory neuron dendrite development and synapse function in the cortex. | Autism spectrum disorders (ASDs), acute myeloid leukemia, cerebral ischemia/reperfusion injury, glioma, prostate cancer, gastric cancer, non-small-cell lung cancer, hepatocellular carcinoma, oral squamous cell carcinoma, colon cancer, retinoblastoma, pancreatic ductal adenocarcinoma, bladder cancer | 127-300 | https://www.protein |

| | | | | | | | | |
|---|---|---|---|---|---|---|---|---|
| Dystonin, 230 kDa bullous pemphigoid antigen, 230/240 kDa bullous pemphigoid antigen, Bullous pemphigoid antigen 1 (BPA), Dystonia musculorum protein, Hemidesmosomal plaque protein | DST, BP230, BP240, BPAG1, DMH, DT, KIAA0728 | 6 | 7570 | Cross-linking | Link intermediate filaments, actin and microtubule cytoskeleton networks. | Hereditary sensory and autonomic neuropathy type 6 (HSAN6), Parkinson's disease, schizophrenia, Dystonia musculorum | CH1 (35-138), CH2 (151-255) | https://www.protein |
| Protein FRG1, FSHD region gene 1 protein | FRG1 | 4 | 258 | Bundling | Involved in muscle structure and integrity. | Prostate cancer, colorectal cancer, lung adenocarcinoma, Facioscapulohumeral muscular dystrophy (FSHD), gastric, colon and oral cavity tumor. | Fascin-like domain? | https://www.protein |
| Kelch repeat and BTB domain-containing protein 13 | KBTBD13 | 15 | 458 | | Involved in muscle relaxation. An adapter of a BCR (BTB-CUL3-RBX1) E3 ubiquitin ligase complex. | Nemaline myopathy, rod-core myopathy. | | https://www.protein |
| Fimbacin; Leucine zipper protein 1 | LUZP1 | 1 | 1076 | Cross-linking | Involved in actin-dependent centrosome to basal body conversion. Control cell division, migration and invasion. | Townes-Brocks Syndrome (TBS), glioma, colorectal cancer, cardiovascular malformations and cardiomyopathy, neural tube closure defect (NTD). | 400-500 | https://www.protein |
| Protein kinase C and casein kinase substrate in neurons protein 2, Syndapin-2, Syndapin-II (SdpII) | PACSIN2 | 22 | 486 | Scaffolding | Regulate the morphogenesis and endocytosis of caveolae. | Diabetic kidney disease, virus infection, acute lymphoblastic leukemia. | F-BAR domain | https://www.protein |
| Pannexin-1 | PANX1, MRS1 | 11 | 426 | Anchoring, Scaffolding | Restrain human skin fibroblast motility, migration, and cell surface actin dynamics. | Renal ischemia/reperfusion injury, epilepsy, stroke, migraine with aura, chronic pain. | C-terminal | https://www.protein |

Vedula et al 2023 selected list of actin binding proteins (adapted from Gao and Nakamua 2022)

| | | | | | | | | |
|---|---|---|---|---|---|---|---|---|
| Refilin-A, Regulator of filamin protein A (RefilinA) | RFLNA, FAM101A | 12 | 216 | Bundling | Regulate lamellipodium protrusion dynamics. | Spondylocarpotarsal synostosis syndrome. | | https://www.protein |
| Refilin-B, Regulator of filamin protein B (RefilinB) | RFLNB | 17 | 214 | Bundling | Regulate apical perinuclear Actin reorganization in early steps of epithelial mesenchymal transition (EMT). | Severe preeclampsia. | | https://www.protein |
| SH3 and multiple ankyrin repeat domains protein 3 (Shank3), Proline-rich synapse-associated protein 2 (ProSAP2) | SHANK3, KIAA1650, PROSAP2, PSAP2 | 22 | 1731 | Scaffolding | Regulate dendritic spine morphology in neurons and autism-linked phenotypes in vivo. | Parkinson disease, Phelan-McDermid syndrome, Alzheimer's disease, autism spectrum disorders (ASDs), schizophrenia (SCZ), a Rett syndrome-like phenotype, and intellectual disability (ID). | Shank/ProSAP N-terminal (SPN) domain | https://www.protein |
| Troponin I, slow skeletal muscle | TNNI1 | 1 | 187 | Stabilization, Scaffolding | Regulate muscle contraction. Calcium-receptive protein for the Calcium -sensitive contraction in striated muscle. | Autosomal dominant proximal arthrogryposis, cardiomyopathy. | 138-148 | https://www.protein |
| Troponin I, fast skeletal muscle, Troponin I, fast-twitch isoform | TNNI2 | 11 | 182 | ? | Regulate muscle contraction. Calcium-receptive protein for the Calcium -sensitive contraction in striated muscle. | Arthrogryposis, distal, 2B1 (DA2B1). | | https://www.protein |
| Troponin I, cardiac muscle, Cardiac troponin I | TNNI3, TNNC1 | 19 | 210 | ? | Regulate muscle contraction. Calcium-receptive protein for the Calcium -sensitive contraction in striated muscle. | Cardiomyopathy, familial hypertrophic 7 (CMH7), Cardiomyopathy, familial restrictive 1 (RCM1), Cardiomyopathy, dilated 2A (CMD2A), Cardiomyopathy, dilated 1FF (CMD1FF). | | https://www.protein |
| Troponin T, slow skeletal muscle, TnTs, Slow skeletal muscle troponin T, TnT. | TNNT1, TNT | 19 | 278 | ? | Regulate muscle contraction. Calcium-receptive protein for the Calcium -sensitive contraction in striated muscle. | Nemaline myopathy 5 (NEM5). | | https://www.protein |

| | | | | | | | | |
|---|---|---|---|---|---|---|---|---|
| Troponin T, cardiac muscle (TnTc), Cardiac muscle troponin T (cTnT) | TNNT2 | 1 | 298 | Stabilization, Scaffolding | Regulate muscle contraction. Calcium-receptive protein for the Calcium -sensitive contraction in striated muscle. | Dilated cardiomyopathy, hypertrophic cardiomyopathy, colorectal cancer, myotonic dystrophy type I, feline cardiomyopathy, myocardial diastolic dysfunction, sporadic hypertrophic cardiomyopathy. | C-terminal | https://www.protein |
| Troponin T, fast skeletal muscle, TnTf, beta-TnTF, Fast skeletal muscle troponin T, fTnT. | TNNT3 | 11 | 269 | ? | Regulate muscle contraction. Calcium-receptive protein for the Calcium -sensitive contraction in striated muscle. | Arthrogryposis, distal, 2B2 (DA2B2). | | https://www.protein |
| Troponin C, slow skeletal and cardiac muscles, TN-C. | TNNC1, TNNC | 3 | 161 | ? | Regulate muscle contraction. Calcium-receptive protein for the Calcium -sensitive contraction in striated muscle. | Cardiomyopathy, dilated 1Z (CMD1Z), Cardiomyopathy, familial hypertrophic 13 (CMH13). | | https://www.protein |
| Troponin C, skeletal muscle | TNNC2 | 20 | 160 | ? | Regulate muscle contraction. Calcium-receptive protein for the Calcium -sensitive contraction in striated muscle. | Distal arthrogryposis (DA). | | https://www.protein |
| Tensin-1 | TNS1, TNS | 2 | 1735 | Cross-linking, Capping | Maintain cellular structure and signal transduction and anchor actin filaments at the focal adhesion. | Cystic kidney diseases, colorectal cancer, acute myeloid leukemia, oral cleft, prostate cancer, kidney cancer, mitral valve prolapse (MVP), chronic obstructive lung disease (COPD), asthma with hay fever phenotype. | ABD Ia, ABD Ib, ABD II | https://www.protein |

| | | | | | | | | |
|---|---|---|---|---|---|---|---|---|
| Tensin-2, C1 domain-containing phosphatase and tensin homolog (C1-TEN), Tensin-like C1 domain-containing phosphatase | TNS2, KIAA1075, TENC1 | 12 | 1409 | Anchoring | Permit Rho-mediated actomyosin contraction and remodeling of collagen fibers. | Pancreatic cancer, kidney cancer, glomerular disease, gastric cancer, sensitive nephrotic syndrome. | ABD Ia, ABD Ib, ABD II | https://www.protein |
| Tensin-3, Tensin-like SH2 domain-containing protein 1, Tumor endothelial marker 6 | TNS3, TEM6, TENS1, TPP | 7 | 1445 | Anchoring | Reorganize actin fiber and regulate cell migration. | Pancreatic cancer, melanoma, breast cancer, gastric cancer, colorectal cancer, hepatocellular carcinoma, lung adenocarcinoma, kidney cancer. | ABD Ia, ABD Ib, ABD II | https://www.protein |
| Titin, Connectin, Rhabdomyosarcoma antigen MU-RMS-40.14 | TTN | 2 | 34350 | Cross-linking | Contribute to muscle compliance, contraction, structural stability, and signaling. | Titinopathies, heart failure, ischemic and non-ischemic cardiomyopathy, dilated cardiomyopathy, tibialis muscular dystrophy, hypertrophic cardiomyopathy, centronuclear myopathy, Hereditary myopathies, Acquired titin diseases, Chronic Obstructive Pulmonay Disease (COPD), disuse atrophy. | N2A domain | https://www.protein |
| Dihydropyrimidinase-related protein 1, GRP-1, Collapsin response mediator protein 1, CRMP-1, Inactive dihydropyrimidinase, Unc-33-like phosphoprotein 3, ULIP-3. | CRMP1, DPYSL1, ULIP3 | 4 | 572 | Scaffolding | Regulate remodeling of the cytoskeleton by dissociating FLNA from F-actin during the axon guidance process. | Schizophrenia, Epilepsy. | | https://www.proteinatlas.org/ENSG00000072832-CRMP1 |

| | | | | | | | | |
|---|---|---|---|---|---|---|---|---|
| Dihydropyrimidinase-related protein 2 (DRP-2), Collapsin response mediator protein 2 (CRMP-2), N2A3, Unc-33-like phosphoprotein 2 (ULIP-2) | DPYSL2, CRMP2, ULIP2 | 8 | 572 | Scaffolding | Promote microtubule assembly and Numb-mediated endocytosis | Alzheimer's disease (AD), Neurodegenerative disease, amyotrophic lateral sclerosis (ALS), Huntington's Disease, neuropathic pain, and Batten disease, Parkinson's disease, lung cancer. | | https://www.protein |
| Dihydropyrimidinase-related protein 3 (DRP-3), Collapsin response mediator protein 4 (CRMP-4), Unc-33-like phosphoprotein 1 (ULIP-1) | DPYSL3, CRMP4, DRP-3, ULIP, ULIP1 | 5 | 570 | Bundling | Inhibit the cell migration and maintain rib-like actin-structures in lamellipodia. Promote dendritic growth and maturation. | Amyotrophic lateral sclerosis (ALS), Autism spectrum disorders (ASD), Parkinson's disease, Alzheimer's disease | C-terminal | https://www.protein |
| Dihydropyrimidinase-related protein 5 (DRP-5), CRMP3-associated molecule (CRAM), Collapsin response mediator protein 5 (CRMP-5), UNC33-like phosphoprotein 6 (ULIP-6) | DPYSL5, CRMP5, ULIP6 | 2 | 564 | Scaffolding | Regulate growth cone development and neurite outgrowth. | Paraneoplastic neurological syndromes (PNS), Alzheimer's disease, small-cell lung cancer, thymoma. | C-terminal | https://www.protein |
| Cysteine and glycine-rich protein 2, Cysteine-rich protein 2 (CRP2), LIM domain only protein 5 (LMO-5), Smooth muscle cell LIM protein (SmLIM) | CSRP2, LMO5, SMLIM | 12 | 193 | Bunding | Contribute to the assembly and/or maintenance of the invadopodium actin backbone. | Colorectal cancer, acute lymphoblastic leukemia, hepatocellular carcinoma. | | https://www.protein |
| ABI gene family member 3, New molecule including SH3, Nesh | ABI3, NESH | 17 | 366 | Stabilization | Regulate dendritic spine morphogenesis and synapse formation. | Alzheimer's disease and Nasu-Hakola disease. | N-terminal | https://www.protein |
| Neuroblast differentiation-associated protein AHNAK, Desmoyokin | AHNAK, PM227 | 11 | 5890 | Bundling | Stabilize muscle contractility. | Pulmonary tumorigenesis | C-terminal | https://www.protein |

| | | | | | | | | |
|---|---|---|---|---|---|---|---|---|
| LIM domain-containing protein ajuba | AJUBA, JUB | 14 | 538 | Scaffolding | Scaffold many protein to regulate several cellular processes such migration and proliferation. | Hepatocellular carcinoma | | https://www.protein |
| T-complex protein 1 subunit epsilon, TCP-1-epsilon, CCT-epsilon T-complex protein 1 subunit theta, TCP-1-theta, CCT-theta, Renal carcinoma antigen NY-REN-15 | CCT5, CCTE, KIAA 0098 CCT8, C21orf112, CCTQ, KIAA0002 | 5<br>21 | 541<br>548 | Folding of actin | Molecular chaperone complex involved in folding of actin and tubulin. | Neuropathy. | | https://www.protein |
| Prefoldin (Prefoldin subunit 1, Prefoldin subunit 2, Prefoldin subunit 3, Prefoldin subunit 4, Prefoldin subunit 5, Prefoldin subunit 6) | PFDN1, PFD1 PFDN2, PFD2 PFDN3, VBP1 PFDN4, PFD4 PFDN5, PFD5 PFDN6, PFD6, HKE2 | 5<br>1<br>X<br>20<br>12<br>6 | 122<br>154<br>197<br>134<br>154<br>129 | Folding of actin | Target premature nascent actin to cytosolic chaperonin. | Alzheimer's disease, Parkinson's disease, and Huntington's disease. | 1-145 | https://www.proteinatlas.org/ENSG00000113068-PFDN1<br>https://www.proteinatlas.org/ENSG00000143256-PFDN2<br>https://www.proteinatlas.org/ENSG00000155959-VBP1<br>https://www.proteinatlas.org/ENSG00000101132-PFDN4<br>https://www.proteinatlas.org/ENSG00000123349-PFDN5<br>https://www.proteinatlas.org/ENSG00000204220-PFDN6 |
| Carcinoembryonic antigen-related cell adhesion molecule 1, Biliary glycoprotein 1, BGP-1, CD66a | CEACAM1, GP, BGP1 | 19 | 526 | Anchoring, Scaffolding | Mediate homophilic cell adhesion. | Inflammatory bowel disease. | Cytoplasmic domain (453-526) | https://www.protein |
| Cingulin | CGN, KIAA1319 | 1 | 1197 | Anchoring | Regulate formation and regulation of the tight junction. | Disease related to tight junction: inflammatory bowel disease. | 101-294 | https://www.protein |
| Alpha-crystallin B chain, Heat shock protein beta-5, HspB5, Renal carcinoma antigen NY-REN-27, Rosenthal fiber component | CRYAB, CRYA2, HSPB5 | 11 | 175 | Scaffolding | Regulate the cell cycle, and reducing caspase-mediated apoptosis. | Myopathy, myofibrillar, 2 (MFM2), Cataract 16, multiple types (CTRCT16), Myopathy, myofibrillar, fatal infantile hypertonic, alpha-B crystallin-related (MFMFIH-CRYAB), Cardiomyopathy, dilated 1II (CMD1II). | ABD | https://www.protein |

| | | | | | | | | |
|---|---|---|---|---|---|---|---|---|
| Cytoskeleton-associated protein 5, Colonic and hepatic tumor overexpressed gene protein, Ch-TOG | CKAP5, KIAA0097(Xenopus laevis: XMAP215) | 11 | 2032 | Scaffolding | Regulate microtubule dynamics and microtubule organization. TOG/XMAP215 family. | | Demonstrated with Xenopus laevis XMAP215. | https://www.protein |
| Emerin | EMD, EDMD, STA | X | 254 | Capping | Stimulate actin polymerization in vitro by binding and stabilizing the pointed end. | Emery-Dreifuss muscular dystrophy 1, X-linked (EDMD1) | | https://www.protein |
| FYVE, RhoGEF and PH domain-containing protein 4, Actin filament-binding protein frabin, FGD1-related F-actin-binding protein, Zinc finger FYVE domain-containing protein 6 | FGD4, FRABP, ZFYVE6 | 12 | 766 | Cross-linking | Capable of changing cell shape and activating c-Jun N-terminal kinase. | Charcot-Marie-Tooth disease 4H (CMT4H). | N-terminal | https://www.protein |
| Heat shock protein beta-7, HspB7, Cardiovascular heat shock protein, cvHsp | HSPB7, CVHSP | 1 | 170 | Monomer binding | HSPB7 binds G actin and inhibits actin polymerization to regulate actin thin filament length in cardiac muscle. | Cardiocmyopathy. | | https://www.protein |
| Myosin-binding protein C, cardiac-type, MyBP-C, C-protein, cardiac muscle isoform | MYBPC3 | 11 | 1274 | | Regulate muscle contraction. | Cardiomyopathy, familial hypertrophic 4 (CMH4), Cardiomyopathy, dilated 1MM (CMD1MM), Left ventricular non-compaction 10 (LVNC10). | N-terminal | https://www.protein |
| AP-4 complex accessory subunit RUSC1, New molecule containing SH3 at the carboxy-terminus (Nesca), RUN and SH3 domain-containing protein 1 | RUSC1, NESCA | 1 | 902 | Scaffolding | Function as an adapter involved in neuronal vesicular transport. | | 231-316 | https://www.protein |

| | | | | | | | | |
|---|---|---|---|---|---|---|---|---|
| NCK-interacting protein with SH3 domain, 54 kDa VacA-interacting protein, 54 kDa vimentin-interacting protein (VIP54), 90 kDa SH3 protein interacting with Nck, AF3p21, Dia-interacting protein 1 (DIP-1), Diaphanous protein-interacting protein, SH3 adapter protein SPIN90, WASP-interacting SH3-domain protein (WISH), Wiskott-Aldrich syndrome protein-interacting protein | NCKIPSD, AF3P21, SPIN90 | 3 | 722 | Polymerization, Scaffolding | Involved in formation of branched actin networks and regulation of the actin cytoskeleton at the leading edge of cells. | Breast cancer, chromosomal aberration involving NCKIPSD/AF3p21. | 582-722 | https://www.protei[illegible] |
| Alpha-1-syntrophin, 59 kDa dystrophin-associated protein A1 acidic component 1, Pro-TGF-alpha cytoplasmic domain-interacting protein 1 (TACIP1), Syntrophin-1 | SNTA1, SNT1 | 20 | 505 | Scaffolding, Anchoring | Regulate intracellular localization and activity of various actin organizing signaling molecules. | Parkinson's disease (PD), long QT syndrome, breast cancer, neuromuscular junctions (NMJ), Duchenne muscular dystrophy (DMD), fukuyama muscular dystrophies (FMD), multisystemic disorder myotonic dystrophy type 1 (DM1), sudden infant death syndrome (SIDS). | 274-315, 449-505 | https://www.protei[illegible] |

| | | | | | | | | |
|---|---|---|---|---|---|---|---|---|
| Beta-1-syntrophin, 59 kDa dystrophin-associated protein A1 basic component 1 (DAPA1B, BSYN2), Syntrophin-2, Tax interaction protein 43 (TIP-43) | SNTB1, SNT2B1 | 8 | 538 | Scaffolding, Anchoring | Link various receptors to the actin cytoskeleton. Involved in synapse formation. | Duchenne muscular dystrophy (DMD) | PDZ domain | https://www.protein |
| Beta-2-syntrophin, 59 kDa dystrophin-associated protein A1 basic component 2, Syntrophin-3 (SNT3), Syntrophin-like (SNTL) | SNTB2, D16S2531E, SNT2B2, SNTL | 16 | 540 | Scaffolding, Anchoring | Link various receptors to the actin cytoskeleton. | Neuromuscular junctions (NMJ), type-2 diabetes mellitus or non-insulin-dependent diabetes mellitus (NIDDM), | PDZ domain | https://www.protein |
| Testin, TESS | TES | 7 | 421 | Scaffoldin | Involve in the cell adhesion, cell spreading and in the reorganization at the actin cytoskeleton. | Ovarian cancer, breast cancer, endometrial cancer, colorectal cancer, gastric cancer, lung cancer, prostate cancer, head and neck squamous cell cancer. | PET domain | https://www.protein |
| Tubby-related protein 1, Tubby-like protein 1 | TULP1, TUBL1 | 6 | 542 | Anchoring | Involve in protein trafficking through the connecting cilium into the outer segment of photoreceptor cells. | Retinal and cochlear degeneration, retinitis pigmentosa (RP), adult-onset obesity associated with insulin resistance. | | https://www.protein |
| Kelch-like protein 23 | KLHL23 | 2 | 558 | Bundling | Suppress F-actin, filopodium and lamellipodium formation. | Bladder urothelial carcinoma, hepatocellular carcinoma (HCC), pancreatic cancer, urothelial carcinoma. | | https://www.protein |
| Kelch-like ECH-associated protein 1, Cytosolic inhibitor of Nrf2 (INrf2), Kelch-like protein 19 | KEAP1, INRF2, KIAA0132, KLHL19 | 19 | 624 | | Stabilize F-actin cytoskeleton structures and inhibit focal adhesion turnover. | Lung adenocarcinoma, large cell carcinoma, prostate carcinoma. | Double glycine repeat domain | https://www.protein |

| | | | | | | | | |
|---|---|---|---|---|---|---|---|---|
| Ectoderm-neural cortex protein 1(ENC-1), Kelch-like protein 37, Nuclear matrix protein NRP/B, p53-induced gene 10 protein | ENC1, KLHL37, NRPB, PIG10 | 5 | 589 | | Require for adipocyte differentiation. | Hairy cell leukemia (HCL), urothelial carcinoma. | | https://www.protein |
| Protein S100-A4, Calvasculin, Metastasin, Placental calcium-binding protein, Protein Mts1, S100 calcium-binding protein A4 | S100A4, CAPL, MTS1 | 1 | 101 | Bundling, Scaffolding | Change in cell morphology, adhesion and migration. | Kidney fibrosis, liver fibrosis, pulmonary fibrosis and artery diseases, cardiac hypertrophy and fibrosis and rheumatoid arthritis, obesity, metastatic colorectal cancer, prostate cancer, brain tumors, breast, colon and lung carcinomas, Alzheimer’s (AD) and Parkinson’s diseases (PD), in cerebral ischemia, epilepsy, and schizophrenia, nervous system acute injuries. | | https://www.protein |

| | | | | | | | | |
|---|---|---|---|---|---|---|---|---|
| Protein S100-A6, Calcyclin, Growth factor-inducible protein 2A9, MLN 4, Prolactin receptor-associated protein, PRA, S100 calcium-binding protein A6 | S100A6, CACY | 1 | 90 | Monomer binding, F-actin binding | Regulate the cell cycle and morphology. | Pancreatic, gastric and prostate cancer, melanoma, non-small cell lung carcinoma, hepatocellular carcinoma, amyotrophic lateral sclerosis (ALS), Alzheimer’s disease, urinary bladder urothelial carcinoma, ovarian cancer, acute coronary syndrome and myocardial infarction. | | https://www.protein |
| Tripartite motif-containing protein 3, Brain-expressed RING finger protein, RING finger protein 22, RING finger protein 97 | TRIM3, BERP, RNF22, RNF97 | 11 | 744 | | Polyubiquitylates γ-actin to regulate synaptic γ-actin turnover and actin filament stability and thus form a transient inhibitory constraint on the expression of hippocampal synaptic plasticity. | Parkinson's disease, esophageal squamous cell carcinoma. | | https://www.protein |
| Septin-9, MLL septin-like fusion protein MSF-A (MLL septin-like fusion protein), Ovarian/Breast septin (Ov/Br septin), Septin D1 | SEPTIN9, KIAA0991, MSF, SEPT9 | 17 | 586 | Cross-linking | Maintain the integrity of growing and contracting actin filaments. | Colorectal cancer, breast cancer, hematological tumors, head and neck squamous cell carcinoma, ovarian cancer, lung cancer, gastric cancer. | Basic domain (B-domain) | https://www.protein |
| Uveal autoantigen with coiled-coil domains and ankyrin repeats protein, Beta-actin-binding protein, Beta cap73 | UACA | | | Capping | Cap barbed end. Act to spatially regulate the intracellular distribution of isoactins, facilitate forward protrusion formation. | | Shown with bovine protein | https://www.protein |

| | | | | | | | | |
|---|---|---|---|---|---|---|---|---|
| Harmonin, Antigen NY-CO-38/NY-CO-37, Autoimmune enteropathy-related antigen AIE-75, Protein PDZ-73, Renal carcinoma antigen NY-REN-3, Usher syndrome type-1C protein | USH1C, AIE75 | 11 | 552 | Bundling, Scaffolding | Involve in the adaptation of mechanoelectrical transduction by sensory hair cells. | Usher syndrome 1C (USH1C), Deafness, autosomal recessive, 18A (DFNB18A). | | https://www.protein |
| Heat shock protein beta-1, HspB1, 28 kDa heat shock protein, Estrogen-regulated 24 kDa protein, Heat shock 27 kDa protein (HSP27), Stress-responsive protein 27 (SRP27) | HSPB1, HSP27, HSP28 | 7 | 205 | Stabilization | A molecular chaperone involved in stress resistance and actin organization. | Charcot-Marie-Tooth disease 2F (CMT2F), Neuronopathy, distal hereditary motor, 2B (HMN2B). | | https://www.protein |
| Myelin basic protein, Myelin A1 protein, Myelin membrane encephalitogenic protein | MBP | 18 | 304 | Bundling, Polymerization | Create a cytosol to membrane signal caused by changes in interaction of the cytoskeleton with the membrane. | Rheumatoid arthritis, multiple sclerosis, demyelinating disease, brain cancer. | | https://www.protein |
| Neurocalcin-delta | NCALD | 8 | 193 | | Control clathrin-coated vesicle traffic. | Alzheimer's disease, spinal muscular atrophy (SMA), diabetic nephropathy. | | https://www.protein |
| Synapsin-1, Brain protein 4.1, Synapsin I | SYN1 | X | 705 | Nucleation, Bundling, Anchoring | Regulate neurotransmitter release by cross-linking synaptic vesicles to the actin cytoskeleton. | Epilepsy X-linked, with variable learning disabilities and behavior disorders (XELBD). | | https://www.protein |
| Synapsin-2, Synapsin II | SYN2 | 3 | 582 | Nucleation, Bundling | Cross-link synaptic vesicles and actin filaments in the nerve terminal. | Schizophrenia (SCZD). | | https://www.protein |
